# Information Dynamics of the Heart and Respiration Rates: a Novel Venue for Digital Phenotyping in Humans

**DOI:** 10.1101/2024.01.21.576502

**Authors:** Soheil Keshmiri, Sutashu Tomonaga, Haruo Mizutani, Kenji Doya

## Abstract

In recent decade, wearable digital devices have shown potentials for the discovery of novel biomarkers of humans’ physiology and behavior. Heart rate (HR) and respiration rate (RR) are most crucial bio-signals in humans’ digital phenotyping research. HR is a continuous and non-invasive proxy to autonomic nervous system and ample evidence pinpoints the critical role of respiratory modulation of cardiac function. In the present study, we recorded longitudinal (up to 6 days, 4.63 ***±*** 1.52) HR and RR of 89 freely-behaving human subjects (Female: 39, age 57.28 ***±*** 5.67, Male: 50, age 58.48 ***±*** 6.32) and analyzed their HR and RR dynamics using linear models and information theoretic measures. While the predictability by linear autoregressive (AR) showed correlation with subjects’ age, an information theoretic measure of predictability, active information storage (***AIS***), captured these correlations more clearly. Furthermore, analysis of the information flow between HR and RR by transfer entropy (i.e., ***HR → RR*** and ***RR → HR***) revealed that ***RR → HR*** is correlated with alcohol consumption and exercise habits. Thus we propose the AIS of HR and the transfer entropy ***RR → HR*** as two-dimensional biomarkers of cardiorespiratory physiology for digital phenotyping. The present findings provided evidence for the critical role of the respiratory modulation of HR, which was previously only studied in non-human animals.

## 1 Introduction

Humans’phenotype refers to observable characteristics and traits [1]. They range from individuals’ developmental processes [2, 3] to their behavior [4] and (electro)physiology [5, 6]. Recent decades have witnessed a substantial growth in human phenotyping [1]. This is due to its invaluable utility for diagnostics [7–10], gene-disease discovery [11, 12], and cohort analytics [13, 14]. It is apparent that these lines of research play a critical role in realization and advancement of the precision medicine [1, 6]. The advent and ubiquity of mobile technologies have substantially expedited the quest for the discovery of humans’ phenotypes [15, 16]. This paper explores human health phenotyping through longitudinal physiological monitoring by a wearable device and data analysis by dynamical systems and information theoretic approaches.

A rich body of research highlights the utility of wearable devices for digital health phenotyping [17–23]. For instance, Smets et al. [24] derived digital phenotypes from individuals’ five-day physiological and contextual measurements. They showed that their digital markers were able to predict their subjects’ depression, anxiety, and stress scores. Similarly, Jacobson and colleagues [25] found that actigraphy data was a reliable marker of changes in symptoms in patients with major depressive disorder and bipolar disorder across a two-week period. Straus and colleagues [20] utilized data from wrist-wearable devices. They identified the reduced variance in 24-hour activity as a marker of pain severity in traumatic stress patients. Using arm acceleration data, Katori et al. [21] found sleep phenotypes that associated with social jet lag, individuals’ chronotype, and insomnia. The use of mobile technologies for humans’ phenotyping becomes more relevant, given the common rhythms in their multi-day traits of use by individuals [22].

Heart rate (HR) and its variability (HRV) form another active frontier of research in humans’ phenotyping [23, 26–29]. Although not under the rubric of digital phenotyping, the search for biomarkers ^1^ of cardiac function has a long-lasting history [32–34, 34]. This is partly due to HR’s continuous and non-invasive proxy to autonomic nervous system whose central role in maintaining physiological homeostasis is well-established [33, 35]. It is also due to the fact that HR alone is an independent risk factor for cardiovascular-related mortality [36, 37].

A longitudinal study by Natarajan et al. [23] using wrist-worn tracking devices found that HRV decreased by aging [38, 39]. They further observed that increased physical activity may benefit cardiac function via improving HRV. Using wearable health-monitoring devices, Golbus and colleagues [29] found that older individuals (*>* 65 of age) had lower HRV. They also observed that HR varied by sex, age, race, and ethnicity and that it was higher in males than females. The relevance of HR and HRV for humans’ phenotyping is evident in their age- and gender-dependent manifestation of cardiac function [38, 40, 41]. They are important indicators of various age-related diseases and syndromes [42–47].

Taken together, these previous findings provide a wealth of evidence for the utility of wearable devices in digital phenotyping of humans’ cardiac function. They also underline the pivotal diagnostics/prognostics role that such biomarkers can play in monitoring individuals’ cardiac health [23, 38]. On the other hand, they fall short in providing adequate explanation for the extent of the relation among desperate markers of cardiac function [34]. Several studies attempted to address this issue through modeling strategies [35, 48–50]. However, their results were limited to parametric methodologies. As a result, they did not allow for a thorough investigation of the underlying nonlinear time-variant dynamics of cardiac function [43, 51, 52]. Additionally, these studies suffered from other shortcomings that included the imposition of broad assumptions and the lack of careful statistical analysis [35]. More importantly, ample findings pinpoint the critical role of cardiorespiratory mechanism in health [53– 60] and disease [61–63]. However, there is a paucity of research on digital phenotyping of the humans’ cardiorespiratory physiology.

In this study, we continuously monitor HR, RR and other physiological an behavioral signals by a wearable device for up to 6 days and seek useful biomarkers using state space reconstruction, linear prediction, and information theoretic frameworks.

## 2 Materials and Methods

### 2.1 Subjects

From an initial 130 healthy adults who participated in this experiment (Females = 60, Age: Mean (M) = 58.37, Standard Deviation (SD) = 5.93, Median (Mdn) = 58.50, Males = 70, Age: M = 58.89, SD = 6.33, Mdn = 58.50), 89 of them (Female: 39, Mean (M) = 57.28, Standard Deviation (SD) = 5.67, Male: 50, M = 58.48, SD = 6.32) who had at least 1 day (out of 6 consecutive days of experiment) of their Vital Patch RTM (https://vitalconnect.com) recordings (sampling rate: 0.25Hz, i.e., 1 data point every 4 seconds) available were included in the present study.

### 2.2 Recorded Physiology and Posture

Participants’ recorded data included their HR, RR, and posture (5 discrete categories based on accelerometer: laying down, leaning back, standing, walking, and running).

### 2.3 Daily Activities and Habits Questionnaire

Participants also filled in a “daily activities and habits” questionnaire that included their sleep/wake hours (Table 1), smoking (Table 2), alcohol consumption (Table 3), and exercise (Table 4) habits.

**Table 1:**
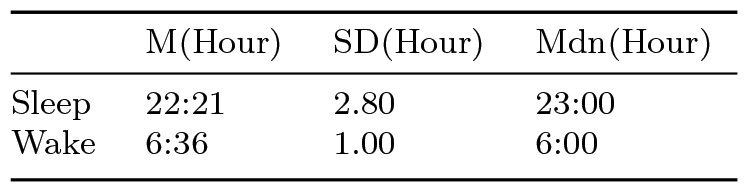
Participants’ Sleep/Wake.

|  | M(Hour) | SD(Hour) | Mdn(Hour) |
| --- | --- | --- | --- |
| Sleep | 22:21 | 2.80 | 23:00 |
| Wake | 6:36 | 1.00 | 6:00 |

**Table 2:**
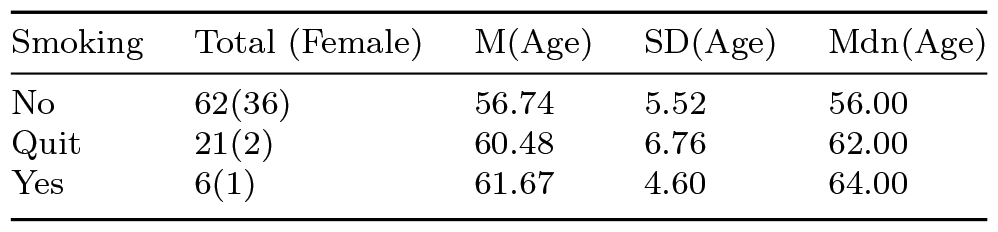
Participants’ Smoking Habit.

**Table 3:**
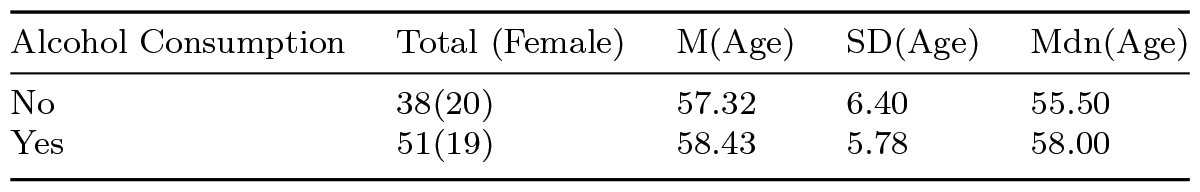
Participants’ Alcohol Consumption Habit.

**Table 4:** Participants’ Exercise Habit.

| Alcohol Consumption | Total (Female) | M(Age) | SD(Age) | Mdn(Age) |
| --- | --- | --- | --- | --- |
| No | 39(22) | 56.59 | 5.66 | 54.00 |
| Yes | 50(17) | 59.02 | 6.17 | 59.00 |

### 2.4 Experiment Procedure

Participants were carefully selected and thoroughly briefed about the study’s procedure. They were given sufficient time to understand the experiment and freely consent to their involvement.

After obtaining written informed consent from the participants, their background data was collected. It included gender, age, height, weight, medical history, medication and/or health supplements, allergy status, and skin condition susceptibility. In addition, we also collected the information about their smoking, exercise, and drinking habits. This information-gathering step carried out online with participants self-reporting all the required information.

We used VitalPatch RTM (https://vitalconnect.com) to record participants’ HR, RR, and posture at real-time (sampling rate: 0.25Hz, i.e., 1 data point every 4 seconds). We instructed the participants to wear the device throughout the experiment (i.e., 6 consecutive days). During this period, the participants carried on with their normal daily lives, routines, and activities.

### 2.5 Modeling of the Cardiorespiratory Dynamics

First, we applied a 1-minute non-overlapping moving average on the participants’ HR and RR time series, per participant, per day, per time series. This resulted in time series with length 60 *×* 24 *×* ⅅ= 1440 *×* ⅅ, per participant, per HR and RR. Here, 60 is the number of data points per hour and ⅅ ∈ [1, 2, 3, 4, 5, 6] denotes the number of days of experiment that were available from every individual. For instance, two individuals with whose number of days ⅅ = 1 and ⅅ = 3 would have HR and RR time series of length 1440 *×* 1 = 1440 and 1440 *×* 3 = 4320, respectively.

#### 2.5.1 State space reconstruction

We reconstructed the individuals’ HR state-space, one per individual, using their HR time series of length 1440 *×* ⅅ. We achieved this by fixing all individuals’ HR state-space embedding dimension to 3 and estimating their respective best delay embedding *τ* using Rosenstein et al. algorithm [64]. We then determined the *μ*_*τ*_ by bootstrapping (10,000 repetitions) the individuals’ *τ* values at 95% confidence interval (CI) (Figure **??**). Last, we set every individuals’ *τ* = *μ*_*τ*_ and reconstructed their respective final HR state-space. This resulted in HR state-space 2-dimensional matrices, one per individual, of size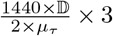where 3 is the HR state-space embedding dimension. As an example, if *μ*_*τ*_ = 128 (i.e., the estimated *μ*_*τ*_ in the present study, Section 3.4 and Supplementary Materials 1 (SM1)), then an individual with whose ⅅ = 1 would have an HR state-space matrix of size 1184 *×* 3.

#### 2.5.2 Linear modeling

We quantified the HR state-space dynamics using both model-based and model-free approaches. In the case of model-based, we fitted an autoregressive (AR) model to the individuals’ mean-centered HR state-space (one AR per participant):

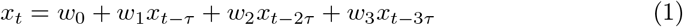

where *t* is the current time step (in minutes), *τ* is the HR state-space delay embedding (set to *μ*_*τ*_ for all participants), and *w*_*i*_, *i* = 1, … 3 are the AR parameters. Given that the participants’ HR state-space were mean-centered, *w*_0_ *≈* 0 (*≈* is exact i.e., *w*_0_ = 0, in the case of perfect fit). Therefore we excluded *w*_0_ from further analyses.

#### 2.5.3 Information theoretic measures

We quantified the participants’ HR state-space variability via computing its entropy *H* (multivariate, given its dimension = 3) [65, 66]. In this context, *H* signifies the degree of dispersion in individuals’ HR state-space (i.e., its instability): *H* is the sum of all positive Lyapunov exponents whose magnitude reflects the rate of information loss (and instability) over time [67].

Furthermore, we computed active information storage (*AIS*) of the participants’ HR state-space. *AIS* quantifies the information in HR state-space’s past that actively contributed to computing its current state [68]:

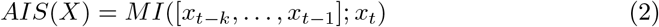

where *MI*(*X*; *Y*) = *H*(*X*) + *H*(*Y*) *− H*(*X, Y*) is the mutual information (*MI*) between *X* and *Y* (here, the present and the past of HR state-space) and *t* denotes time (in minutes). In essence, *AIS* signifies the component of the HR state-space that is directly in use in the computation of its next state.

Additionally, we determined the presence of long-range correlation in the participants’ HR state-space (and therefore the deviation from the independence among its components) by computing HR state-space integration (*I*, a.k.a multi-information) [69]:

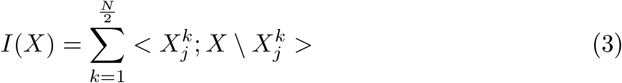

where *j* is the number of bipartitions of *X* composed of *k* components. Since individuals’ HR state-space were reconstructed in 3 dimensions, there were 6 such bipartitions: the univariate cases *x*_*t*_, *x*_*t−τ*_, *x*_*t−*2*τ*_, and the bivariate cases [*x*_*t*_, *x*_*t−τ*_], [*x*_*t*_, *x*_*t−*2*τ*_], and [*x*_*t−τ*_, *x*_*t−*2*τ*_].

#### 2.5.4 Transfer entropy

We captured the statistical precedence between HR and RR to quantify the functional effect of each process on the other. For this purpose, we used HR and RR original univariate time series (both mean-centered as in the case of HR state-space *H, AIS*, and *I*) to compute their transfer entropy (TE) [70]:

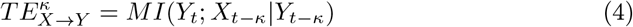

where *κ* is the time lag, *MI*(*X*; *Y* |*Z*) = *H*(*X*|*Z*) *− H*(*X*|*Y, Z*) is the *MI* between *X* and *Y* conditioned on *Z*, and *H*(*X*|*Z*) computes the entropy (*H*) of *X* conditioned on *Z*. Throughout the manuscript, we used *HR → RR* and *RR → HR* to refer to 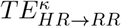 and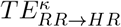, respectively.

Similar to the case of HR state-space delay embedding *τ*, we first computed *κ*, per individual, such that it maximized *HR → RR* (SM1). This was achieved through a brute-force strategy [71] in which *κ* was incremented (in a 1-minute increment steps, i.e., 1-data-point at a time) within a given interval. We used the same interval as in the case of HR state-space *τ* (SM1). We retained the value of *κ* that resulted in largest *TE*, per individual. We then bootstrapped (10,000 repetitions) these individuals’ *κ* values, thereby estimating the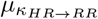(SM1). We set every individuals’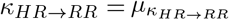and recomputed their final *HR → RR*. We repeated this procedure for the case of *κ*_*RR→HR*_ (SM1) to recompute their final *RR → HR*. We used JIDT [72] (version v1.5 for Python) implementation of *H* and *AIS* that are based on Kraskov-Stoegbauer-Grassberger (KSG) algorithm [73], its implementation of *I* that is based on Tononi-Sporns-Edelman (TSE) algorithm [69], and its KSG-based implementation of Schreiber [70] TE computation. We carried out all the computations and analyses in Python 3.10.4.

#### 2.5.5 AR, HR, and HR – RR Interplay

First, we examined the extent at which HR state-space dynamics was captured within the model-based and the model-free planes (model-based (AR) planes: *w*_1_ – *w*_2_, *w*_1_ – *w*_3_, *w*_2_ – *w*_3_, and model-free plane: *RR → HR* – *HR → RR*). Specifically, we investigated (1) the degree at which the respective axes of each of these planes (e.g., *RR → HR* and *HR → RR*) correlated and (2) the extent by which such correlations corresponded to AR’s coefficient of determination (i.e., its accuracy *R*^2^), on the one hand, and HR state-space information dynamics, *AIS* and *I*, on the other hand (See accompanying supplementary materials SM3 for these observations based on HR’s moments (i.e., mean, variance ^2^, skewness, kurtosis), Poincaré SD1 and SD2 [74], and power (low frequency, high frequency, and total) that are frequently used in the literature [34]). We corrected the correlation (Spearman) results using Bonferroni-correction (corrected p = 0.05*/*89 *≈* 6.0*e*^*−*04^, given 89 individuals in the present study). Consequently, we considered an observed correlation significant if and only if its corresponding p *<* 6.0*e*^*−*04^. For clarity and ease of comparison, we reported the uncorrected p-values throughout the manuscript. We reported the AR-related results in SM1.

#### 2.5.6 HR Information Dynamics, HR – RR Interplay, and Individuals’ Biological Phenotypes and Habits

We conducted the permutation tests based on the participants’ (biological) phenotypic information (i.e., age and gender). We also repeated these tests on their responses to “daily activities and habits” questionnaire (i.e., their alcohol consumption, exercise, and smoking habits). While examining the case of “smoking habit,” we divided the individuals into two groups that corresponded to “non-smokers” and “smokers or those who quit smoking.” This choice was due to the limited number of smokers in our sample (6 smokers in total, Table 2). We reported these smoking-related results in SM1.

To better examine the potential effects of individuals’ daily habits on their HR state-space information dynamics and HR – RR interplay, we further applied our permutation tests on the following four scenarios.

1. individuals who consumed alcohol (N = 17, females: 10, Age = 56.06 *±* 5.05) versus those who did not (N = 12, Females = 10, Age = 53.75 *±* 3.29) but otherwise none of them smoked (or quit smoking) or exercised.
2. individuals who exercised (N = 17, females = 9, Age = 57.94 *±* 6.30) versus those who did not (N = 12, females = 10 Age = 53.75 *±* 3.29) but otherwise none of them consumed alcohol or smoked (or quit smoking).
3. individuals who exercised but differed on their habit of alcohol consumption: those who did not consume alcohol (i.e., same as the case “2” above) versus those who did (N = 16, Females = 7, Age = 58.44 *±* 5.37) but otherwise none of them smoked (or quit smoking).
4. individuals who consumed alcohol but differed in their exercise habit: those who did not exercise (i.e., same as the case “1” above) versus those who exercised (N = 16, Females = 7, Age = 58.44 *±* 5.37) but otherwise none of them smoked (or quit smoking).

Considering the smoking habit, number of participants did not allow us to include it in this further analysis steps: there were only 4 participants who smoked (or quit smoking) but did not consume alcohol or exercised (females = 1, Age = 58.50 *±* 7.53), 2 participants who quit smoking but did not consume alcohol or exercise (females = 1, Age = 58.00 *±* 8.00), and 5 participants who smoked and exercised but did not drink (all males, Age = 62.8 *±* 6.43).

We reported the effect of age and gender on *AIS* and *I* and the effect of weekly alcohol consumption on *RR → HR* and *H* in the main manuscript. We included the results on the correlation between HR and RR (overall, sleep, and wake) along with the age-related differences in the participants’ exercise habit, the gender-related differences in their body-mass-index (BMI) and HR, the effect of weekly alcohol consumption on participants’ BMI and their HR (afternoon period, 12:00 – 18:00 PM), and the effect of smoking on individuals’ BMI and their early morning HR (6:00 – 7:00 AM) in SM2.

#### 2.5.7 Alcohol Consumption, Exercise, and *RR → HR*: Group-Level Analyses

We further verified the effects of the combination of the alcohol – exercise on the participants’ *RR → HR* as follows. First, we divided the participants to four groups, namely, (1) those who exercised but did not consume alcohol (2) those who neither exercised nor consumed alcohol (3) those who did not exercise but consumed alcohol (4) those who exercised as well as consumed alcohol. We then carried out two additional analyses: a two-way ANOVA and a non-parametric Kruskal-Wallis test of significant differences at group levels.

In the case of two-way ANOVA, we followed up its results by applying posthoc pairwise two-sample Welch test (equivalent of two-sample t-test for unequal variances) between these four groups. We corrected the results of both ANOVA and posthoc Welch test using false discovery rate (FDR).

In the case of Kruskall-Wallis test, we used pairwise non-parametric Wilcoxon ranksum as follow-up posthoc tests. Similar to the case of parametric tests (i.e., ANOVA and Welch tests), we corrected both Kruskal-Wallis and posthoc Wilcoxon rank-sum tests using FDR.

#### 2.5.8 Reported Effect Sizes

To quantify the strength of the observed significant differences in the permutation tests as well as parametric two-sample Welch test, we reported the Hedges “g” effect size [75]:

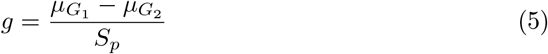

where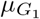and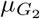are the two groups’ (i.e., *G*_1_ and *G*_2_ in equation (5)) mean and *S*_*p*_ represents their pooled standard deviation:

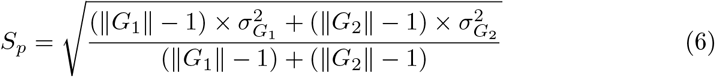

with ∥*G*_1_∥ and ∥*G*_2_∥ representing the *G*_1_’s and *G*_2_’s sample size and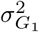 and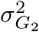 are their respective variance. Hedges effect size “g” is interpreted as small, medium, or large for *g* = 0.2, 0.5, or 0.8, respectively [76, 77]. Here *G*_1_ and *G*_2_ stand for two arbitrary groups being compared.

For the parametric ANOVA and non-parametric Kruskal-Wallis, we reported *η*^2^ effect-size. This effect is considered small, medium, or large for *η*^2^ = 0.02, 0.13, and 0.26, respectively [78, 79].

For non-parametric two-sample Wilcoxon rank -sum test, we reported the effect-size 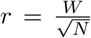[80] where *W* and *N* refer to Wilcoxon rank-sum test’s statistics and total sample size, respectively. *r* is considered small, SSSmedium, or large if *r ≤* 0.3, 0.3 *< r <* 0.5, and *r ≥* 0.5 [81].

We reported the two-way ANOVA and its associated posthoc pairwise two-sample Welch tests in the main manuscript. We presented the results associated with non-parametric Kruskal-Wallis and posthoc pairwise two-sample Wilcoxon rank-sum tests in SM1.

## 3 Results

### 3.1 Overview of HR and RR Data

Figure 1 shows the overall distribution of subjects’ HR and RR within the mean (*μ*) – SD (*σ*) plane.

**Fig. 1:**
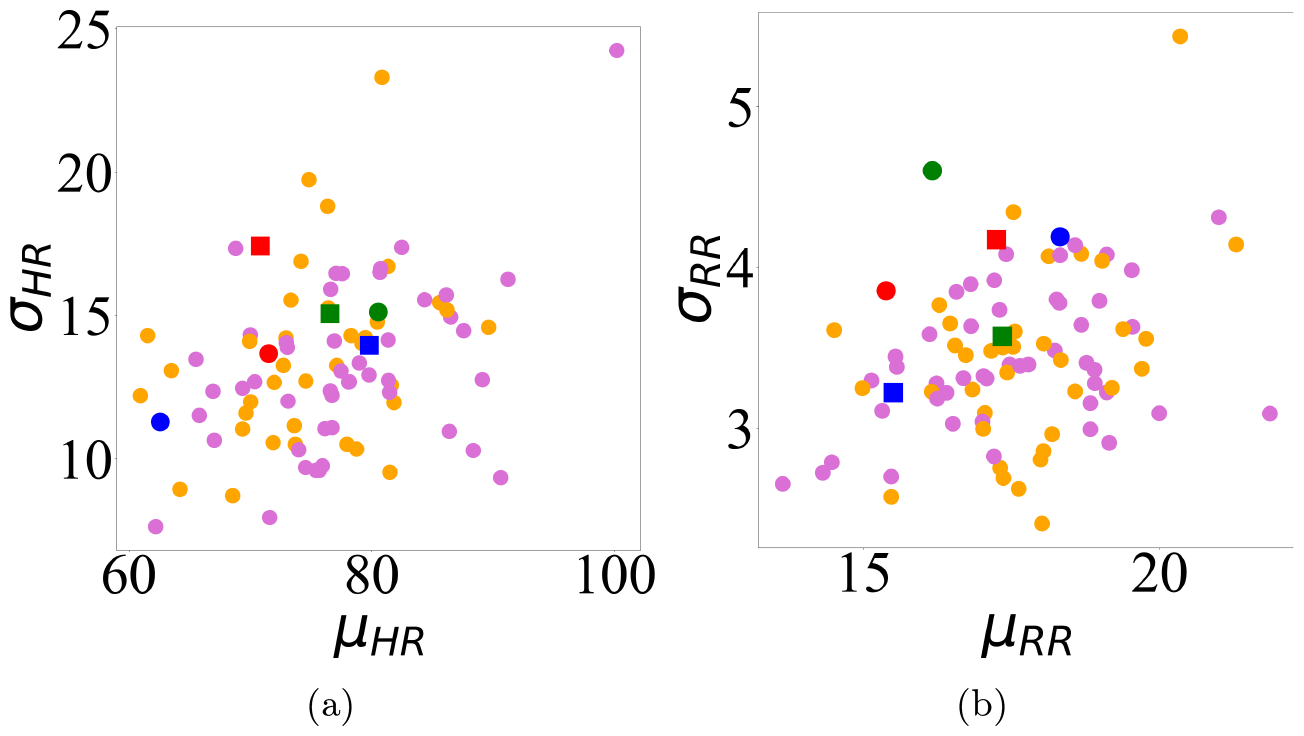
Mean (*μ*) – standard deviation (*σ*) scatter plots for (a) HR (b) RR. In these subplots, females’ and males’ data are colored in orange and pink, respectively. Markers in red, green, and blue highlight female (slightly larger circles) and male (square) participants in Figure 2.

In Figure 2, we show sample HR and RR time series for the participants whose respective average HR corresponded to the overall HR percentiles i.e., 25.00% (in red) 50.00% (in green) and 75.00% (in blue). In each subplot, the grand averages are shown in black (HR – Females: M = 67.2652, SD = 5.947, Males: M = 63.6022, SD = 8.2032; RR – M = 16.2507 SD = 2.2947, Males: M = 14.1811 SD = 0.5047). See SM1 for all participants’ HR and RR time series. SM1 also includes sample cardiorespiratory trajectories (i.e., HR – RR plane) of four randomly selected female and male individuals with 1 through 6 days of experiment (not necessarily the same participants). Section 2.1 summarizes the HR and RR descriptive statistics for females, males, and the overall sample (i.e., all 89 individuals combined).

**Fig. 2:**
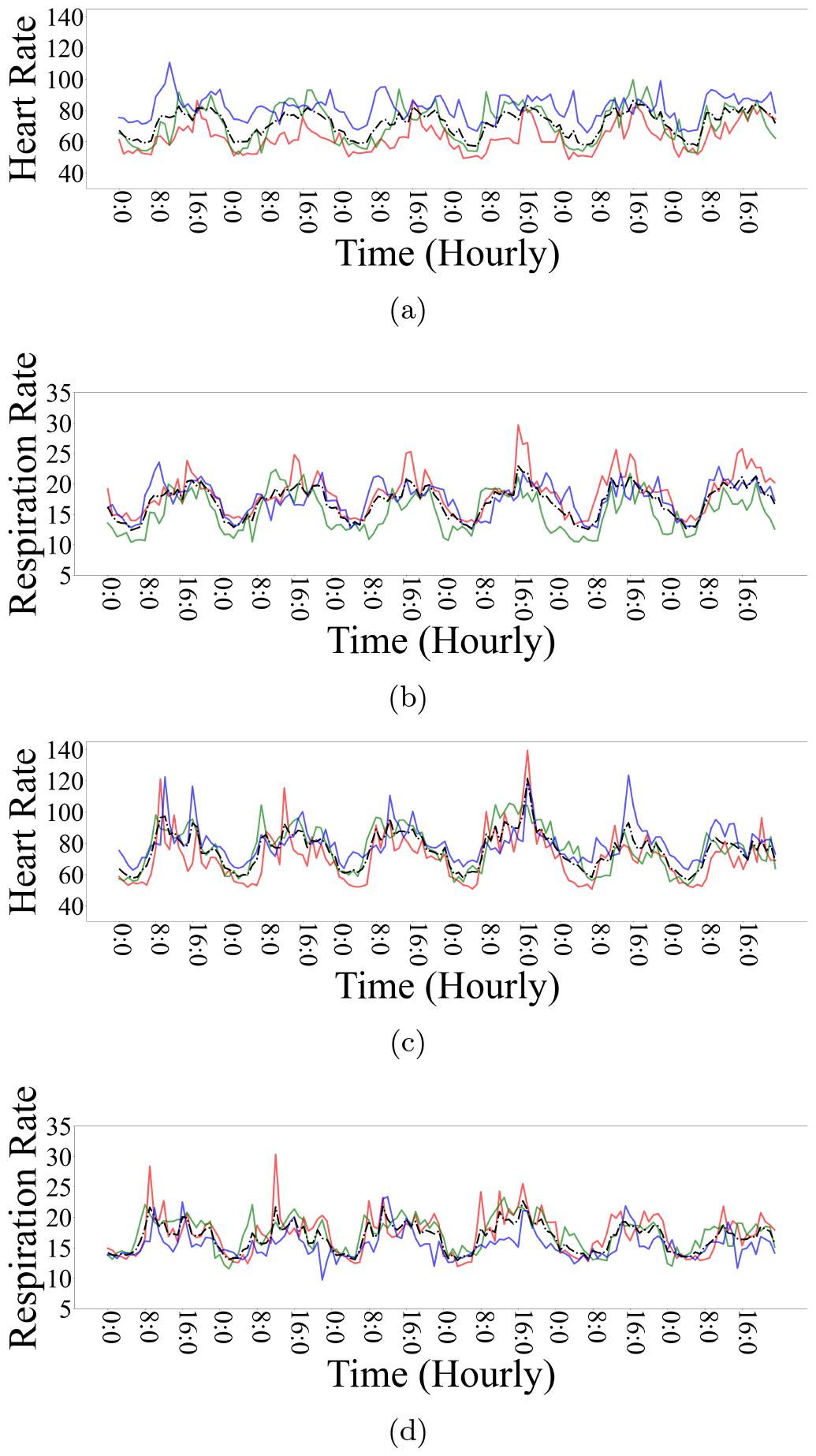
Three representative samples of female (a) HR (b) RR and male (c) HR (d) RR time series whose respective mean HR corresponded to percentile HR i.e., 25.00% (in red), 50.00% (in green), and 75.00% (in blue). In these subplots, HR and RR time series, per participant, are averaged in a 1-hour non-overlapping interval. Dotted black line represents grand average of time series within a subplot. See SM1 for all participants’ HR and RR time series.

Figure 3 depicts the grand averages (in 24-hour time span) of the participants’ HR (Figure 3a, all participants’ M = 76.54, SD = 15.67, Minimum (Min) = 32.33, Maximum (Max) = 197.33; Females: M = 74.84, SD = 15.62, Mdn = 73.33, CI_95%_ = [49.33, 109.47], Min = 34.93, Max = 164.00; Males: M = 77.82, SD = 15.58, Mdn = 76.13, CI_95%_ = [53.33, 112.73], Min = 32.33, Max = 197.33) and RR (Figure 3b, all participants’ M = 17.52, SD = 3.86, Min = 4.07, Max = 42.0; Females: M = 17.64, SD = 3.89, Mdn = 17.27, CI_95%_ = [11.20, 26.47], Min = 4.07, Max = 42.00; Males: M = 17.42, SD = 3.83, Mdn = 17.07, CI_95%_ = [11.07, 25.87], Min = 4.13, Max = 41.93).

**Fig. 3:**
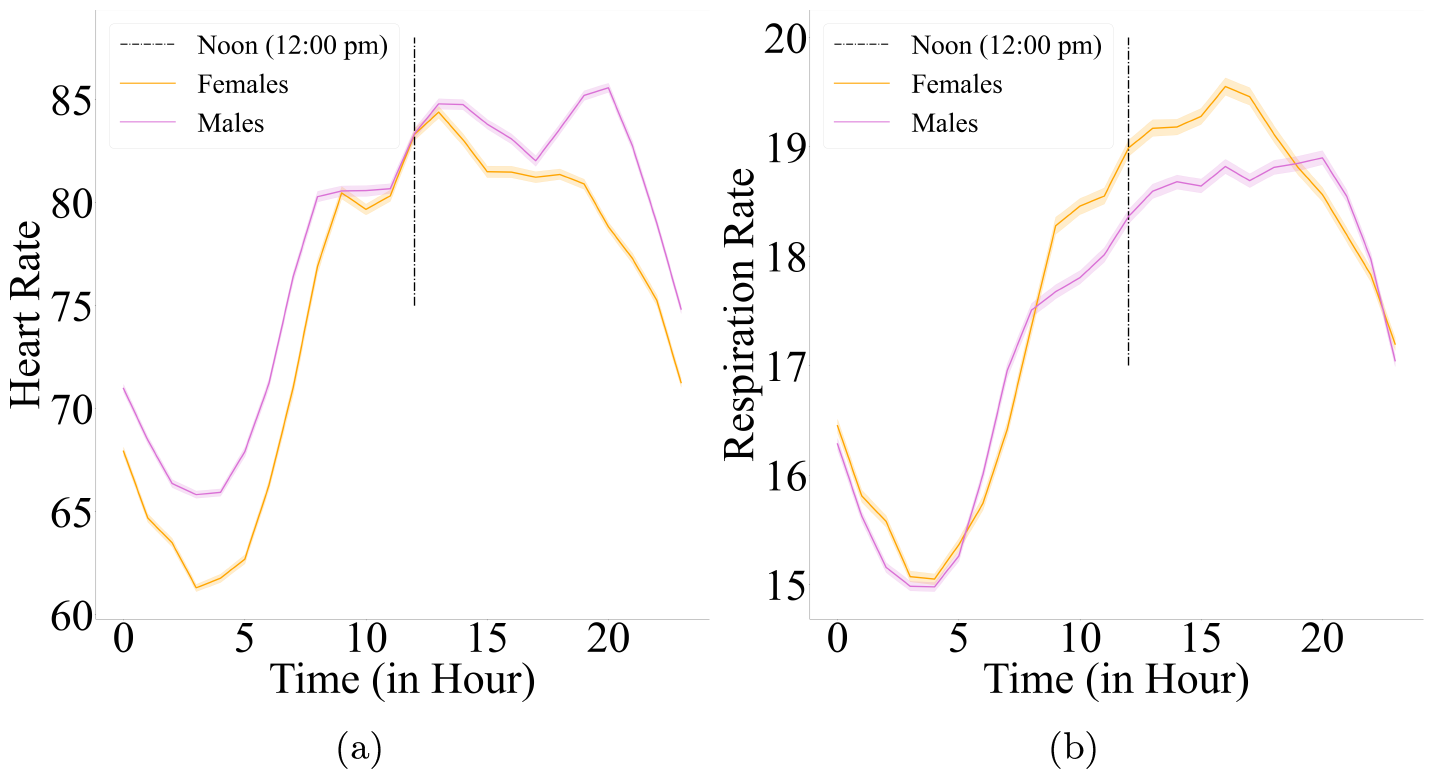
Grand averages (24-hour time span) of (a) HR (b) RR daily trajectories for female (in orange) and male (in pink) participants. In these subplots, black dash-line marks noon (i.e., 12:00 pm).

### 3.3 HR – RR Correlation

Figure 4 shows the HR – RR scatter plots for the same female and male participants in Figure 2. In these subplots, data points are color-coded as per their respective time-of-day (in an hourly basis i.e., Figure 4c).

**Fig. 4:**
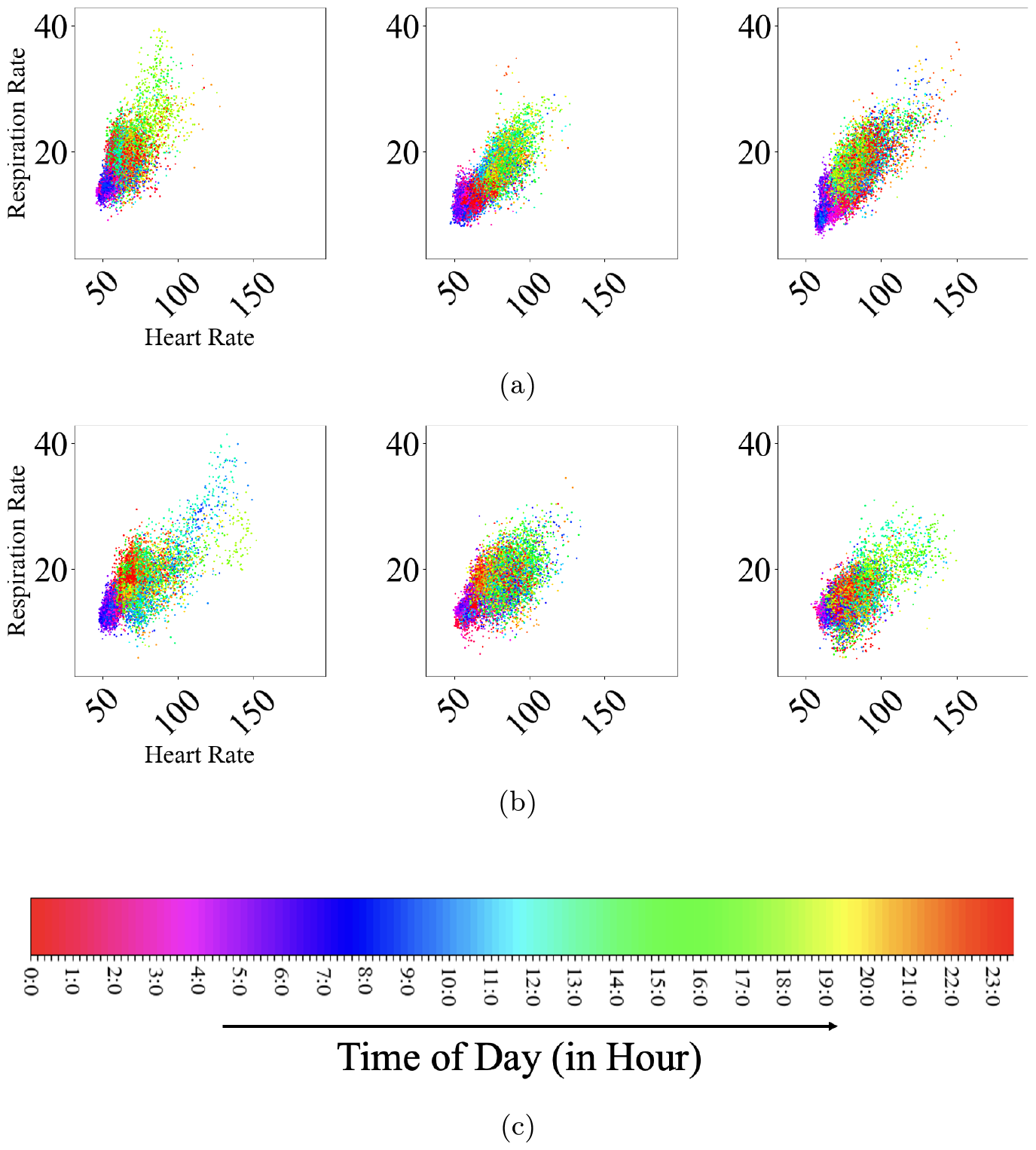
HR – RR scatter plots for the same example of female and male participants in Figure 2. (a) Females (b) Males participants. Data points correspond to 1-minute non-overlapping moving average on HR and RR time series. They are color-coded as per time-of-day (in an hourly basis) according to (c). See SM1 for plots of all participants.

Within this sub-sample (i.e., Figure 2), the average HR – RR correlation showed a stronger HR – RR correlation among females (Spearman’s rank correlation coefficient r = 0.7639, SD = 0.0583, Mdn = 0.7766, CI_95%_ = [0.6915, 0.8255], Min = 0.6870, Max = 0.8281) than males (Spearman’s rank correlation coefficient r = 0.6257, SD = 0.0546, Mdn = 0.6272, CI_95%_ = [0.5616, 0.6886], Min = 0.5581, Max = 0.6918). However, this difference disappeared after taking all female and male participants into account (Spearman’s rank correlation coefficient, females: M = 0.5713, SD = 0.1755, Mdn = 0.6052, CI_95%_ = [0.2228, 0.8078], Min = 0.0927, Max = 0.8281, males: M = 0.5573, SD = 0.1312, Mdn = 0.5522, CI_95%_ = [0.2676, 0.7503], Min = 0.2344, Max = 0.7694). See SM1 for all participants’ HR – RR scatter plots.

### 3.3 HR – RR Cross-Correlation

The combined females and males cross correlations for their days-averaged and within their *±*30 minutes interval are shown in Figures 5a (M = -1.65e^*−*18^, SD = 8.06e^*−*18^, Mdn = 0.0, CI_95%_ = [-1.88e^*−*17^, 9.87e^*−*18^]) and 5b (M = 0.8805, SD = 0.0865, Mdn = 0.9015, CI_95%_ = [0.5836, 0.9653], Min = 0.5366, Max = 0.9790). These subplots indicate a rather long delay lag (i.e., *≈* 1000 minutes i.e., *≡* 16.67 hours) between HR and RR.

**Fig. 5:**
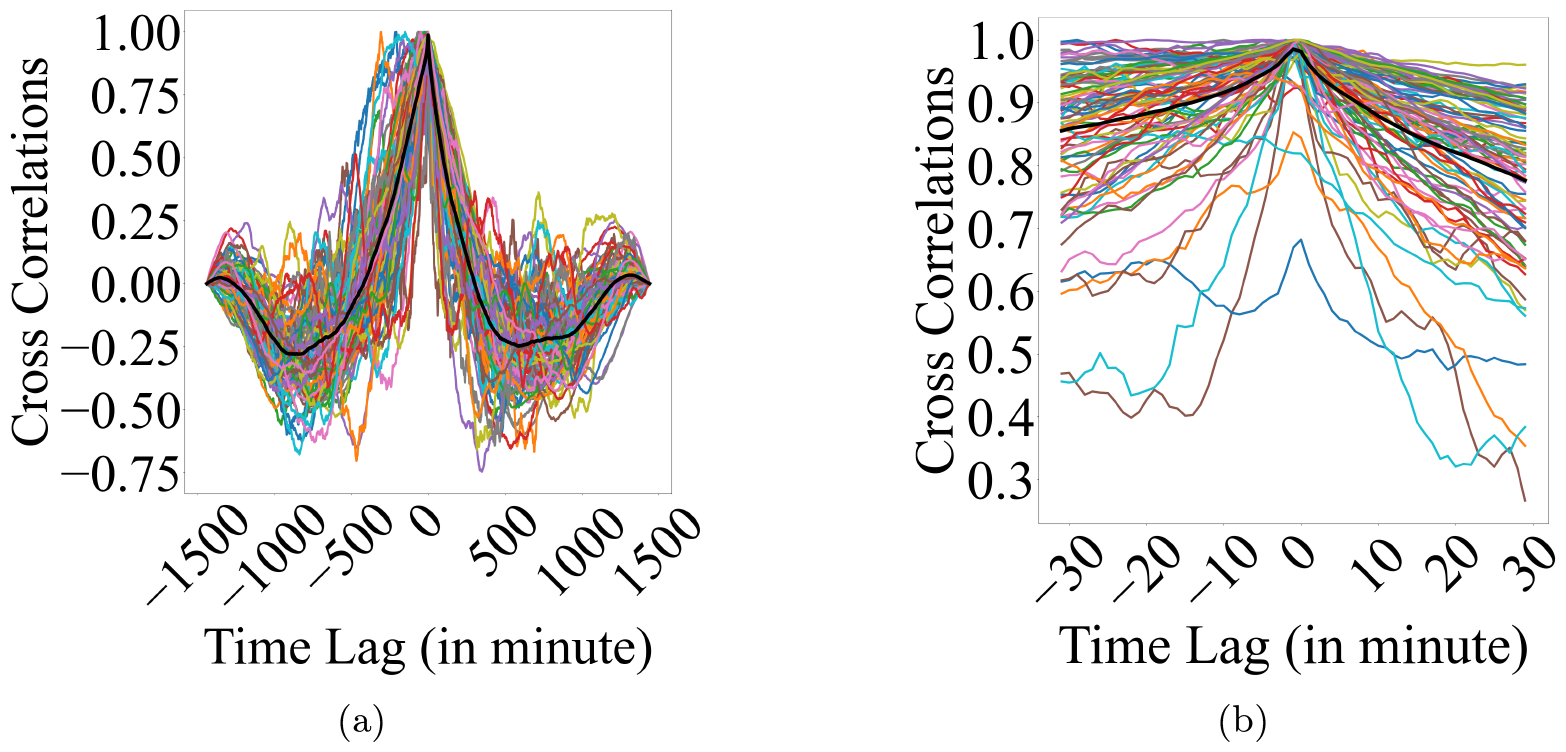
HR – RR days averaged cross-correlation of combined females and males for their (a) days-averaged (b) within *±*30 minutes window. In these subplots, HR and RR are mean-centered at individual level. Black tick line depicts the grand average of their corresponding cross-correlation plots in each subplot.

### 3.4 HR State-Space Delay Embedding

We derived optimal delay time *τ* for state space embedding by the principle of reconstruction expansion using Rosenstein et al. algorithm [64]. Figure 6 shows the distribution of the participants’ optimal HR state-space embedding delay time *τ* (M = 128.1236, SD = 40.5806, Mdn = 135.00, CI_95%_ = [35.9636, 45.9617], Min = 28.00, Max = 179.00).

**Fig. 6:**
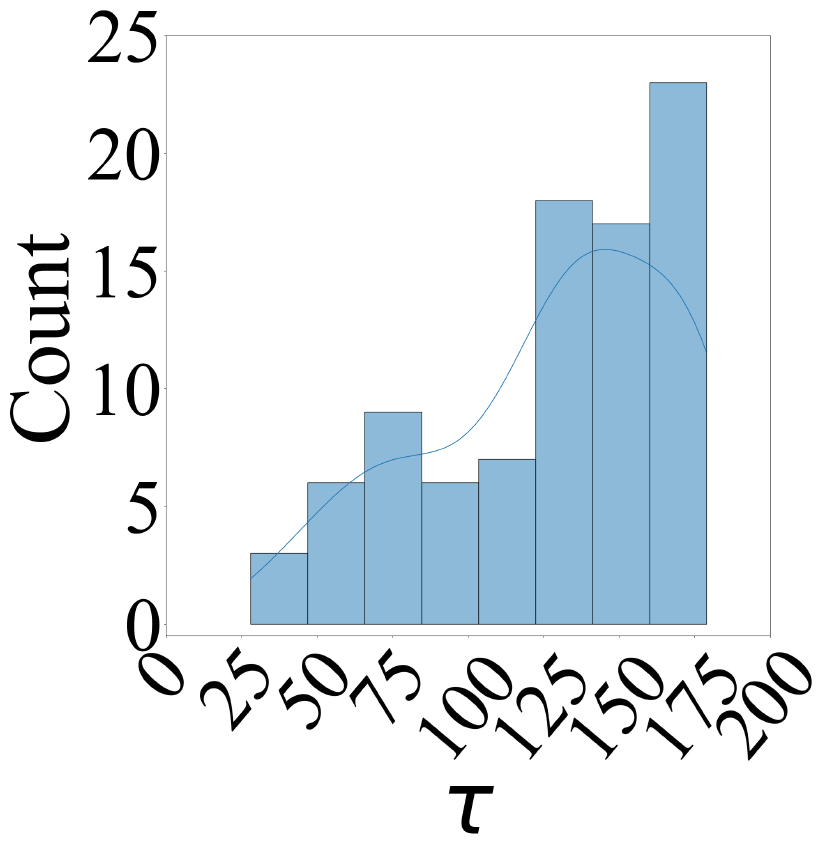
Distribution of individuals’ HR state-space delay embedding *τ*. While calculating individuals’ optimal *τ*, we performed a brute-force search over the range *τ ∈* [1, …, 240] (i.e., 1 through 240 minutes) using Rosenstein et al. algorithm [64]. Bootstrapping participants’ respective *τ* (Figure **??**) yielded *μ*_*τ*_ = 128 (i.e., 2 hours and 8 minutes). We set every participants’ *τ* = *μ*_*τ*_ to reconstruct their final HR state-space.

In SM1, we show the result of the bootstrap (10,000 repetitions) estimate of *μ*_*τ*_ at 95% CI (M = 128.1602, Mdn = 128.2472, CI_95%_ = [119.6739, 136.2921]). We used this *μ*_*τ*_ = 128 (i.e., 2 hours and 8 minutes) as fixed *τ* value to reconstruct all participants’ final HR state-space (i.e., *τ* = *μ*_*τ*_, for all participants).

The *μ*_*τ*_ = 128 indicated that, on average, HR state-space corresponded to the aggregate of heart rate values that exhibited long-range dependencies (i.e., *>* 2 hours). This is due to the fact (1) that every HR state is defined by three HR values that constituted its coordinates at that state within its overall state-space (2) that these values must, in principle, uniquely and smoothly determine HR states over time [82] and (3) that *τ* plays a crucial role in satisfying these uniqueness and smoothness requirements by quantifying the proper expansion of HR time series from the identity line of its embedding space [64]. Figures 7a and 7b depict the state-space for the same female and male participants whose HR and RR time series are depicted in Figure 2 using this *μ*_*τ*_ = 128. See SM1 for all participants’ state-space plots.

**Fig. 7:**
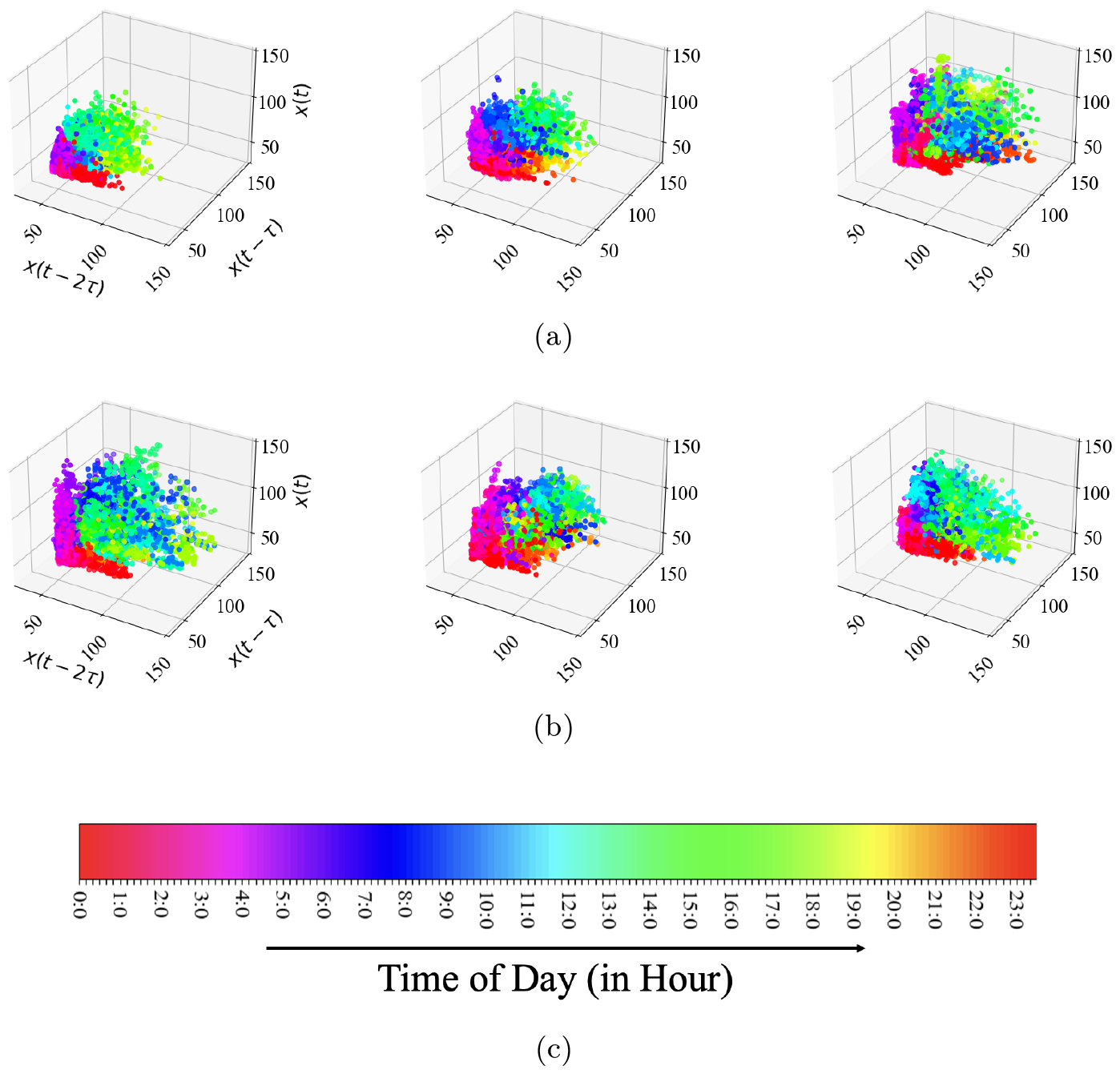
HR state-space for same sample of (a) female and (b) male participants in Figure 2, using delay embedding *μ*_*τ*_ = 128 i.e., 2 hours 8 minutes. Data points are color-coded as per time-of-day (in an hourly basis) according to (c). See SM1 for plots of all participants’ state-space plots.

### 3.5 HR AR Accuracy *R*^2^

We found no correlation between *μ*_*HR*_ and AR accuracy *R*^2^ (Figure 8a, r = 0.1387, p = 1.95e^*−*01^). Considering the AR accuracy *R*^2^ within *μ*_*HR*_ – *σ*_*HR*_ plane (Figure 8b), we observed no correspondence between *R*^2^ and the HR’s first and second moments.

**Fig. 8:**
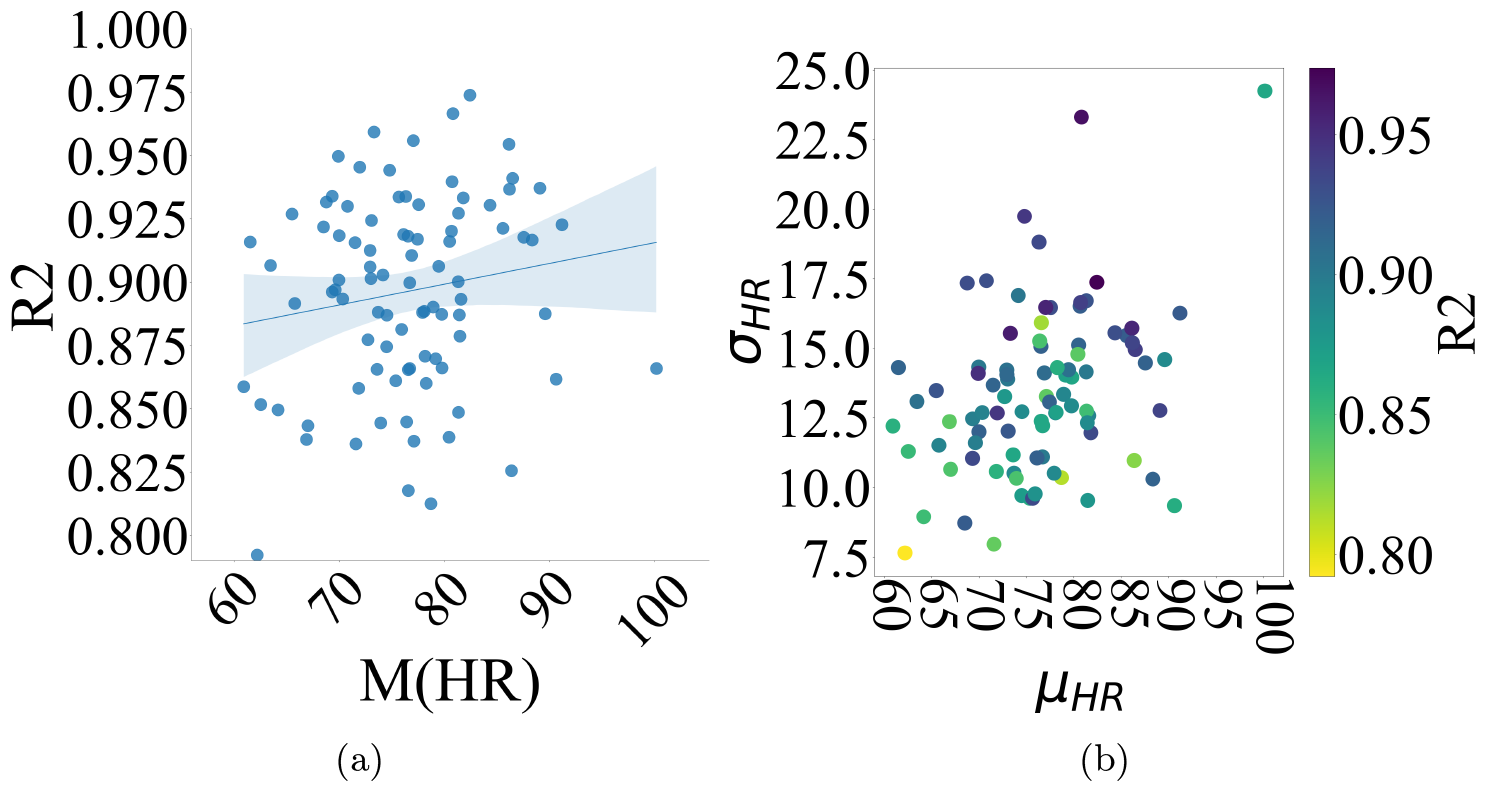
(a) Average HR versus AR accuracy *R*^2^. (b) AR accuracy *R*^2^ within *μ*_*HR*_ – *σ*_*HR*_ plane. We observed no discernible correspondence between *R*^2^ and HR’s first and second moments.

#### 3.5.1 TE Time Lags *κ*_*HR→RR*_ and *κ*_*RR→HR*_

Figure 9 plots the *κ* values for the range *κ ∈* [1, …, 15] minutes (SM1, i.e., up to the bootstrapped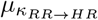). Whereas *κ*_*HR→RR*_ exhibited (Figure 9a) a marked maximum at *μ*_*κ*_ = 6 minutes, the evolution of *κ*_*RR→HR*_ was gradual (Figure 9b), attaining its maximum at *μ*_*κ*_ = 15 minutes. This difference in time lag between *HR → RR* and *RR → HR* resonated with their respective sympathetic and parasympathetic regulatory axes.

**Fig. 9:**
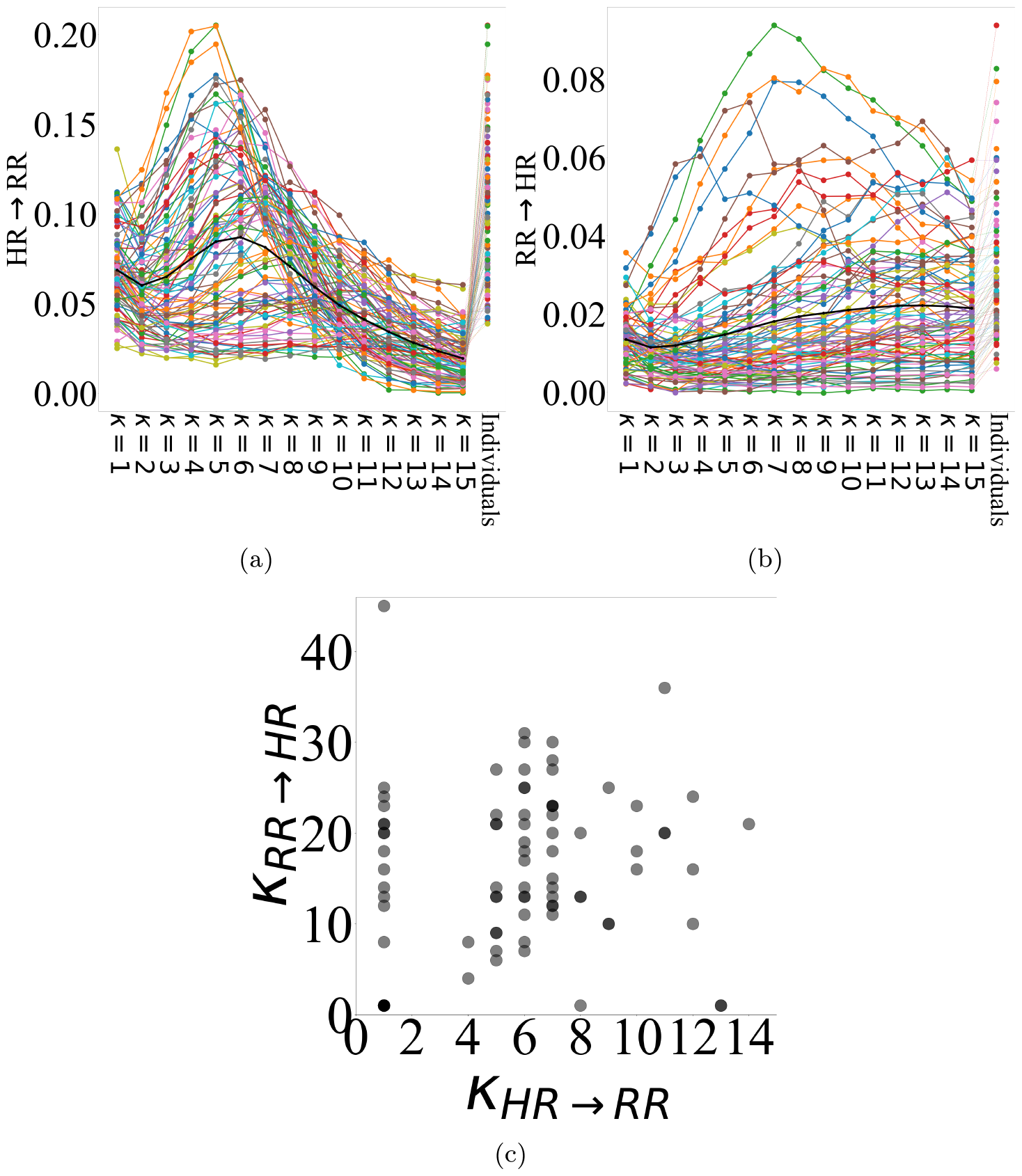
*κ* values for range *κ ∈* [1, …, 15] minutes (i.e., up to bootstrapped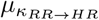). (a) *HR → RR* (b) *RR → HR*. Whereas *κ*_*HR→RR*_ exhibited a marked maximum at *κ* = 6, evolution of its value in case of *κ*_*RR→HR*_ was gradual, attaining its maximum at *κ* = 15. In these subplots, “Individuals” in x-axis refers to participant-specific *κ*_*HR→RR*_ and *κ*_*RR→HR*_. (c) *κ*_*HR→RR*_ – *κ*_*RR→HR*_ scatter plot. Similar to Figure 1, females’ and males’ data are colored in orange and pink, respectively. Markers in red, green, and blue highlight female (slightly larger circles) and male (square) participants in Figure 2. Weak correlation between *κ*_*HR→RR*_ and *κ*_*RR→HR*_ did not pass Bonferroni-correction (r = 0.2537, p = 1.64e^*−*02^). See SM1 for results of individuals’ *κ*_*HR→RR*_ and *κ*_*RR→HR*_ and their bootstrapping.

In these subplots, “Individuals” in the x-axis refers to the participant-specific *κ*_*HR→RR*_ and *κ*_*RR→HR*_. Although we observed a weak correlation between *κ*_*HR→RR*_ and *κ*_*RR→HR*_, this correlation did not pass the Bonferroni-correction (Figure 9c, r = 0.2537, p = 1.64e^*−*02^).

### 3.6 HR AR Accuracy and State-Space Information Dynamics

AR accuracy *R*^2^ showed strong correlations with HR state-space *AIS* (Figure 10b, r = 0.8396, p = 8.81e^*−*25^) and *I* (Figure 10c, r = 0.6884, p = 8.96e^*−*14^). On the other hand, *R*^2^’s substantially weaker correlation with *H* (Figure 10a, r = 0.2955, p = 4.93e^*−*03^) did not survive the Bonferroni-correction.

**Fig. 10:**
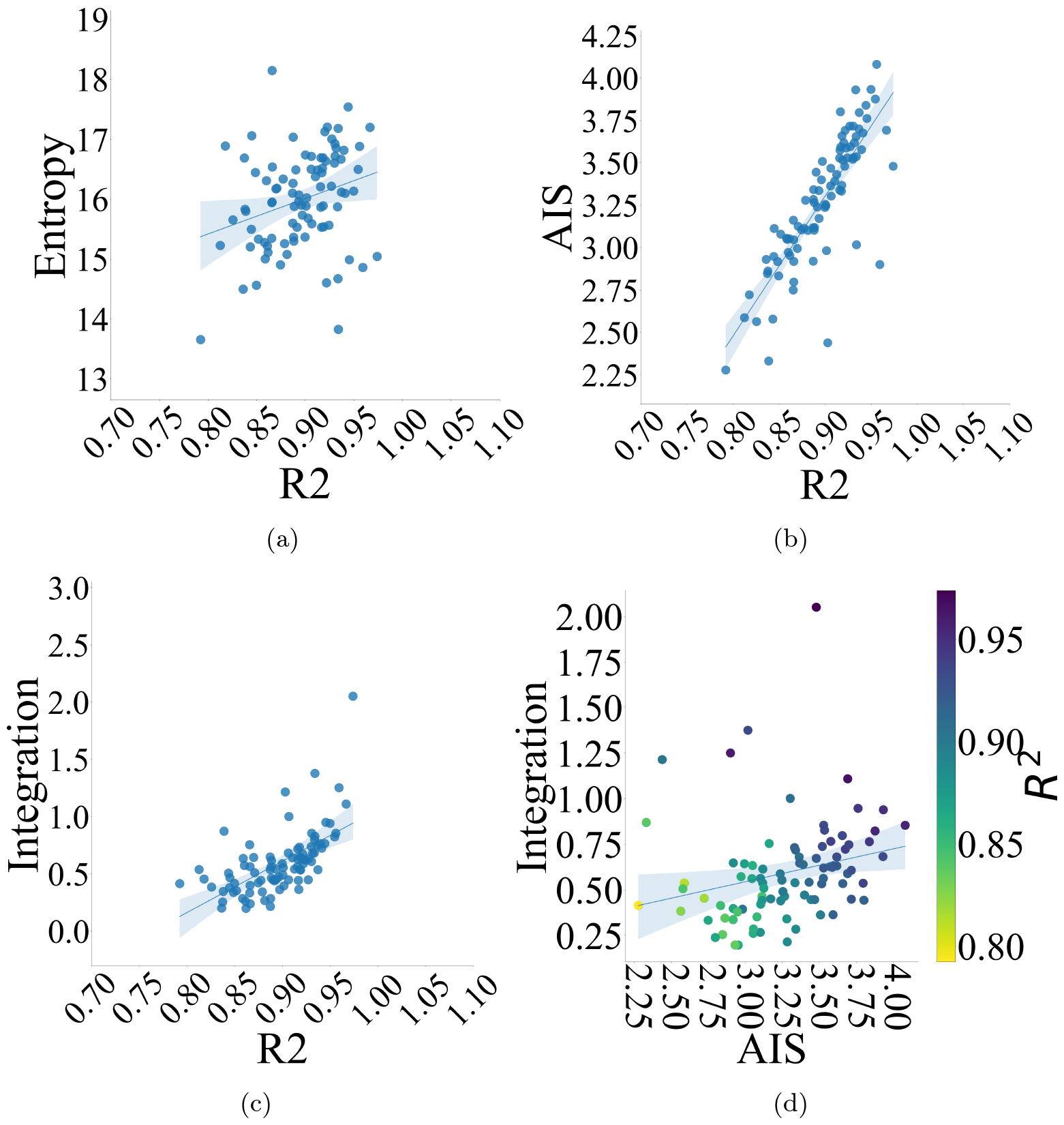
Correlations between (a) *R*^2^ and entropy (*H*) (b) *R*^2^ and *AIS* (c) *R*^2^ and *I* (d) *R*^2^ within *AIS* – *I* plane. These subplots verify that whereas AR accuracy *R*^2^ is a linear function of HR state-space *AIS* and *I*, it does not directly relate to the HR state-space entropy (i.e., variation).

Interestingly, *R*^2^ followed *AIS* – *I* covariation (Figure 10d, r = 0.4352, p = 2.03e^*−*05^) where the higher AR accuracy *R*^2^ corresponded to the larger HR state-space *AIS* and *I* values.

### 3.7 HR – RR Transfer Entropy and HR Dynamics

We observed a significant correlation between *RR → HR* and *HR → RR* (Figure 11, r = 0.5534, p = 1.86e^*−*08^). We further observed that within *RR → HR* – *HR → RR* plane, distribution of the AR accuracy *R*^2^ markedly resembled those of HR state-space *AIS* and *I*.

**Fig. 11:**
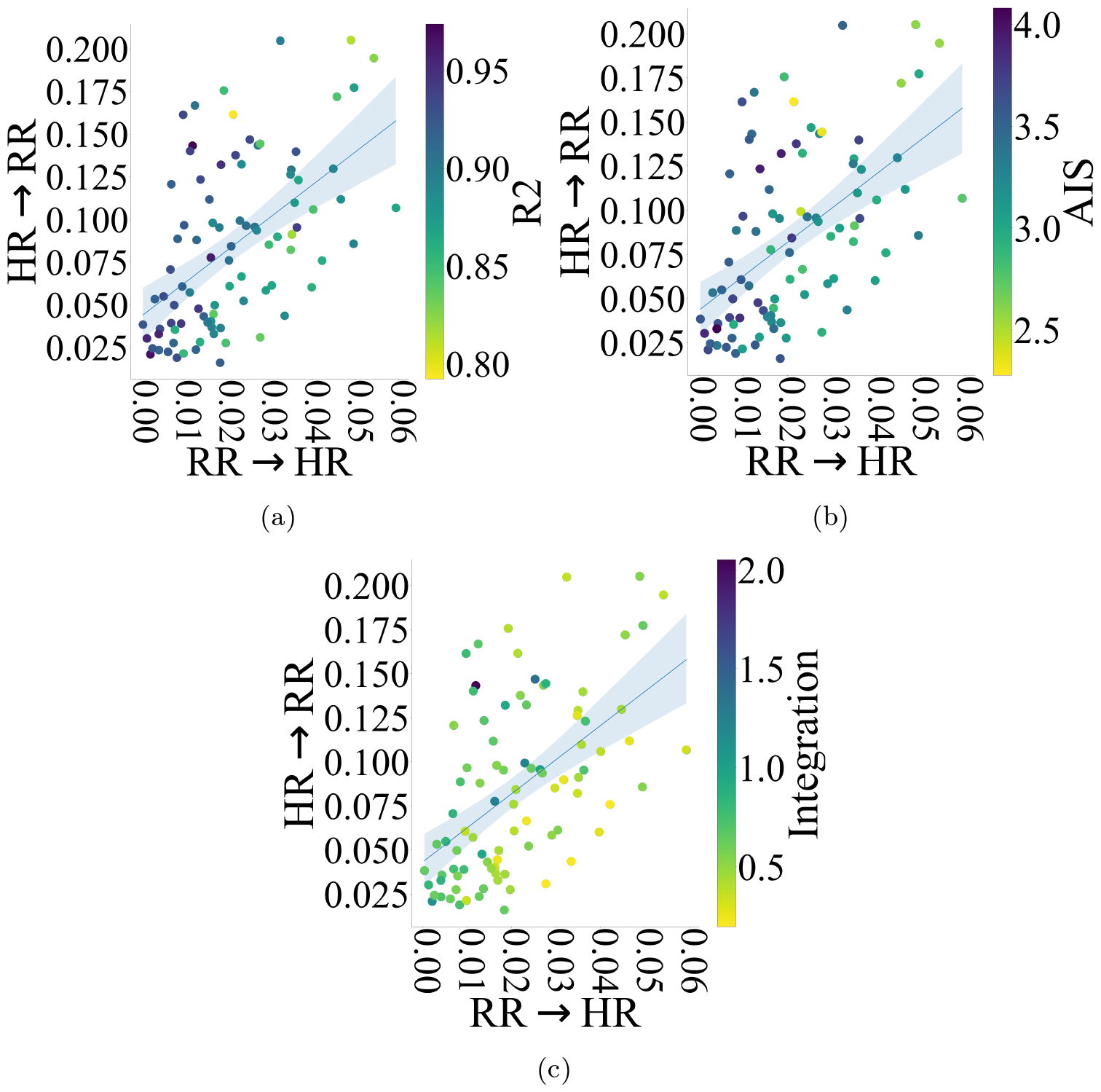
*RR → HR* – *HR → RR* plane with respect to (a) *R*^2^ (b) *AIS* (c) *I*. Distribution of AR accuracy *R*^2^ markedly resembled those of HR state-space *AIS* and *I*.

*RR → HR* was negatively correlated with AR accuracy *R*^2^ (Figures 12a and 12b, r = -0.6186, p = 1.05e^*−*10^), *AIS* (Figure 12c, r = -0.5856, p = 1.66e^*−*09^), and *I* (Figure 12d, r = -0.4914, p = 1.01e^*−*06^). Figure 12 also verifies that the higher AR accuracy *R*^2^ that corresponded to larger *AIS* and *I* (Figure 10d), pertained to the smaller *RR → HR*. It further indicates that the correspondence between AR accuracy *R*^2^ and *RR → HR* followed those of *AIS* and *I* in relation to *RR → HR*. However, such correspondences were absent in the case of *HR → RR* (SM1) as well as other HR measures (SM3).

**Fig. 12:**
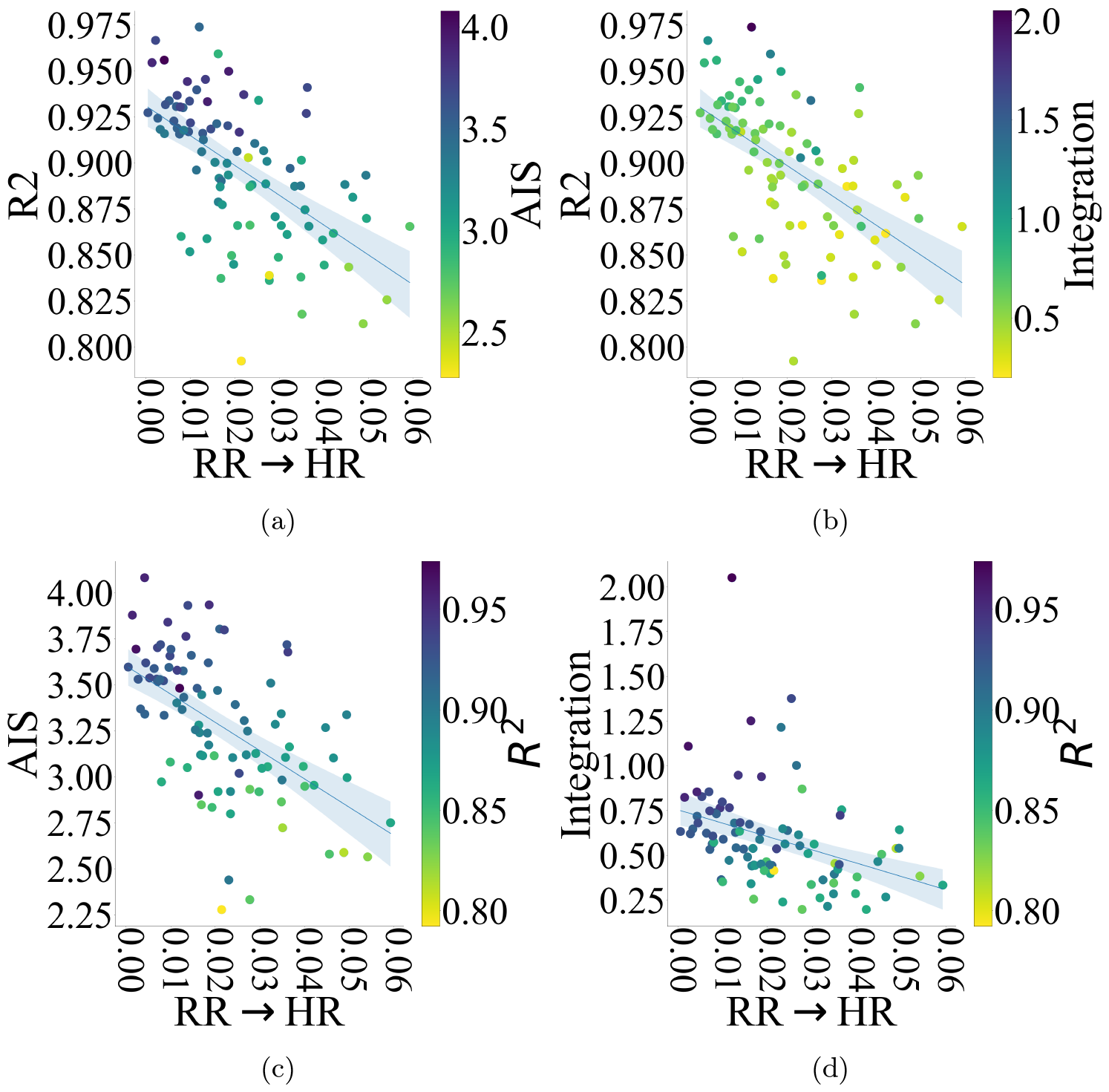
(a) *AIS* within *RR → HR* – *R*^2^ plane (b) *I* within *RR → HR* – *R*^2^ plane (c) *R*^2^ within *RR → HR* – *AIS* plane (d) *R*^2^ within *RR → HR* – *I* plane. These subplots indicate that higher *R*^2^ that corresponded to larger *AIS* and *I*, pertained to lower *RR → HR*. Such correspondences were absent in case of *HR → RR* (SM1).

### 3.8 HR Dynamics and Age

We observed no correlations between the participants’ age, on the one hand, and their AR accuracy *R*^2^ (Figure 13a, r = 0.109, p = 3.09e^*−*01^) or their HR state-space information dynamics *AIS* (Figure 13b, r=0.1948 p = 6.73e^*−*02^), on the other hand. The participants’ age also did not correlate with *I* (Figure 13c, r=-0.0672 p = 5.32e^*−*01^) or *H* (Figure 13d, r=-0.0283 p = 7.92e^*−*01^).

**Fig. 13:**
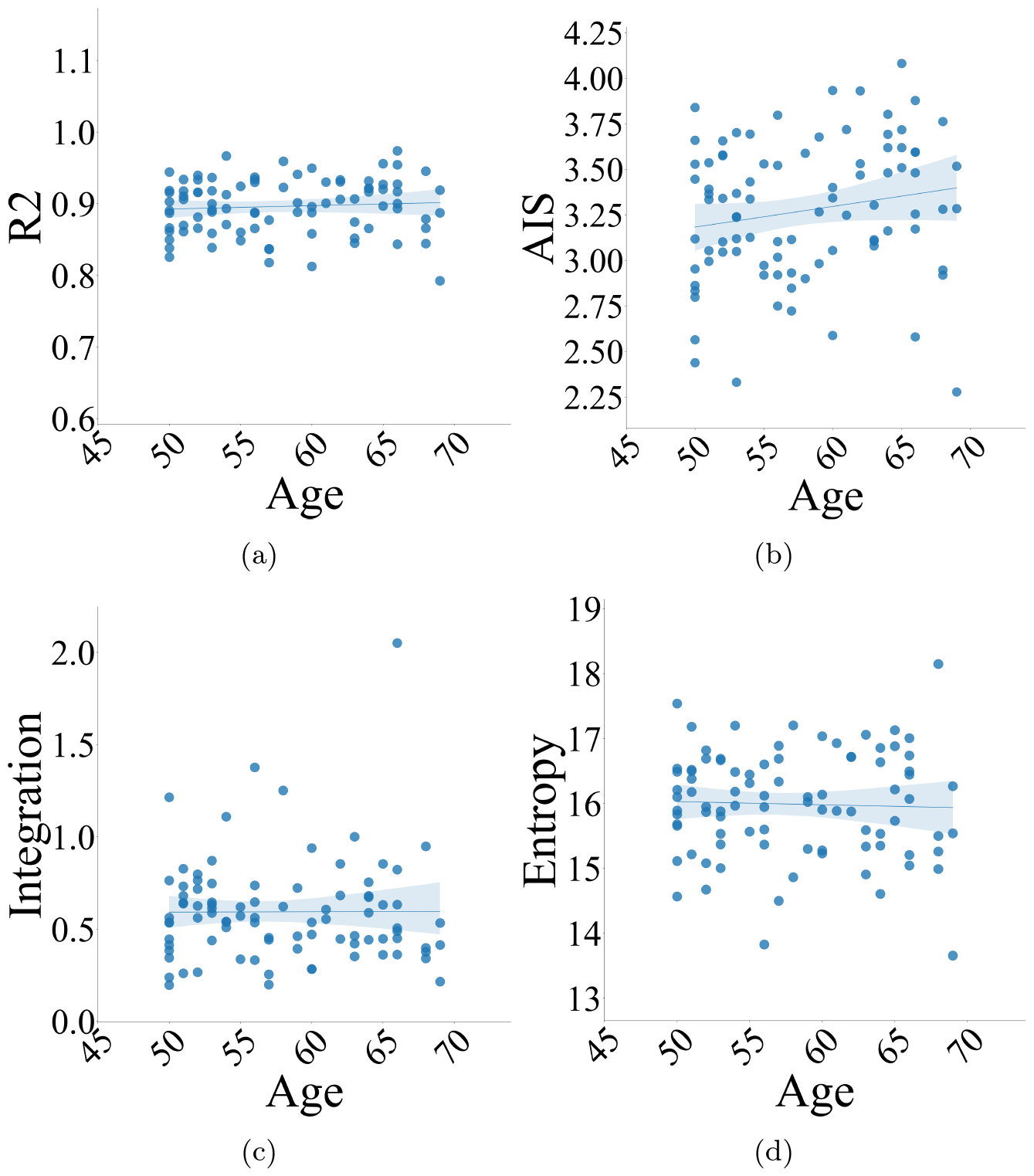
Correlation between participants’ age and (a) AR accuracy *R*^2^ (b) *AIS* (c) *I* (d) *H*. Participants’ age did not correlate with AR accuracy and/or HR state-space information dynamics.

Table 5 summarizes the Spearman’s rank correlation coefficients of our candidate biomarkers and the participants’ gender, age, smoking, alcohol consumption, and exercise habits.

**Table 5:** Spearman’s rank correlation coefficients associated with the HR AR accuracy *R*^2^, HR information dynamics *H, AIS, I*, transfer entropy *HR → RR* and *RR → HR*, and participants’ gender, age, smoking, alcohol consumption, and exercise habits (*⋆ p* = 3.63*e*^*−*02^, *⋆ ⋆ p* = 2.06*e*^*−*02^. These correlations did not pass Bonferroni correction).

|  | Gender | Age | Smoking | Alcohol Consumption | Exercise |
| --- | --- | --- | --- | --- | --- |
| $R^2$ | $r = -0.02$ | $r = 0.11$ | $r = 0.20$ | $r = -0.06$ | $r = -0.04$ |
| $H$ | $r = 0.06$ | $r = -0.03$ | $r = -0.08$ | $r = -0.13$ | $r = -0.06$ |
| $AIS$ | $r = -0.02$ | $r = 0.20$ | $r = 0.22^\star$ | $r = 0.03$ | $r = -0.08$ |
| $I$ | $r = -0.25^{\star\star}$ | $r = -0.06$ | $r = 0.02$ | $r = -0.13$ | $r = -0.21$ |
| $HR \rightarrow RR$ | $r = 0.02$ | $r = -0.01$ | $r = 0.08$ | $r = 0.02$ | $r = 0.07$ |
| $RR \rightarrow HR$ | $r = 0.09$ | $r = -0.03$ | $r = -0.07$ | $r = 0.20$ | $r = 0.17$ |

#### 3.8.1 Gender

We observed that the female participants had a significantly higher *I* than the male participants (Figure 14, test-statistics = 0.1199, p = 2.44e^*−*02^, g = 0.4190, Females: M = 0.6574, Mdn = 0.6272, SD = 0.2684, CI_95%_ = [0.4448, 1.0006], Males: M = 0.5432, Mdn = 0.5073, SD = 0.2756, CI_95%_ = [0.3796, 1.0596]). SM1 summarizes the gender’s non-significant differences with respect to other measures.

**Fig. 14:**
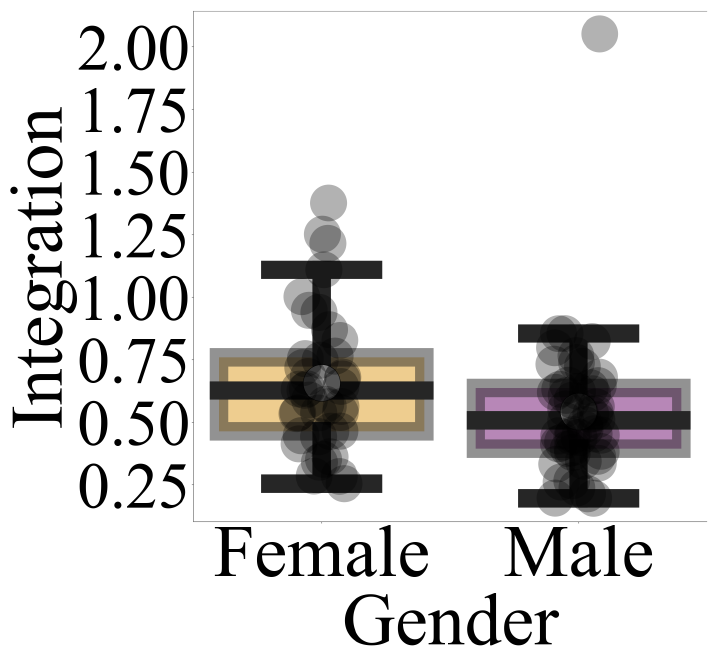
Gender differences with respect to *I* based on non-parametric bootstrap (10,000 repetitions) permutation test of difference in two groups’ Mdn.

#### 3.8.2 Age

We observed that *AIS* of the participants with whose age *≥* 60 was significantly higher than those whose age was *<* 60 (Figure 15, test-statistics = -0.3463, p = 6.80e^*−*03^, g = -0.4976, Age *≥* 60: M = 3.1959, Mdn = 3.1221, SD = 0.3512, CI_95%_ = [2.8931, 3.6161], Age *<* 60: M = 3.3792, Mdn = 3.4684, SD = 0.3914, CI_95%_ = [3.0687, 3.8925]). SM1 summarizes the age’s non-significant differences with respect to other measures.

**Fig. 15:**
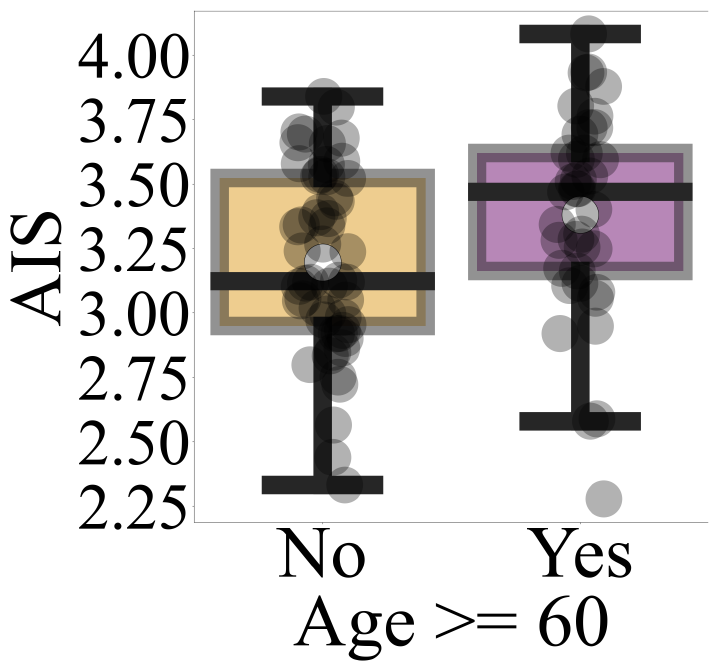
Individuals’ differences with respect to *AIS* and age group (i.e., age *≥* 60 vs. age *<* 60) based on non-parametric bootstrap (10,000 repetitions) permutation test of difference in two groups’ Mdn.

### 3.9 HR Information Dynamics and Habits

#### 3.9.1 Alcohol Consumption

Alcohol consumption had significant effects on *RR → HR* and HR state-space *H*. Compared to “Non-Drinkers,” “Drinkers” had significantly higher *RR → HR* (Figure 16a, test-statistics = -0.0070, p = 2.66e^*−*02^, g = -0.6309, Non-Drinkers: M= 0.0168, Mdn = 0.0164, SD = 0.0098, CI_95%_ = [0.0087, 0.0286], Drinkers: M = 0.0250, Mdn = 0.0234, SD = 0.0150, CI_95%_ = [0.0119, 0.0425]). This significant difference was also present in Wake period *RR → HR* (SM1).

**Fig. 16:**
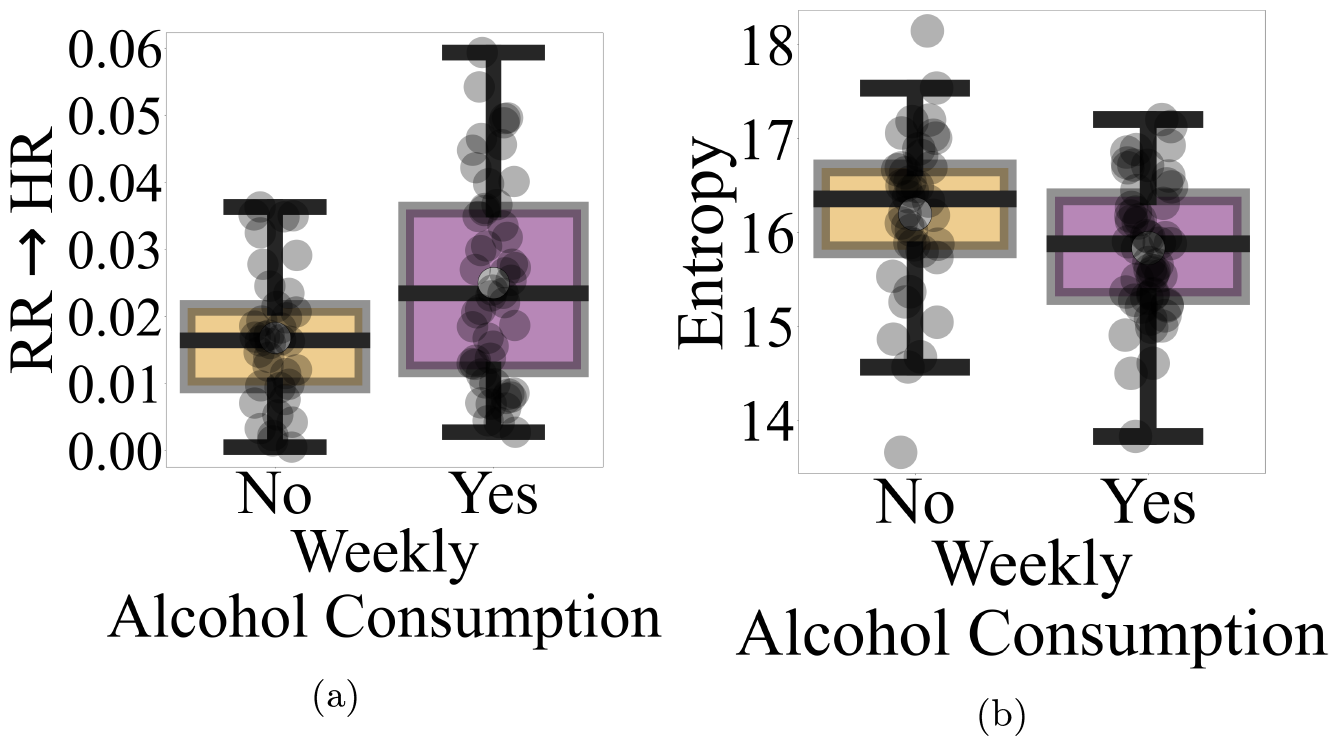
Effect of weekly alcohol consumption on (a) overall *RR → HR* (b) HR state-space *H*, based on non-parametric bootstrap (10,000 repetitions) permutation test of difference in two groups’ Mdn. Compared to “Non-Drinkers,” “Drinkers” had higher *RR → HR* and HR state-space *H*.

On the other hand, “Drinkers” had a significantly lower HR state-space *H* than “Non-Drinkers” (Figure 16b, test-statistics = 0.4762, p = 1.92e^*−*02^, g = 0.4569, Non-Drinkers: M = 16.1944, Mdn = 16.3556, SD = 0.8767, CI_95%_ = [15.4978, 17.3789], Drinkers: M = 15.8332, Mdn = 15.8795, SD = 0.7203, CI_95%_ = [15.217, 16.7261]). SM1 summarizes the alcohol’s non-significant differences with respect to other measures.

#### 3.9.2 Exercise

Exercise did not show any overall significant effects (SM1). However, when smoking and alcohol consumption habits were absent, it did show an effect on *RR → HR*. Specifically, we observed that those who exercised showed a significantly higher overall *RR → HR* (test-statistics = 0.0078, p = 5.80^*−*03^, g = 1.0884, Exercise: M = 0.02, Mdn = 0.02, SD = 0.01, CI_95%_ = [0.0142, 0.0346], No Exercise: M = 0.01, Mdn = 0.01, SD = 0.01, CI_95%_ = [0.006, 0.0232]). This significant difference was also present in the Wake period (SM1).

#### 3.9.3 Alcohol, Exercise and RR – HR

Interestingly, we observed no significant differences between those who exercised and did not consume alcohol versus those who exercised and consumed alcohol (SM1).

This trend of reduced *RR → HR* was preserved among the individuals who consumed alcohol and exercised versus those who consumed alcohol but did not exercise (SM1): the latter showed higher *RR → HR*. However, the difference between these two subgroups was non-significant (SM1).

Figure 17 depicts the grand-averages of the the participants’ HR state-space in each of the four groups (Section 2.5.7) i.e., (a) those who neither exercised nor consumed alcohol (Figure 17a) (b) those who did not exercise but consumed alcohol (Figure 17b) (c) those who exercised but did not consume alcohol (Figure 17c) (d) those who exercised as well as consumed alcohol (Figure 17d).

**Fig. 17:**
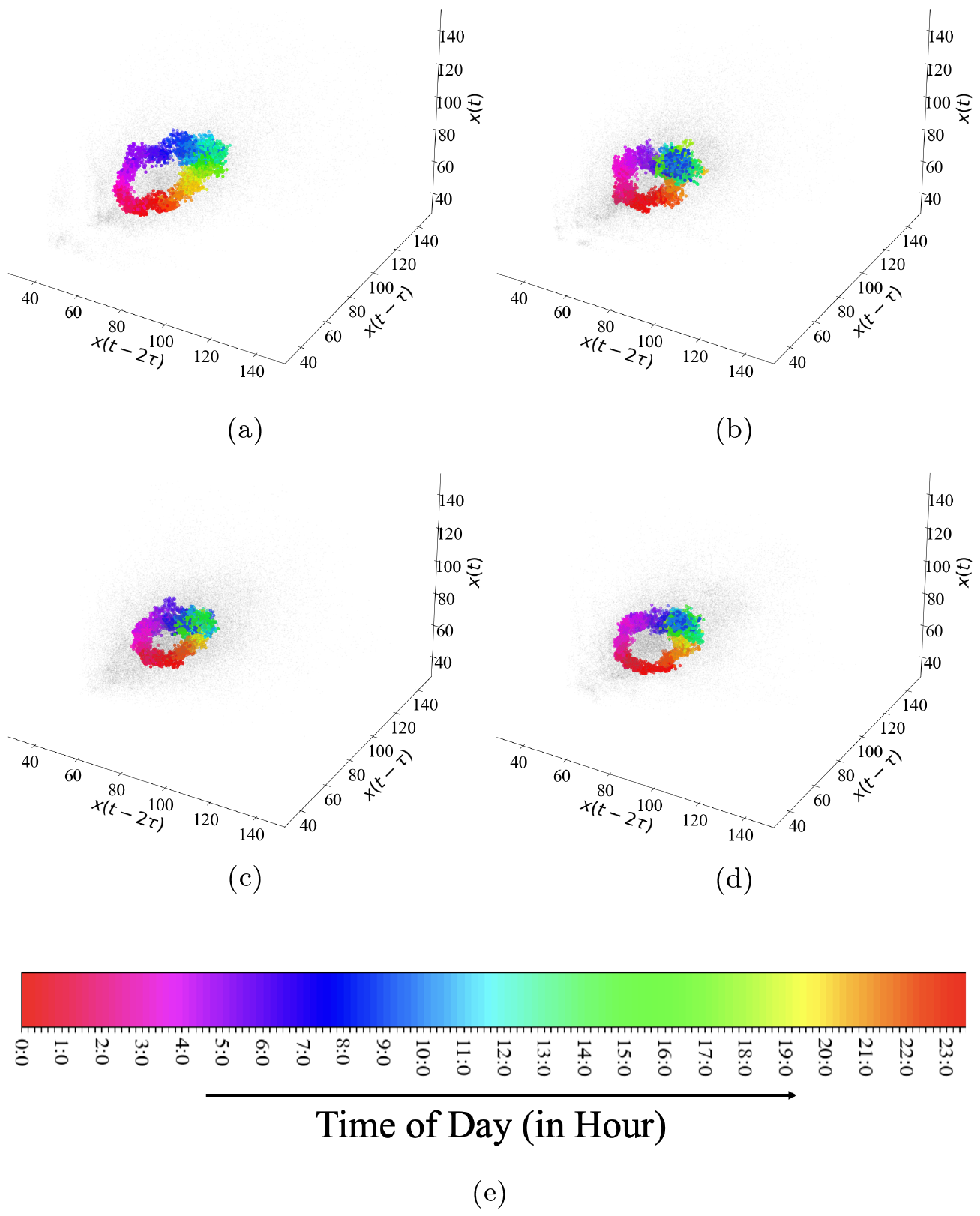
Grand-average of participants’ HR state-space (a) who neither exercised nor consumed alcohol (b) who did not exercise but consumed alcohol (c) who exercised but did not consume alcohol (d) who exercised as well as consumed alcohol. Data points are color-coded as per time-of-day (in an hourly basis) according to (e). See SM1 for plots of all participants. Clouds of black dots represent all participants’ HR data. A clear effect of alcohol on individuals’ HR circadian space is evident in this figure.

Two-factor ANOVA showed significant effects of alcohol (F = 6.4119, FDR-corrected p = 2.11e^*−*02^, *η*^2^ = 0.1106) and alcohol *×* exercise interaction (F = 9.6124, FDR-corrected p = 8.94e^*−*03^, *η*^2^ = 0.1657) on *RR → HR*. On the other hand, it revealed that exercise had no significant effect on *RR → HR* (F = 0.2261, FDR-corrected p = 0.6362, *η*^2^ = 0.0039).

Follow-up posthoc two-sample Welch test indicated significant differences in *RR → HR* between (Figure 18) (1) those who exercised but did not consume alcohol versus those who neither exercised nor consumed alcohol (t = 2.7885, FDR-corrected p = 2.88e^*−*02^, g = 1.0884) and (2) those who did not exercise but consumed alcohol versus those who neither exercised nor consumed alcohol (t = 3.5393, FDR-corrected p = 8.86e^*−*03^, g = 1.3778). These results were also significant in the case of the Wake period *RR → HR* (SM1).

**Fig. 18:**
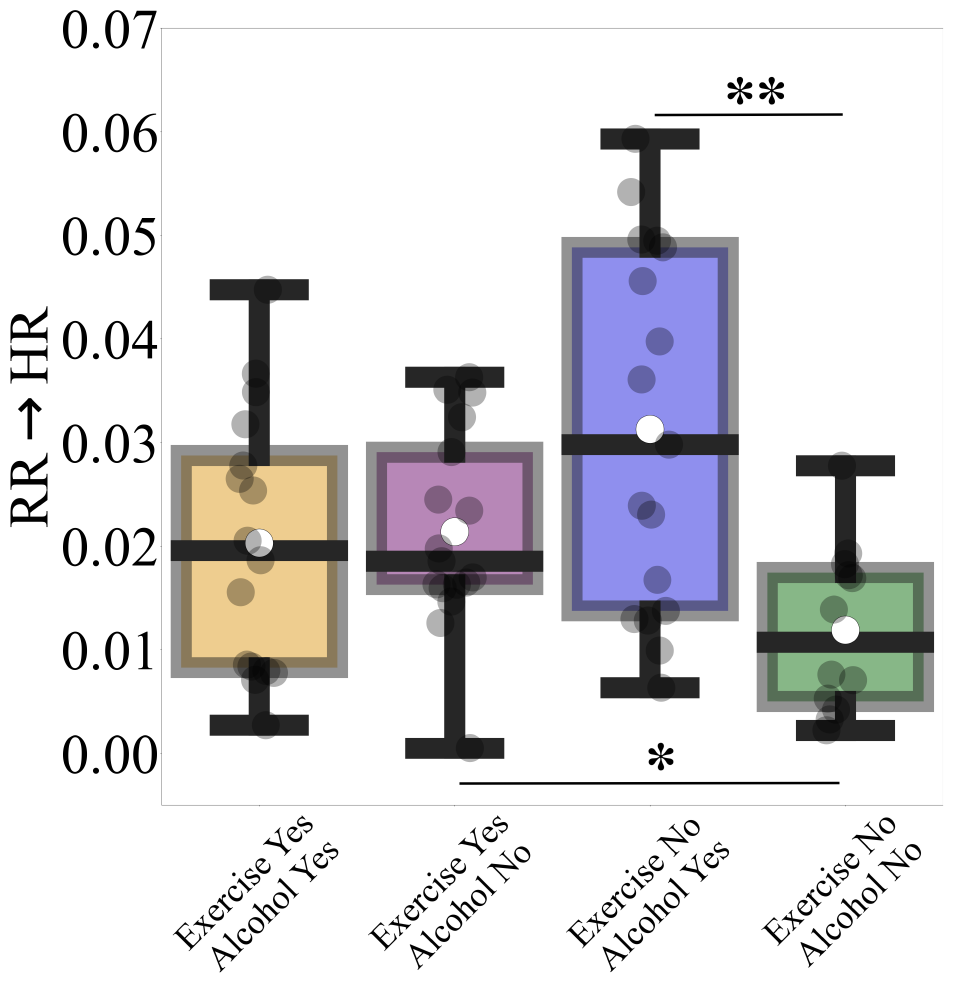
ANOVA analysis of alcohol – exercise interplay. Whereas alcohol consumption and its interaction with exercise were associated with significant effects, effect of exercise was non-significant. Additionally, there were significant differences between (1) those who exercised but did not consume alcohol versus those who neither exercised nor consumed alcohol and (2) those who did not exercise but consumed alcohol versus those who neither exercised nor consumed alcohol (*: p *<* 0.05, **: p *<* 0.01).

On the other hand, we found no significant differences between those who exercised and consumed alcohol versus (1) those who exercised but did not consume alcohol (F = -0.2796, FDR-corrected p = 7.82e^*−*01^, g = -0.1005), (2) those who did not exercise but consumed alcohol (F = -2.0421, FDR-corrected p = 6.74e^*−*02^, g = -0.7337), or (3) those who neither exercised nor consumed alcohol (F = 1.9989, FDR-corrected p = 6.74e^*−*02^, g = 0.7904). Last, we did not find any significant difference between those who exercised but did not consume alcohol and those who did not exercise but consumed alcohol (F = -2.0315, FDR-corrected p = 6.74e^*−*02^, g = -0.7182).

These results held true while using non-parametric Kruskal-Wallis with posthoc Wilcoxon rank-sum tests of significant differences among these four groups (SM1).

### 3.10 HR *AIS* – *RR → HR* Plane, Age and Habits

Individuals who consumed alcohol more frequently (Figure 19a, i.e., *≥* 4 days a week) were mostly associated with the “high *AIS* and low *RR → HR*” subspace (i.e., upper-left corner of the *AIS* – *RR → HR* plane).

**Fig. 19:**
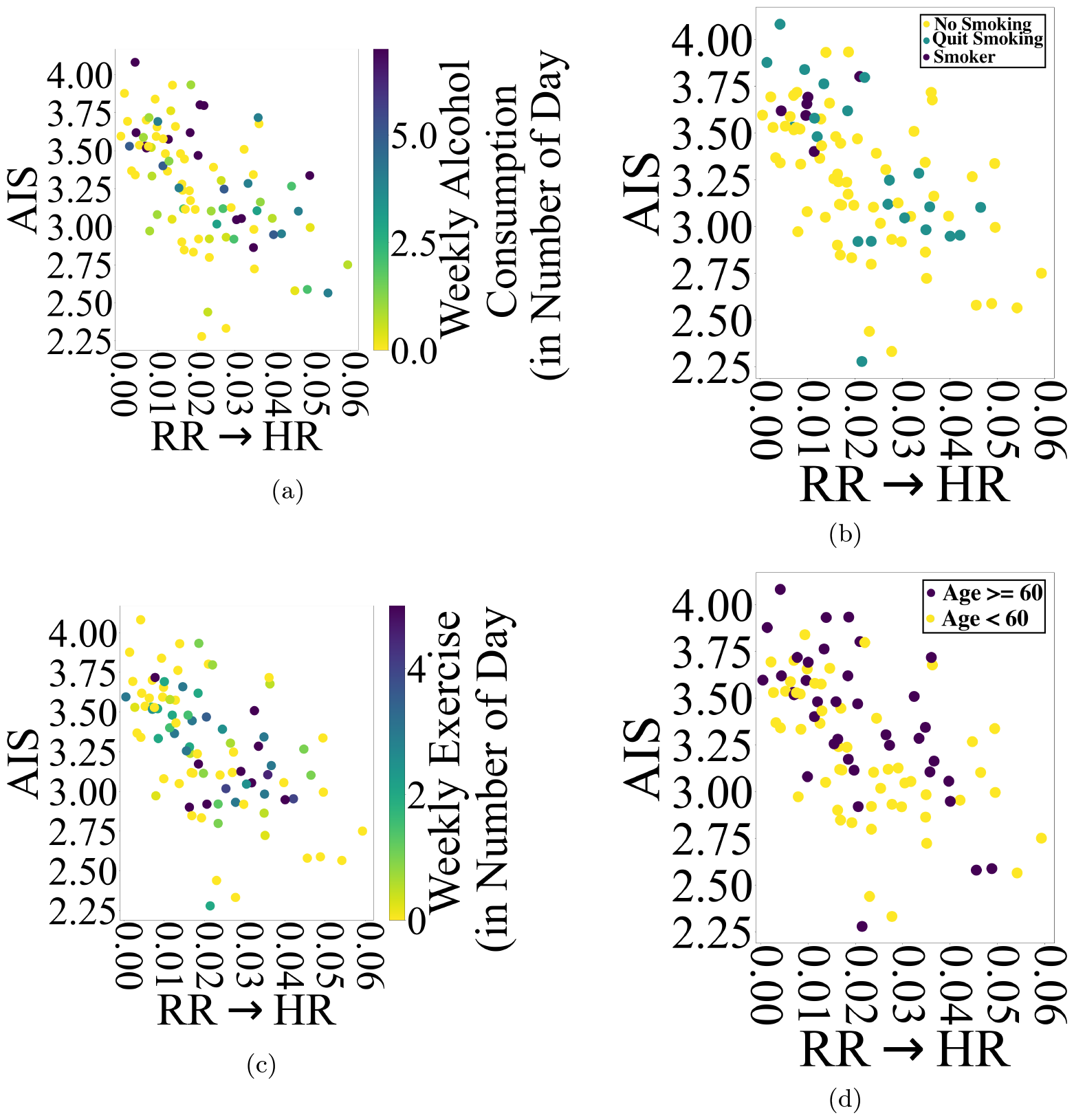
*AIS* – *RR → HR* plane with respect to participants’ (a) weekly alcohol consumption (b) weekly exercise (c) smoking habit (d) age *≥* 60.

Interestingly, this trend was also present among individuals with an active smoking habit (Figure 19b). Additionally, those who quit smoking formed two distinct clusters.

However, the lack of information on their date of quit, unfortunately, did not allow any further investigation of these two sub-clusters.

On the other hand, those individuals who exercised more frequently (Figure 19c, i.e., *≥* 4 days a week) were mostly distributed in the middle of *AIS* – *RR → HR* plane.

Considering the age group, the older individuals (Figure 19d, i.e., Age *≥* 60) were predominantly associated with the upper half of the *AIS* axis in the *AIS* – *RR → HR* plane.

Figure 20 shows the distribution of individuals within HR *AIS* – *RR → HR* plane and with respect to their exercise *×* alcohol consumption habits’ interplay. Whereas those who neither exercised nor consumed alcohol (in blue) were dominantly associated with high HR *AIS* and low *RR → HR*, those who did not exercise but consumed alcohol (in purple) were mostly in the lower quarter of *AIS* – *RR → HR* plane (i.e., low HR *AIS* and high *RR → HR*).

**Fig. 20:**
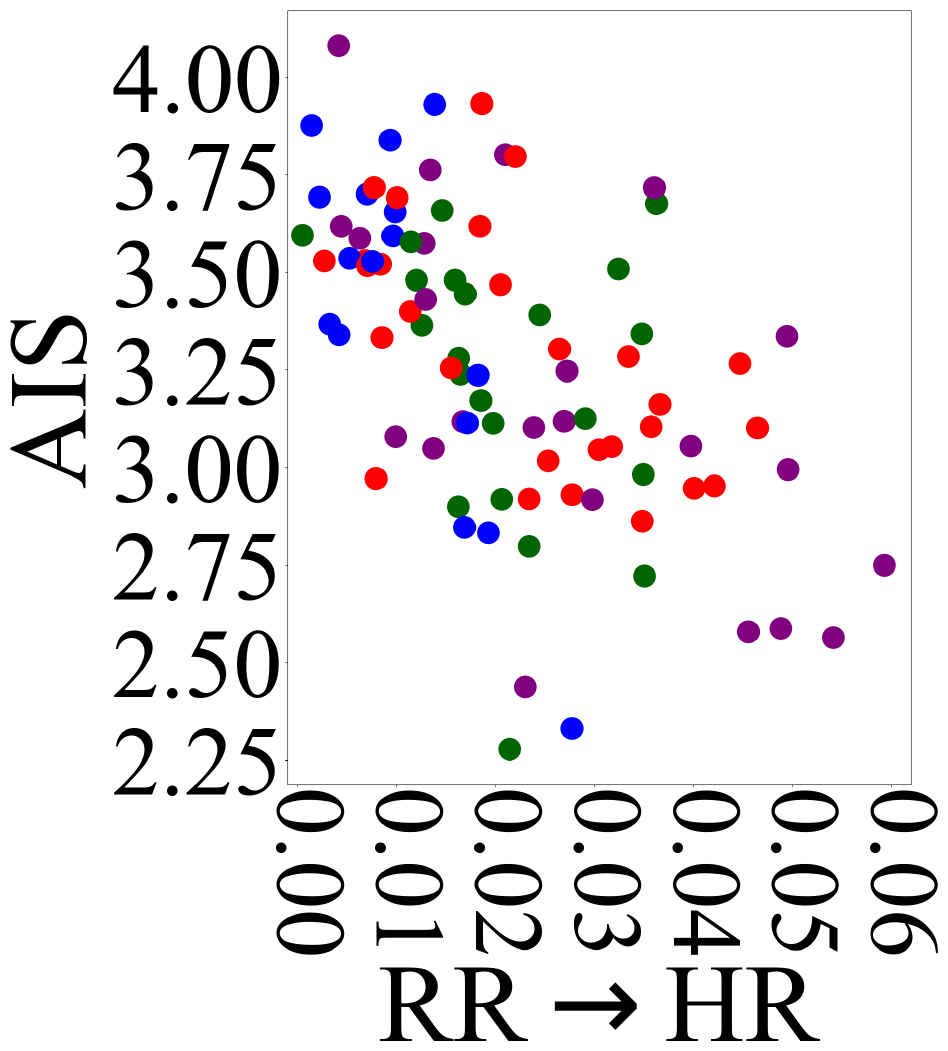
*AIS* – *RR → HR* plane with respect to participants’ Exercise *×* Alcohol Consumption habits. The color-coding in this plot corresponds to those who neither exercised nor consumed alcohol, those who exercised but did not consume alcohol, those who did not exercise but consumed alcohol, and those who exercised as well as consumed alcohol.

On the other hand, individuals who exercised but did not consume alcohol (in green) were mostly in the middle of *AIS* – *RR → HR* plane. Those who exercised as well as consumed alcohol (in red) were distributed along the HR *AIS* – *RR → HR* plane’s off-diagonal.

SM1 presents HR *AIS* – *RR → HR* plane with respect to average daily hours that each participant spent on sleeping, sitting/leaning, and standing/walking.

## 4 Discussion

In this study, we used freely-behaving human subjects’ longitudinal HR and RR recordings to investigate their utility for humans’ cardiorespiratory digital phenotyping [8, 83–86]. While doing so, we placed emphasis on interpretability of our analyses. We achieved this by quantifying the relation between various aspects of HR dynamics, on the one hand, and the HR – RR interplay, on the other hand.

Cohen and Taylor [35] observed that the imposition of broad assumptions (e.g., sole reliance on parametric approaches, over-generalization, etc.) and the lack of careful statistical analysis (e.g., uncorrected p-values, missing effect sizes, etc.) formed common characteristics of current research. According to Lombardi [52], only an integrative approach that incorporates both, linear and nonlinear properties of the cardiac function can provide proper means for quantification of its function and dynamics.

To this end, our study contributed to addressing these shortcomings by (1) accounting for linear and nonlinear HR dynamics through an integrative model-based and model-free analysis pipeline (2) adapting a non-parametric approach while applying these models (3) imposing stringent non-parametric statistics (4) taking into account the HR – RR interplay. Previous research considered the model-based and model-free methodologies [35, 48] and their combination [49, 50]. However, their results were limited in that they opted for parametric formalisms whose uncorrected statistics lacked effect sizes.

In the case of HR dynamics, we considered the reconstructed HR state-space than its univariate time series. Rhythmicity is a hallmark of many biological and physiological processes [87, 88]. Given the periodic nature of cardiac function, the study of HR dynamics within its reconstructed state-space was a self-evident natural choice [89].

This adaptation also enabled us to address another important shortcoming of the AR-based (i.e., model-based) approaches [35]: AR’s requirement of potentially large number of parameters. This requirement is imposed by the history length or lag (e.g., *μ*_*τ*_ = 128 in the present study) [90, 91].

The inherent periodicity of physiological processes allows for robust reconstruction of their (quasi)periodic sequences in terms of low-dimensional processes [87, 89, 92, 93]. In this respect, we reconstructed the participants’ HR state-space in 3 dimensions ^3^. This reduced the number of AR parameters to 4. Out of these, 3 parameters only were required for further analyses (i.e., discarding the bias term, given the mean-centered HR time series, Section 2.5). The use of state-space for the study of HR dynamics is not a novel approach. However, the previous research primarily adapted its parametric formulation (e.g., Gaussian assumption that allowed for closed-form analytic solution) [49, 50]. This, in turn, limited its utility for a thorough quantification of underlying nonlinear time-variant dynamics [33, 51, 92, 94, 95] of HR.

In the same vein, natural rhythms rarely exhibit absolute periodicity [87, p. 172] (e.g., they form non-uniform oscillations [93, Ch. 4], limit cycles [93, Ch. 7], etc.) [89]. Therefore, our quantification of HR state-space dynamics in terms of its nonlinear non-parametric informational properties allowed for accommodation of this aspect of HR in our analyses.

In this respect, it was interesting to note that our reconstructed state-space of participants’ HR (Figure 17) recovered the underlying circadian cycle of their HR dynamics. This verified the advantage of studying such nonlinear processes as HR within their reconstructed state-space than their univariate recordings [96, p. 242]. More importantly, the signature of individuals’ habits was evidently preserved within this reconstructed HR state-space. Precisely, we observed that the habit of alcohol consumption was associated with the reduced volume of the HR state-space.

### 4.1 HR AR and Information Dynamics

We observed that AR accuracy *R*^2^ correlated with HR state-space information-storage *AIS* and its long-range behavior *I*. We also observed that its variation within AR’s *w*_1_ – *w*_2_ and *w*_1_ – *w*_3_ planes closely resembled those of *AIS* and *I* within these planes. Moreover, we found that the increase in AR accuracy *R*^2^ was a direct function of the (positively) co-varing *AIS* and *I*. Interestingly, these results were absent in the case of other HR measures (see accompanying SM3).

These observations indicated that AR’s accuracy *R*^2^ was dependent on HR state-space information dynamics. This finding was in accord with Amari [97, Theorem 10.2, p. 224] who demonstrated that AR models follow the principle of maximum entropy [98, 99].

Precisely, AR accuracy intimately relates ^4^ to realization of the solution space (i.e., typical set [65, p. 59]). Crucially, the solution space’s volume and surface area are quantified by the entropy and the Fisher information, respectively [65, p. 247]. As a result, the dependence of the AR on HR information dynamics is explained by observing (1) that the Fisher information sets a lower bound on the mutual information (*MI*) ^5^ [100] (2) that *MI* operationalizes the *AIS* and *I* (i.e., equations (2) and (3)) (3) that *AIS* quantifies a process’s predictability [68]. i.e., the deviation of its components (in the present case, HR state-space dimensions) from independence and (4) that this (lack of) deviation from independence is analogous to the (presence of) long-range correlation [33, 101, 102] *I*) ^6^.

### 4.2 HR Information Dynamics and *RR → HR*

We also observed that HR state-space information dynamics was strongly *RR → HR* dependent. This resonated with the highly evolutionarily conserved respiratory entrainment of cardiovascular activity [53, 54] ^7^. It also aligned with the parasympathetic cardioinhibitory modulation of HR [105] by PreBötzinger complex neurons (preBötC) [106]. PreBötC plays a critical role in breathing [107] via regulating the respiration. Menuet et al. [108] demonstrated that preBötC’s inhibition resulted in suppression of the respiratory sinus arrhythmia (RSA). RSA modulates the HR and its synchrony with RR [55, 56]. It also reduces cardiac energetic cost [57, 58]. As a result, RSA is considered a measure of cardiac age [59] and control [60].

From a broader perspective, this dependence provided further evidence for the critical role of the inhibition in generation of (cardiorespiratory) rhythms [109]. In this respect, the absence of any significant contribution from *HR → RR* in our results echoed its association with cardiorespiratory pathology (i.e., sleep apnea [110]) [70] [111, p. 178].

Our results on the pivotal role of *RR → HR* in HR dynamics addressed the lack of focus on humans’ cardiorespiratory phenotyping and its critical role in health [53–60] and disease [61–63].

### 4.3 HR State-Space *I* and Gender-Differences

Gender differences in cardiac function is well-documented [38, 40, 41, 63]. Research has found that HRV was significantly lower in females than in males, that for individuals under the age of 50, HR was faster in females than in males, that whereas females exhibited a higher parasympathetic activity, males had higher sympathetic activity, and that these differences were disappeared after the age of 50 [41, 112–115].

Considering the participants’ age in the present study (Section 2.1), we also observed this diminishing gender differences by age in HR (Section 2.1). Our analyses also extended this observation to the case of RR (Section 2.1), HR – RR correlation (Section 3.2), and HR – RR cross-correlation (Section 3.3).

On the other hand, we found that females had higher HR state-space *I* than males. This posited *I* as a digital marker of cardiac function that was not affected by aging. The previous research also found that females had higher parasympathetic cardiac autonomic activity than males [115]. Therefore, the observed higher *I* in females than males may suggest that *I* was more sensitive to parasympathetic than sympathetic activity. This suggestion found support in females’ significantly lower sleep-time HR than males (SM1) potentially due to their parasympathetic dominance during the sleep [37, p. 159].

### 4.4 HR State-Space *AIS*: a Digital Phenotype of the Aging Heart

We observed that aging was associated with an increase in HR state-space *AIS*. This resonated with characterization of the biological aging with a progressive impairment of the physiological control mechanisms [43, 116]. These control mechanisms and their dynamics are necessary for maintaining the systems-level homeostatsis [33, 45, 51, 116]. Ample evidence demonstrates the occurrence of such an impairment in cardiovascular [42–46] and respiratory [47, 63] mechanisms.

In this respect, *AIS* complements these observations by attributing an increased cardiac regularity to the process of aging. This suggestion aligns with the proposal that associated the process of aging and disease with the replacement of rhythmic processes by relatively constant dynamics [87, p. 173].

### 4.5 *RR → HR*: a Cardiorespiratory Digital Phenotype

We found that the habit of alcohol consumption was identifiable by a significant increase in *RR → HR*. This observation echoed the strong association between the alcohol and hypertension [117–120].

Interestingly, the effect of increased *RR → HR* was also present in the reconstructed HR state-space of those who reported the alcohol consumption as their primary habit. Specifically, the reconstructed HR state-space of this sub-sample showed a visible reduction in its area and increased perturbation that was more dominant during the waking period (Figure 17b).

The World Health Organization (WHO) considered the alcohol consumption to be responsible for 3 million deaths in 2016 globally and 5.1% of the burden of disease and injury [121]. Furthermore, recent findings associated this habit with 740,000 new cancer cases each year [122, 123]. A third of such cases were attributable to light to moderate drinking [124].

In view of these findings, *RR → HR* can be construed as a digital biomarker of the alcohol’s detrimental effect on cardiorespiratory mechanism. This suggestion becomes more plausible, considering that *RR → HR* was agnostic to the amount of alcohol consumed by the individuals. Concretely, the increase in *RR → HR* was primarily due to the presence or absence of this habit in the participants’ lifestyle and activity. *RR → HR* was also sensitive to the effect of exercise. In the absence of drinking and smoking habits, those who exercised showed significantly higher *RR → HR* than those who did not exercise. Interestingly, this increase in *RR → HR* did not differ between those who exercised and consumed alcohol, on the one hand, and those who did not consume alcohol but did exercise, on the other hand. This implied that exercise may benefit those who consume alcohol by (down-) regulating their alcohol-induced increase in respiratory modulation of their cardiac function [117–120].

Taken together, our results posited *RR → HR* as a digital marker of cardiorespiratory (mal)function [61, 125–127]. For instance, it can prove useful in monitoring those who suffer from hypertension [61, 128]. In the case of the alcohol consumption [117–120], this measure may present a prognostic marker of recovery during rehabilitation.

*RR → HR* can also serve as a prognostic digital marker of recovery from cardiorespiratory complication(s) through exercise. This suggestion becomes plausible, given the association between decreased respiratory modulation of the cardiac function and the cardiorespiratory pathology [129, 130]. Considering the benefit of exercise-therapy to COVID-19 recovery [131], *RR → HR* can help monitor the patients’ progress and their recovery by quantifying the degree of increase in their respiratory modulation of their cardiac function.

## 5 Concluding Remarks

In this study, we showed the potentials that the HR state-space information dynamics can offer to the solution concept of humans’ cardiorespiratory digital phenotyping. We identified *AIS* as an age-related digital biomarker that associated the process of aging with the increased regularity in HR dynamics [111] [87, p. 173].

More importantly, we proposed *RR → HR* as a digital biomarker of respiratory modulation of cardiac function. We presented its potential for quantifying the impact of alcohol on this mechanism. Our results also suggested its capacity for measuring the effect of physical activity to potentially compensate for the alcohol’s impact on the cardiorespiratory mechanism [117–120]. These results posited *RR → HR* as a digital diagnosis/prognosis marker of cardiorespiratory pathology [61, 62, 117–120, 125–128, 132, 133].

We also realized *I* as a digital biomarker (1) that quantified the gender differences in HR long-range behavior (2) that was invariant to the effect of aging [38, 41] and (3) that potentially pertained to the gender-specific parasympathetic activity [115] [37, p. 159].

These measures quantified different factors that affected distinct aspects of cardirespiratory mechanism. They included aging, gender, and habits. This indicated that these measures were functionally reduced with whose quantification of cardiorespiratory function nonredundant. To this end, they were validated as (digital) biomarkers [30, 31]. As a result, they contributed to the objective of phentopying research. That is, the discovery of the set of organism’s traits and characteristics that can be objectively measured and evaluated as indicators of its biological processes and pathology [1].

Torous et al. [15] formalized the ultimate goal of humans’ digital phenotyping as “moment-by-moment quantification of the individual-level human phenotype in situ using data from personal digital devices.” This goal still remains illusive. It goes without saying that a number of studies have utilized the digital technologies to successfully identify several behavioral signatures of physiological phenotypes. These included the pain severity [20], sleep patterns [21], behavioral traits in smartphone use [22], and the potential correlate of HRV and physical activity [23]. However, the extension of these results to continuous monitoring of humans’ physiology is nontrivial. Our results are of no exception. For instance, it remains unclear how *RR → HR* diagnosis / prognosis potentials can be extended to round-the-clock monitoring of cardiorespiratory malfunction in individuals at risk or during rehabilitation. This becomes even more difficult in the case of freely behaving daily life routines and activities.

One important limitation of the present study is the absence of data on individuals’ comorbidities. Such a knowledge can prove invaluable for real-time monitoring of cardiorespiratory function at the individual-level. In particular, it can allow for better estimate and adjustment of the biomarkers and phenotypic measures. This, in turn, reduces the possibility of inflated outcomes that could be attributable to other issues than cardiorespiratory function at any specific moment in time. Along the same direction, it is also critical to incorporate any possible family history of cardiovascular and cardiorespiratory pathology in such analyses. This necessity is apparent, given the role of genetic factors and heritability in HR [134] and its variability [135–138].

In the same vein, our study relied on the individuals’ self-report (Section 2.4). As a reult, it lacked proper information about individuals’ personality and behavioral traits. This information can be effectively collected through standardized questionnaires [139– 143]. Their use can prove invaluable for reducing the confounding effects of people tendency to overestimate their health and abilities [144–146].

Additionally, our sample did not include younger adults (i.e., age *<* 50). A sample with a better age heterogeneity can allow for refining the extent of observed changes and variation in measured biomarkers. This, in turn, can lead to more informed conclusion on underlying factors that can affect the cardiorespiratory function.

The sampling rate of VitalPatch (i.e., 0.25 Hz) was in accord with one of two cardiovascular preferential frequencies [147, p. 17]. However, it remains open to future investigation to further verify whether and how an increase in sampling rate may allow for improving the present results and observations.

## Supplementary information

This manuscript contains supplementary materials (SM) available in accompanying files SM1, SM2, and SM3.

## Acknowledgments

The authors would like to extend their gratitude to Noriyuki Ando, Representative Director, President & CEO, Research Strategy Planning Department, Suntory Global Innovation Center Limited, Suntory, Kyoto, whose commendable efforts in organizing and fostering this collaborative project significantly contributed to the successful completion of this research.

## Declarations

### Ethics approval

The present results was based on a joint research between OIST and Suntory Global Innovation Center. This research was approved by the Ethics Review Committee at OIST (Approval Number: HSR-2021-010-2), based on Suntory’s research plan SIC-2021-07-DBS (The Ethics Committee of Miura Clinic, Medical Corporation Kanonkai (IRB No. 17000161)).

### Consent to participate

All participants provided written consent in accordance with Declaration of Helsinki.

### Consent for publication

All authors agreed with the publication of these results.

### Authors’ contribution

H.M. designed and conducted the experiment. S.K. and K.D. carried out the analyses. S.K., S.T., and K.D. prepared the manuscript.

## Footnotes

1 A biomarker is a characteristic that can be objectively measured and evaluated as an indicator of typical biological processes, pathogenic processes or pharmacological responses to a therapeutic intervention [30, 31].

2 In SM3, we reported standard deviation (SD) instead.

3 The state of a (forced) harmonic oscillator is indeed three dimensional [93, p. 7].

4 This relation is through the Fisher Information. The Fisher information quantifies the accuracy of an unbiased estimator [65, p. 393–394]. It is a measure of amount of “information” about the model’s parameters (in the present scenario AR’s *w*_1_, *w*_2_, and *w*_3_) that is present in the data, thereby providing a lower bound on the error in estimating these parameters from the data [65, p. 397

5 *MI* magnitude is ultimately determined by the entropy of the variables involved (i.e., *MI*(*X*; *Y*) = *H*(*X*) + *H*(*Y*) *− H*(*X, Y*).

6 It is worth noting that the solution space (i.e., typical set [65, p. 59]) is associated with the target densities that exhibit strong correlations [103, p. 393].

7 On a time scale between 1 – 30 seconds, parasympathetic and sympathetic activities in humans at rest is mainly modulated by respiratory activity [104, p. 193]

