## Supplementary Materials 1 for "Information Dynamics of the Heart and Respiration Rates: a Novel Venue for Digital Phenotyping in Humans"

### Supplementary Materials 1 (SM1):
### Respiratory Modulation of the Heart Rate: a Potential Marker of the Effect of Alcohol on Cardiorespiratory Function in Human

Soheil Keshmiri[1*], Sutashu Tomonaga[2], Haruo Mizutani[3], Kenji Doya[2]

[1]Optical Neuroimaging Unit, Okinawa Institute of Science and Technology, Okinawa, Japan.
[2]Neural Computation Unit (NCU), Okinawa Institute of Science and Technology, Okinawa, Japan.
[3]Suntory Global Innovation Center Limited (SGIC), Suntory, Kyoto, Japan.

Contributing authors:;
;

#### Contents

**1 HR State-Space Delay Embedding $\tau$** **2**

**2 $HR \rightarrow RR$ and $RR \rightarrow HR$ Time Lag $\kappa$** **2**

**3 $\mu_{R^2}$ Estimation** **3**

**4 $\mu_{HR}$ versus AR Parameters** **3**

**5 $\mu_{HR}$ versus HR's $H$, $AIS$, and $I$** **3**

**6 AR Accuracy Reflects the Covariation Between HR State-Space's Present vs. Its Past Dimensions** **5**

**7 $AIS$ and $I$ within $w_1 - w_3$ and $w_2 - w_3$ Planes** 5

**8 $R^2$ within $w_2 - w_3$ Plane** 6

**9 $HR \rightarrow RR$, $AIS$, $I$, and $R^2$** 8

**10 Effect of Weekly Alcohol Consumption on $RR \rightarrow HR$ During Wake** 12

**11 Alcohol Consumption and Exercise** 12
- 11.1 Individuals Who Exercised But Differed in Their Alcohol Consumption Habit . . . 13
- 11.2 Individuals Who Consumed Alcohol But Differed in Their Exercise Habit 15

**12 Factor Analysis of the Alcohol – Exercise Interplay: Wake $RR \rightarrow HR$** 15

**13 Non-Parametric Analysis of Alcohol – Exercise Interplay Interplay: $RR \rightarrow HR$** 17

**15 HR $AIS$ – $RR \rightarrow HR$ Plane, Average Daily Sleep, Walking/Standing, and Sitting/Leaning** 25

### 1 HR State-Space Delay Embedding $\tau$

Figure 1 plots the result of the bootstrap (10,000 repetitions) estimate of $\mu_\tau$ from the participants' respective HR state-space delay embedding $\tau$ (see main manuscript) at 95% CI (M = 128.1602, Mdn = 128.2472, $CI_{95\%}$ = [119.6739, 136.2921]). We used this $\mu_\tau = 128$ (i.e., 2 hours and 8 minutes) as fixed $\tau$ value to reconstruct all participants' final HR state-space (i.e., $\tau = \mu_\tau$, for all the participants).

### 2 $HR \rightarrow RR$ and $RR \rightarrow HR$ Time Lag $\kappa$

Figures 2a and 2c show the distribution of the participants' $HR \rightarrow RR$ and $RR \rightarrow HR$ time lag $\kappa$ ($HR \rightarrow RR$: M = 5.5831, SD = 3.5096, Mdn = 6.00, $CI_{95\%}$ = [3.1162, 3.9885], Min = 1.00, Max = 14.00, $RR \rightarrow HR$: M = 15.2360, SD = 9.3352, Mdn = 15.00, $CI_{95\%}$ = [8.2634, 11.0363], Min = 1.00, Max = 45.00). Figures 2b and 2d plot the results of the bootstrap (10,000 repetitions) estimates of their respective $\mu_\kappa$ at 95% CI ($HR \rightarrow RR$: M = 5.5833, Mdn = 5.5831, CI = [4.7640, 6.2135], $RR \rightarrow HR$: M = 15.2304, Mdn = 15.2247, $CI_{95\%}$ = [13.3483, 17.1910]).

#### 3 $\mu_{R^2}$ Estimation

Figure 3a plots the individuals' AR accuracy $R^2$ (M = 0.8962, SD = 0.03806, Mdn = 0.9002, CI = [0.0338, 0.0437]). Figures 3b shows the result of the bootstrap (10,000 repetitions) estimate of $\mu_{R^2}$ at 95% CI (M = 0.8962, Mdn = 0.8963, SD = 0.0041, CI = [0.8882, 0.9039]).

#### 4 $\mu_{HR}$ versus AR Parameters

We found no correlation between $\mu_{HR}$ and AR parameters $W_0$ (Figure 4a, r = 0.1811, p = 8.94e$^{-02}$), $W_1$ (Figure 4c, r = -0.1014, p = 3.44e$^{-01}$), $W_2$ (Figure 4d, r = -0.1129, p = 2.92e$^{-01}$), and $W_3$ (Figure 4b, r = 0.1108, p = 3.01e$^{-01}$).

#### 5 $\mu_{HR}$ versus HR's $H$, $AIS$, and $I$

We found no correlation between $\mu_{HR}$ and $AIS$ (Figure 5b, r = 0.1608, p = 1.32$^{-01}$) and $I$ (Figure 5c, r = 0.0289, p = 7.88$^{-01}$).

On the other hand, we observed a significant correlation between $\mu_{HR}$ and $H$ (Figure 5a, r = 0.4263, p = 3.11e$^{-05}$). However, this correlation did not show any correspondence to AR's $R^2$ (Figure 5d), $HR \rightarrow RR$ (Figure 5e), $RR \rightarrow HR$ (Figure 5f), $AIS$ (Figure 5g), or $I$ (Figure 5h).

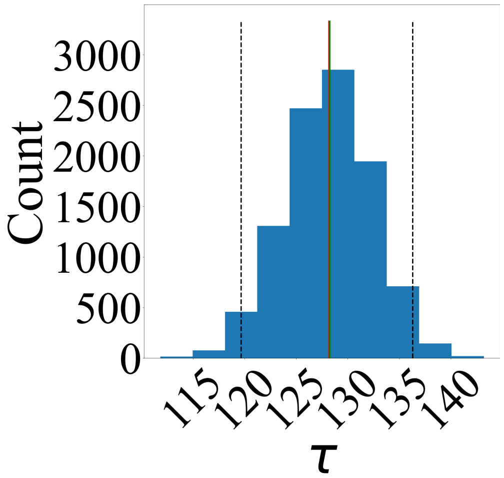

**Fig. 1**: Bootstrap (10,000 repetitions) estimate of HR state-space delay embedding $\mu_\tau$ at 95% confidence interval (CI). Bootstrapping participants' respective $\tau$ yielded $\mu_\tau$ = 128 (i.e., 2 hours and 8 minutes). We set every participants' $\tau = \mu_\tau$ to reconstruct their final HR state-space. In this plot, red and green lines mark estimated $\tau$'s mean (M) and median (Mdn), respectively (M = 128.1602, Mdn = 128.2472). Black dashed lines mark lower and upper $CI_{95\%}$ ($CI_{95\%}$ = [119.6739, 136.2921]).

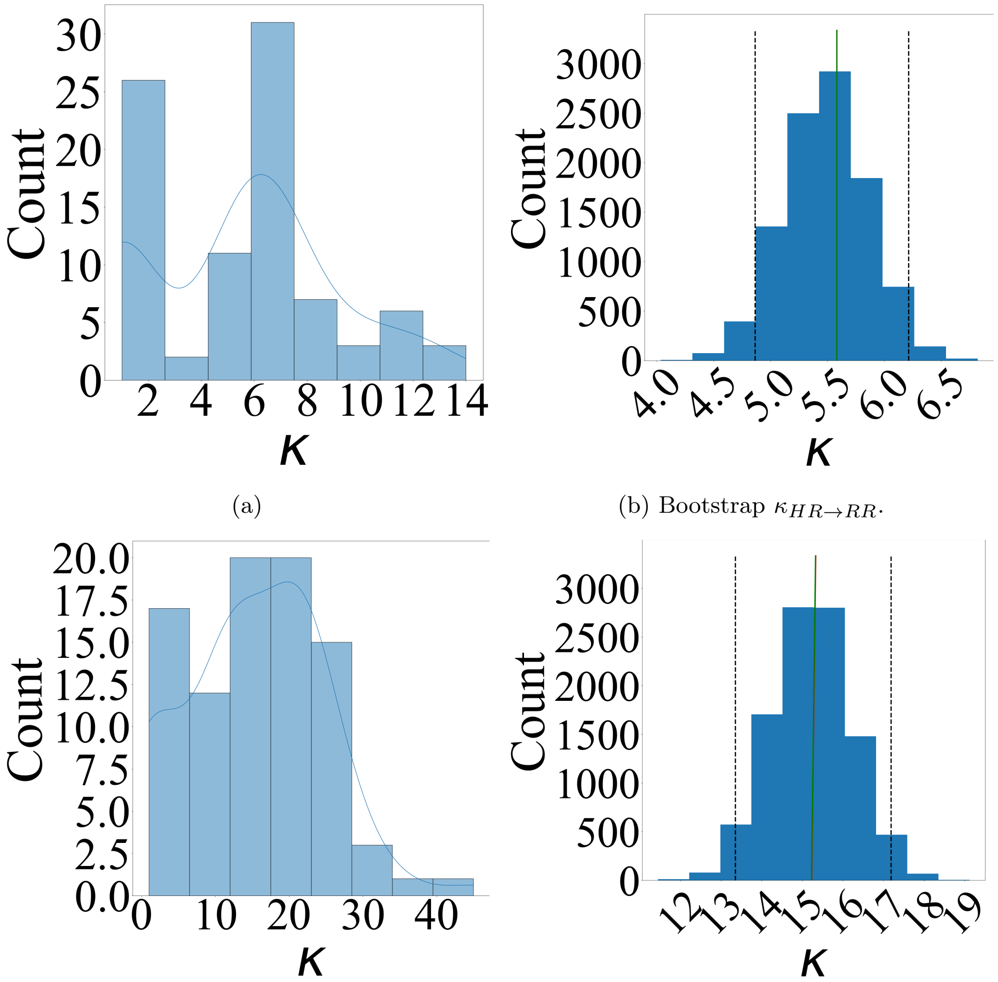

(a)

(b) Bootstrap $\kappa_{HR \to RR}$.

(c)

(d)

**Fig. 2**: Estimation of TE's time lag $\kappa$. (a) Individuals' $\kappa_{HR \to RR}$ distribution (b) Bootstrap (10,000 repetitions) estimate of $\mu_{\kappa_{HR \to RR}}$ at 95% CI (c) Individuals' $\kappa_{RR \to HR}$ distribution (d) Bootstrap (10,000 repetitions) estimate of $\mu_{\kappa_{RR \to HR}}$ at 95% CI. To obtain individuals' TE maximizing time lag $\kappa$ (both, $HR \to RR$ and $RR \to HR$) we performed a brute-force search [**?** ] where we considered same range of values as in case of HR state-space delay embedding $\tau$ (i.e., $\kappa \in [1, \ldots, 240]$ or 1 through 240 minutes). In subplots (b) and (d), red and green lines mark estimated $\kappa$'s mean (M) and median (Mdn), respectively ($HR \to RR$: M = 5.5833, Mdn = 5.5831, $RR \to HR$: M = 15.2304, Mdn = 15.2247). Black dashed lines mark lower and upper $\text{CI}_{95\%}$ in these subplots ($HR \to RR$: CI = [4.7640, 6.2135], $RR \to HR$: $\text{CI}_{95\%}$ = [13.3483, 17.1910]).

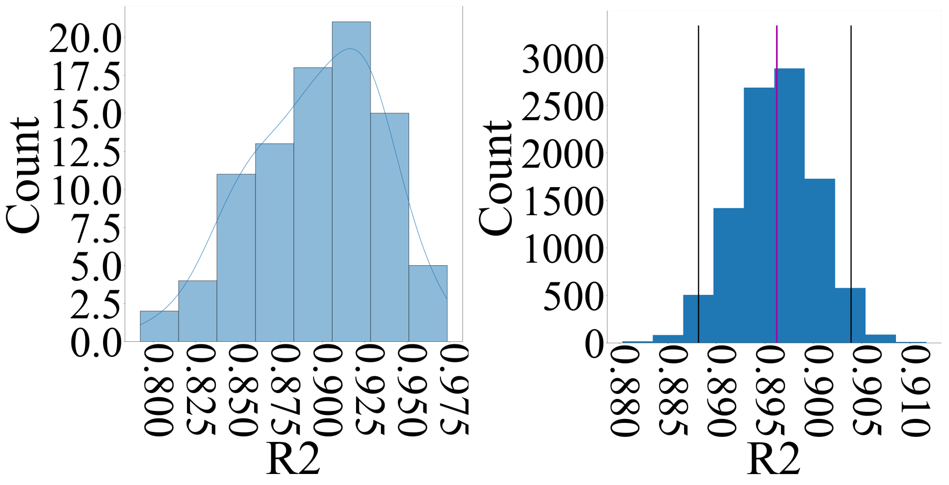

(a) (b)

**Fig. 3**: (a) individuals' AR accuracy $R^2$ distribution (b) Bootstrap (10,000 repetitions) estimate of $\mu_{R^2}$ at 95% confidence interval (CI) (M = 0.8962, Mdn = 0.8963, SD = 0.0041, CI = [0.8882, 0.9039]).

#### 6 AR Accuracy Reflects the Covariation Between HR State-Space's Present vs. Its Past Dimensions

We observed significant correlations between AR's $w_1 - w_2$ (Fig. 6a, r = -0.4610, p = 5.46e$^{-06}$) and $w_1 - w_3$ (Fig. 6b, r = -0.3880, p = 1.72$^{-04}$). Whereas AR accuracy $R^2$ followed the (inverse) covariation between $w_1$ (i.e., HR state-space most recent dimension) on the one hand, and $w_2$ and $w_3$ (i.e., HR state-space past history), on the other hand, such a relation was absent in $w_2 - w_3$ plane (i.e., within its history, Appendix 8).

Given the considerably stronger correlation within $w_2$ and $w_3$ plane (Appendix 8) than $w_1 - w_2$ (Figure 6a) and $w_1 - w_3$ (Figure 6b) planes, these results suggested that AR accuracy $R^2$ was not correlation-driven.

Figure 7 shows the covariation between AR's $w_1 - w_2$ with respect to HR state-space information dynamics $AIS$ and $I$. A comparison between this figure and Figure 6 verifies that AR accuracy $R^2$ followed HR state-space $AIS$ and $I$. It is worth noting that although this relation was also well-preserved in $w_1 - w_3$ plane (Appendix 8, Figure 8a for $AIS$ and Figure 8b for $I$), it was absent in $w_2 - w_3$ plane (Appendix 8, Figure 8c for $AIS$ and Figure 8d for $I$). We observed that such relations were absent in the case of other HR measures (SM2).

#### 7 $AIS$ and $I$ within $w_1 - w_3$ and $w_2 - w_3$ Planes

A comparison between Figures 8, 6, and 7 verifies that whereas AR accuracy $R^2$ followed HR state-space $AIS$ and $I$ within $w_1 - w_2$ (Figure 7a for $AIS$ and Figure 7b for $I$) and $w_1 - w_3$ (Figure 8a for $AIS$ and Figure 8b for $I$) planes, it was absent in $w_2 - w_3$ (Figure 8c for $AIS$ and Figure 8d for $I$) plane.

#### 8 $R^2$ within $w_2 - w_3$ Plane

We observed significant correlation between AR's $w_2 - w_3$ (Fig. 9, r = 0.7399, p = $1.20^{-16}$). However, AR's $R^2$ was not associated the covariation between $w_2$ and $w_3$ (i.e., within HR state-space history). Given the considerably stronger correlation

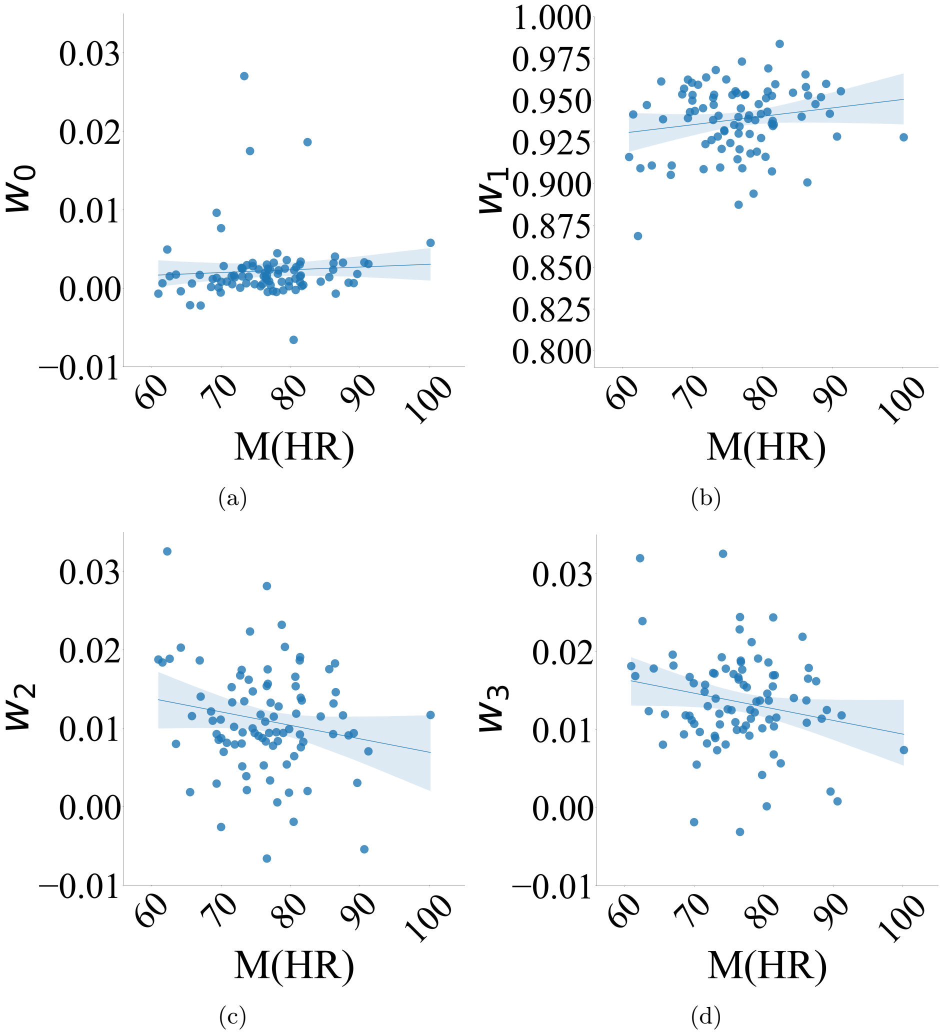

**Fig. 4**: Average HR versus AR parameters' (a) $W_0$ (b) $W_1$ (c) $W_2$ (d) $W_3$. We observed no correlation between $\mu_{HR}$ and AR parameters or its accuracy.

within $w_2$ and $w_3$ plane (Figure 9) than $w_1 - w_2$ (Figure 6a) and $w_1 - w_3$ (Figure 6b) planes, these results suggested that AR accuracy $R^2$ was not correlation-driven.

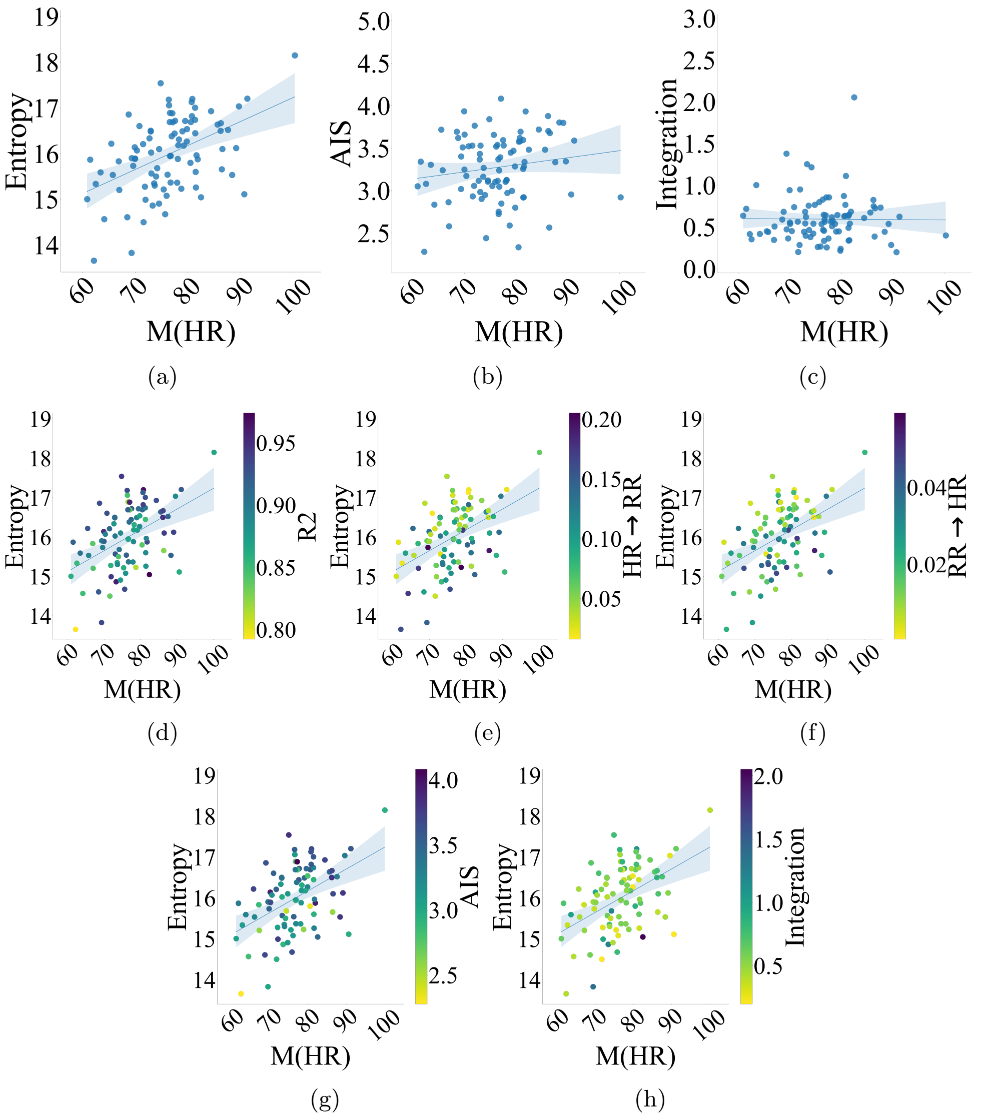

**Fig. 5**: Average HR versus HR (a) entropy ($H$) (b) $AIS$ (c) integration ($I$). Whereas we observed no correlation between $\mu_{HR}$ and $AIS$ or $I$, $\mu_{HR}$ was significantly correlated with $H$. However, this correlation did not present any correspondence to (d) AR accuracy $R^2$ (e) $HR \rightarrow RR$ (f) $RR \rightarrow HR$ (g) $AIS$ or (h) $I$.

### 9 $\boldsymbol{HR \rightarrow RR}$, $\boldsymbol{AIS}$, $\boldsymbol{I}$, and $\boldsymbol{R^2}$

Whereas $HR \rightarrow RR$ did not show any correlation with $R^2$ (Figure 10a, r = -0.1623, p = 1.29e$^{-01}$) and HR state-space $I$ (Figure 10b, r = -0.0157, p = 8.87e$^{-01}$), its correlation with $AIS$ did not pass the Bonferroni-correction (Figure 10c, r = -0.2816, p = 7.51e$^{-03}$). Additionally, this figure confirms that, unlike the case of $RR \rightarrow HR$ (see main manuscript), AR accuracy $R^2$ as well as HR state-space $AIS$ and $I$ did not bear any correspondence with $HR \rightarrow RR$.

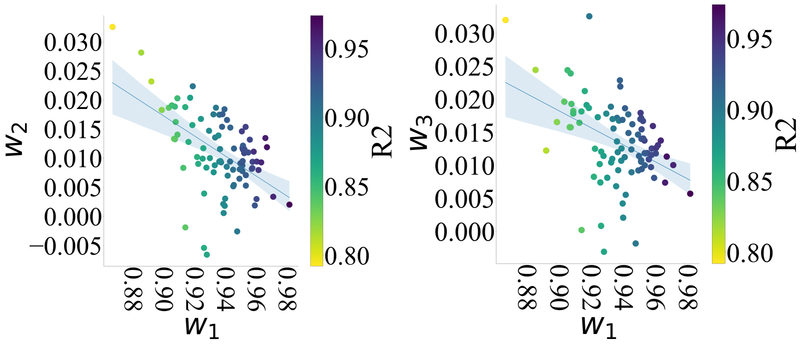

(a) (b)

**Fig. 6**: AR accuracy $R^2$ within AR's (a) $w_1 - w_2$ (b) $w_1 - w_3$ planes. Each of paired AR parameters were significantly correlated within their respective plane. This figure indicates that AR's $R^2$ was captured by $w_1$'s (inverse) covariations with $w_2$ and $w_3$ than evidently more strongly correlated $w_2$ and $w_3$ (Appendix 8).

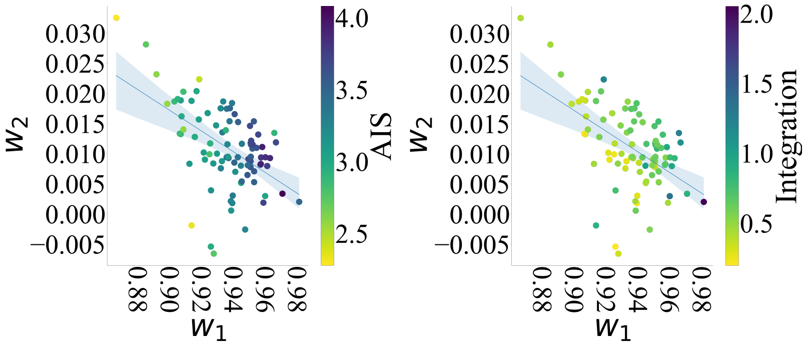

(a) (b)

**Fig. 7**: HR state-space information dynamics within AR's $w_1 - w_2$ plane (a) $AIS$ (b) $I$. Correspondence between AR accuracy $R^2$ (Figure 6) and HR state-space $AIS$ and $I$ within $w_1 - w_2$ is evident (See Appendix 7 for $AIS$ and $I$ within AR's $w_1 - w_3$ and $w_2 - w_3$ planes.

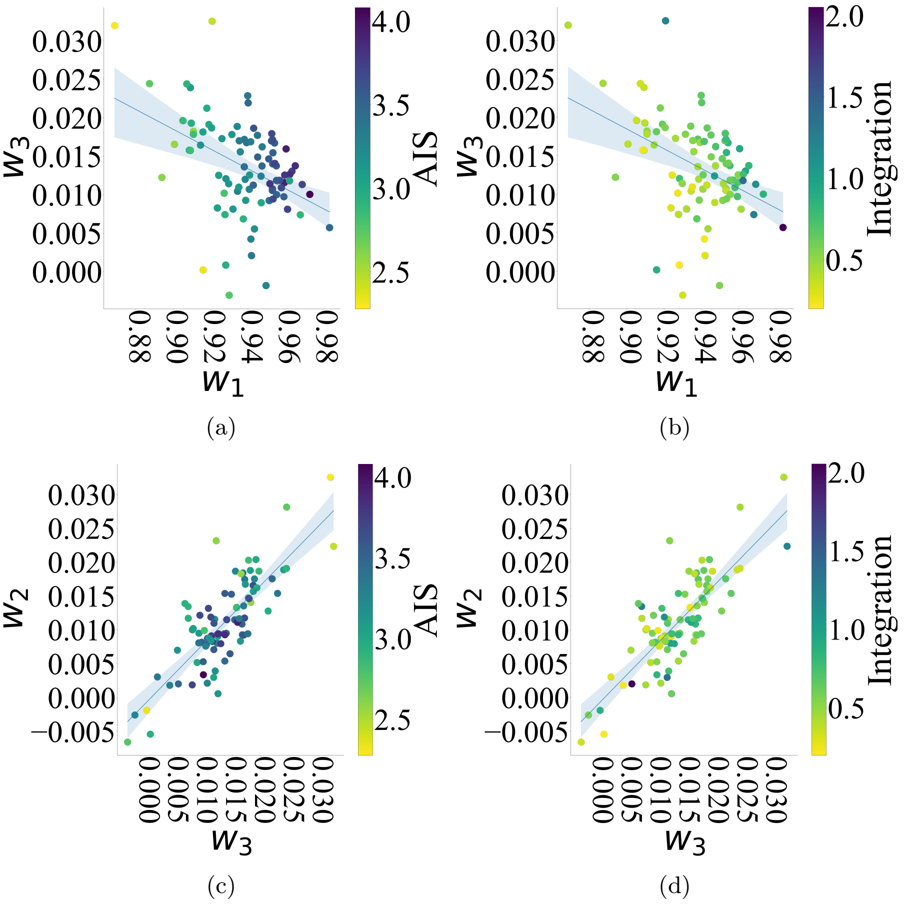

**Fig. 8**: $AIS$ and $I$ within (a-b) $w_1 - w_3$ (c-d) $w_2 - w_3$ planes. AR accuracy $R^2$ follows $AIS$ and $I$ within $w_1 - w_2$ (Figure 7) and $w_1 - w_3$ planes. Absence of such a correspondence in $w_2 - w_3$ plane is evident in this figure.

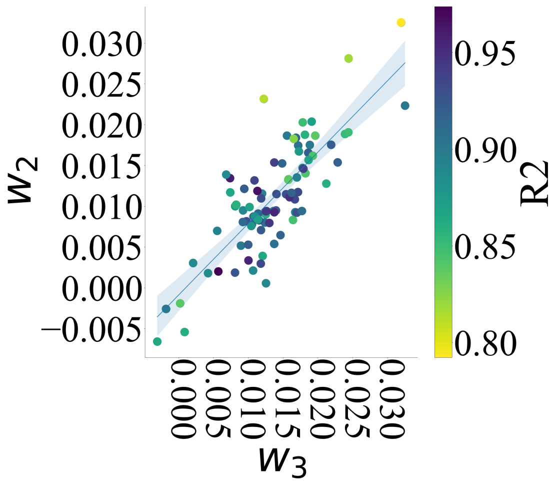

**Fig. 9**: AR accuracy $R^2$ within its $w_2 - w_3$ plane. Although $w_2$ and $w_3$ were evidently strongly correlated, their covariation did not relate with the AR's accuracy $R^2$.

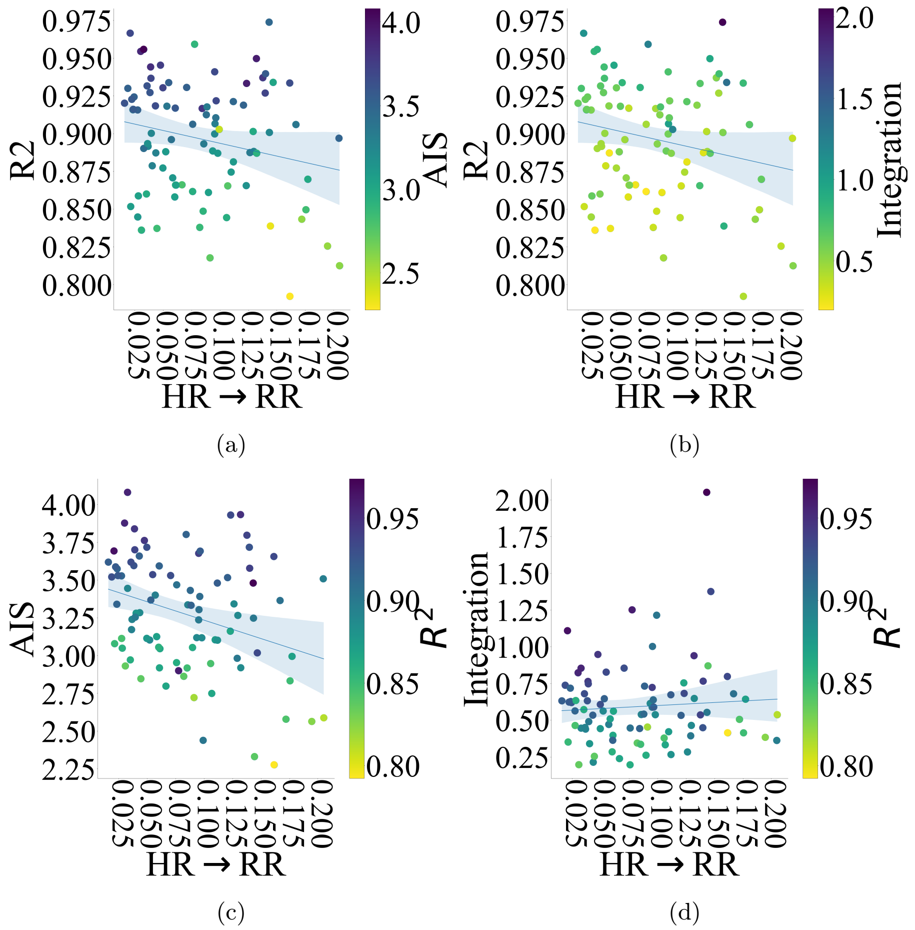

**Fig. 10**: (a) $AIS$ within $HR \to RR - R^2$ plane (b) $I$ within $HR \to RR - R^2$ plane (c) $R^2$ within $HR \to RR - AIS$ plane (d) $R^2$ within $HR \to RR - I$ plane. Unlike $RR \to HR$ (Figure **??**), no discernible correspondence between $HR \to RR$, on the one hand, and $R^2$, $AIS$, or $I$, on the other hand, was present.

#### 10 Effect of Weekly Alcohol Consumption on $RR \to HR$ During Wake

Alcohol consumption had significant effects on Wake $RR \to HR$. Compared to “Non-Drinkers”, “Drinkers” had significantly higher Wake $RR \to HR$ (Figure 11, test-statistics = -0.0081, p = 3.90e$^{-02}$, g = -0.6458, Non-Drinkers: M = 0.0195, Mdn = 0.0192, SD = 0.0119, CI$_{95\%}$ = [0.0099, 0.0344], Drinkers: M = 0.0297, Mdn = 0.0272, SD = 0.0180, CI$_{95\%}$ = [0.0141, 0.0509]).

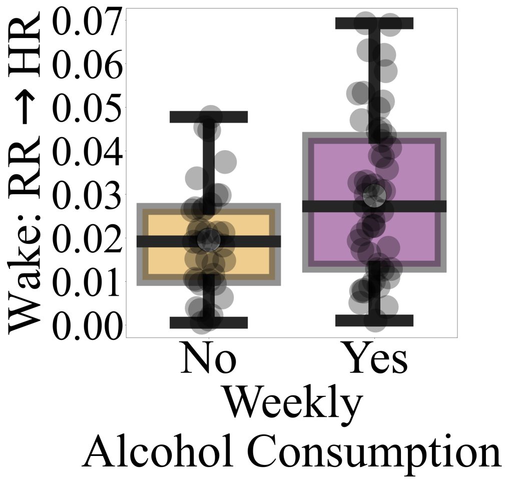

**Fig. 11**: Effect of weekly alcohol consumption on Wake $RR \to HR$ based on non-parametric bootstrap (10,000 repetitions) permutation test of difference in two groups' Mdn. Compared to “Non-Drinkers,” “Drinkers” had higher wake $RR \to HR$.

Alcohol consumption on its own (i.e., in the absence of smoking and exercise habits) had a significant effect on Wake $RR \to HR$. Precisely, “Drinkers” had a significantly higher Wake $RR \to HR$ (Figure 12, test-statistics = -0.026, p = 1.40e$^{-03}$, g = -1.4363, Non-Drinkers: M = 0.01, Mdn = 0.01, SD = 0.01, CI$_{95\%}$ = [0.0068, 0.0247], Drinkers: M = 0.04, Mdn = 0.04, SD = 0.02, CI$_{95\%}$ = [0.0207, 0.0621]).

#### 11 Alcohol Consumption and Exercise

We observed a significant effect of exercise (in the absence of smoking and alcohol consumption habits) on $RR \to HR$. Specifically, we observed that those who exercised showed a significantly higher Wake $RR \to HR$ (Figure 13, test-statistics = 0.0088, p = 1.12e$^{-02}$, g = 1.0857, Exercise: M = 0.02, Mdn = 0.02, SD = 0.01, CI$_{95\%}$ = [0.0167, 0.0401], No Exercise: M = 0.01, Mdn = 0.01, SD = 0.01, CI$_{95\%}$ = [0.0068, 0.0247]).

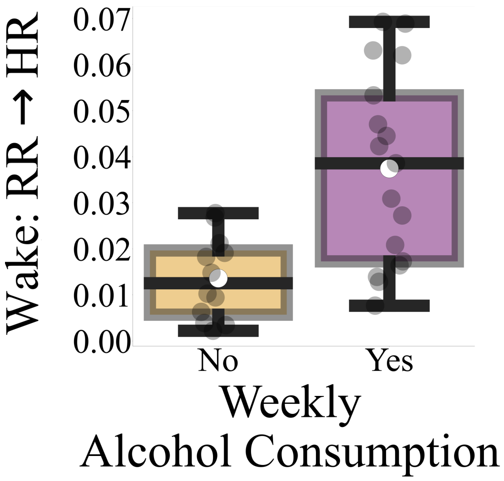

**Fig. 12**: Effect of alcohol consumption habit in the absence of smoking and exercise habits on Wake $RR \rightarrow HR$, based on non-parametric bootstrap (10,000 repetitions) permutation test of difference in two groups' Mdn.

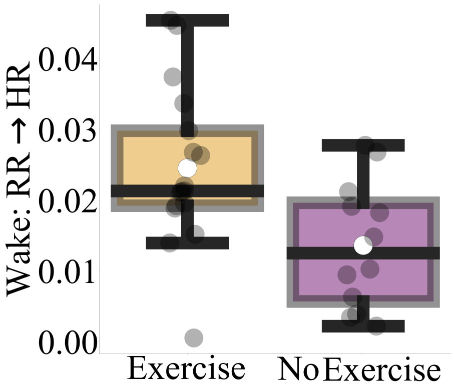

**Fig. 13**: Effect of exercise habit in the absence of smoking and alcohol consumption habits on Wake $RR \rightarrow HR$, based on non-parametric bootstrap (10,000 repetitions) permutation test of difference in two groups' Mdn.

##### 11.1 Individuals Who Exercised But Differed in Their Alcohol Consumption Habit

Once alcohol consumption was taken into account, the observed difference in $RR \rightarrow HR$ due to the exercise habit disappeared in both overall $RR \rightarrow HR$ (Figure 14a, test-statistics = -0.001, p = $8.45^{-01}$, g = 0.1005, Exercise but Not Drinking: M = 0.02, Mdn = 0.02, SD = 0.01, $CI_{95\%}$ = [0.0142, 0.0346], Exercise and Drinking: M = 0.02, Mdn = 0.02, SD = 0.01, $CI_{95\%}$ = [0.0104, 0.0365]) and Wake $RR \rightarrow HR$ (Figure 14b, test-statistics = -0.0039, p = 6.18e$^{-01}$, g = 0.0236, Exercise but Not Drinking: M =

0.02, Mdn = 0.02, SD = 0.01, $\text{CI}_{95\%}$ = [0.0167, 0.0401], Exercise and Drinking: M = 0.02, Mdn= 0.03, SD = 0.01, $\text{CI}_{95\%}$ = [0.0122, 0.043]).

Figure 15 verifies that (i.e., red arrows) the increase in $RR \rightarrow HR$ by exercise did not exceed the $RR \rightarrow HR$ of those who did not consume alcohol.

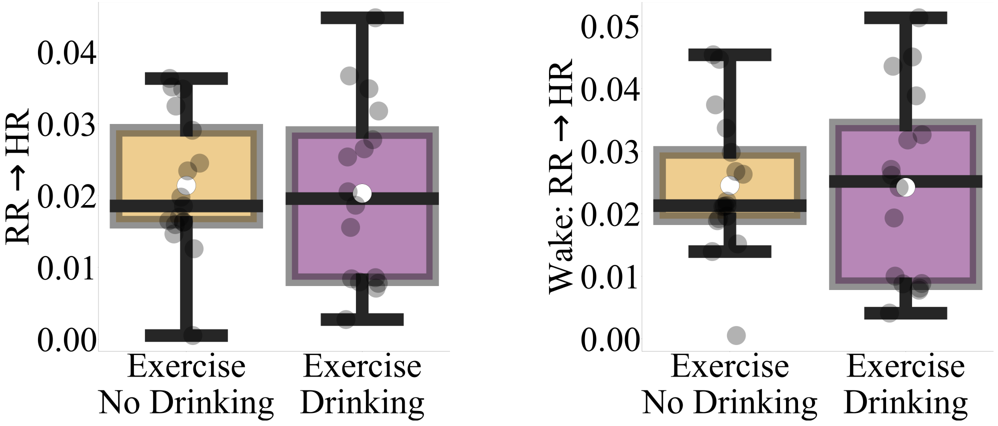

(a) (b)

**Fig. 14**: Alcohol consumption and exercise. (a) overall $RR \rightarrow HR$ (b) wake $RR \rightarrow HR$, based on non-parametric bootstrap (10,000 repetitions) permutation test of difference in two groups' Mdn.

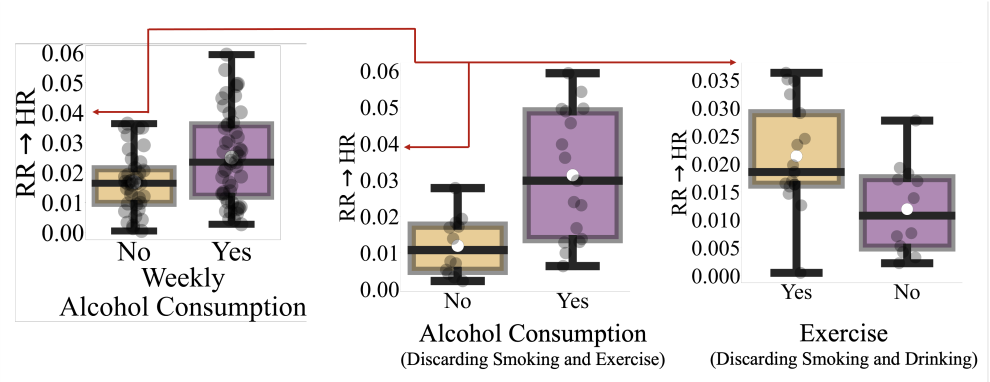

(a) (b) (c)

**Fig. 15**: Change in $RR \rightarrow HR$ with respect to (a) weekly alcohol consumption (b) weekly alcohol consumption after discarding smoking and exercise habits (c) exercise after discarding smoking and alcohol consumption habits. The arrows mark the upper bound for $RR \rightarrow HR$ as observed in our data. It is evident in this figure that increase in $RR \rightarrow HR$ due to exercise did not exceed the upper bound of $RR \rightarrow HR$ of those who did not consume alcohol.

##### 11.2 Individuals Who Consumed Alcohol But Differed in Their Exercise Habit

We observed that among the individuals who consumed alcohol, those exercised had lower $RR \rightarrow HR$ than those who did not exercise. However, the difference between these two sub-samples was non-significant in terms of both, overall $RR \rightarrow HR$ (Figure 16a, test-statistics = 0.0102, p = $2.33^{-01}$, g = 0.7337, Drinking & No-Exercise: M = 0.03, Mdn = 0.03, SD = 0.02, $CI_{95\%}$ = [0.0168, 0.0518], Drinking & Exercise: M = 0.02, Mdn = 0.02, SD = 0.01, $CI_{95\%}$ = [0.0104, 0.0365]) as well as wake $RR \rightarrow HR$ (Figure 16b, test-statistics = 0.0134, p = $1.93^{-01}$, g = 0.7378, M = 0.04, Mdn = 0.04, SD = 0.02, $CI_{95\%}$ = [0.0207, 0.0621], Drinking & Exercise: M = 0.02, Mdn = 0.03, SD = 0.01, $CI_{95\%}$ = [0.0122, 0.043]).

#### 12 Factor Analysis of the Alcohol – Exercise Interplay: Wake $RR \rightarrow HR$

Two-factor ANOVA showed significant effect of alcohol (F = 7.6678, p = $1.13e^{-02}$ (FDR corrected), $\eta^2$ = 0.1322) and alcohol × exercise interaction (F = 9.5010, p = $9.42e^{-03}$ (FDR corrected), $\eta^2$ = 0.1638). On the other hand, it revealed that exercise had no significant effect (F = 0.2873, p = 0.5940, $\eta^2$ = 0.0050).

Follow-up posthoc two-sample Welch test (equivalent of two-sample t-test for unequal variances) indicated significant differences between two scenarios (Figure 17) (1) those who exercised but did not consume alcohol versus those who neither exercised nor consumed alcohol (t = 2.7824, p = $2.92e^{-02}$ (FDR Corrected), g = 1.0857) and (2) those who did not exercise but consumed alcohol versus those who neither exercised nor consumed alcohol (t = 3.6903, p = $5.99e^{-03}$ (FDR corrected), g = 1.4363).

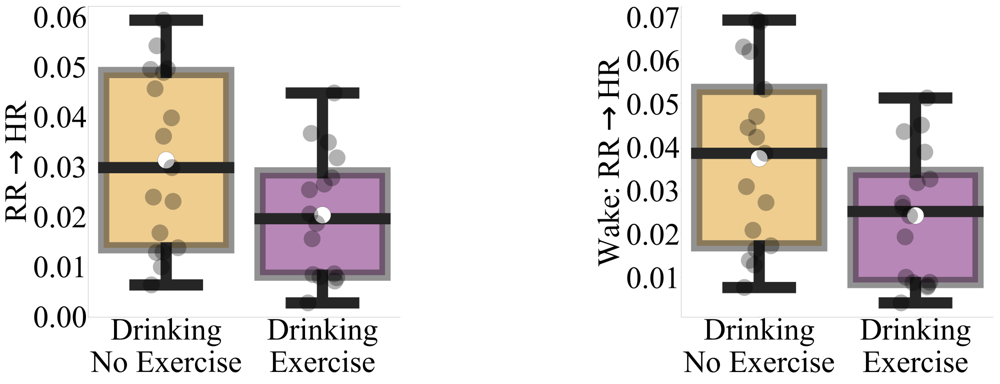

(a) (b)

**Fig. 16**: Individuals who consumed alcohol but differed in their exercise habits: those who did not exercise versus those who exercised but otherwise none of them smoked (or quit smoking) (a) overall $RR \rightarrow HR$ (b) wake $RR \rightarrow HR$, based on non-parametric bootstrap (10,000 repetitions) permutation test of difference in two groups' Mdn.

On the other hand, we found no significant differences between those who exercised as well as consumed alcohol versus, those who exercised but did not consume alcohol (F = -0.0656, (FDR Corrected) p = 0.9481, g = -0.0236), those who did not exercise but consumed alcohol (F = -2.0536, (FDR-Corrected) p = 5.82e$^{-02}$, g = -0.7378), or those who neither exercised nor consumed alcohol (F = 2.1285, (FDR-Corrected) p = 5.82e$^{-02}$, g = 0.8414). However, it was interesting to note the observed strong effect in the case of this latter scenario. Last, we did not find any significant difference between those who exercised but did not consume alcohol and those who did not exercise but consumed alcohol (F = -2.2356, (FDR-Corrected) p = 5.82e$^{-02}$), g = -0.7904).

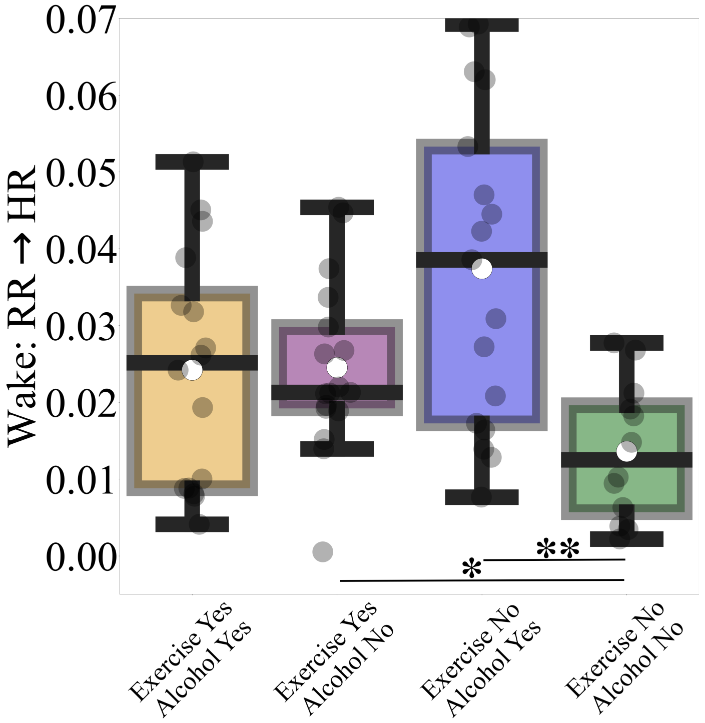

**Fig. 17**: ANOVA analysis of alcohol – exercise interplay during Wake period. Whereas alcohol consumption and its interaction with exercise were associated with significant effects on Wake $RR \rightarrow HR$, effect of exercise was non-significant. Additionally, there were significant differences in Wake $RR \rightarrow HR$ between (1) those who exercised but did not consume alcohol versus those who neither exercised nor consumed alcohol and (2) those who did not exercise but consumed alcohol versus those who neither exercised nor consumed alcohol (*: p < 0.05, **: p < 0.01).

### 13 Non-Parametric Analysis of Alcohol – Exercise Interplay Interplay: $\boldsymbol{RR \rightarrow HR}$

#### 13.1 Overall $\boldsymbol{RR \rightarrow HR}$

Kruskal-Wallis indicated a significant difference in Overall $RR \rightarrow HR$ between the four groups (see Materials and Methods, main manuscript) i.e., those who exercised but did not consume alcohol, those who neither exercised nor consumed alcohol, those who did not exercise but consumed alcohol, and those who exercised as well as consumed alcohol (H = 10.7001, p = 1.35e$^{-02}$ (FDR corrected), $\eta^2$ = 0.1328).

Posthoc Wilcoxon rank-sum test indicated a significant difference in Overall $RR \rightarrow HR$ between two scenarios those who did not exercise but consumed alcohol versus those who neither exercised nor consumed alcohol (W = 2.7897, p = 5.30e$^{-03}$ (FDR corrected), $\eta^2$ = 0.1328).

On the other hand, we found no significant differences between those who exercised as well as consumed alcohol versus, those who exercised but did not consume alcohol (W = -0.3602, (FDR-Corrected) p = 7.19e$^{-01}$, g = 0.0473), those who did not exercise but consumed alcohol (W = -1.8371, (FDR-Corrected) p = 9.27e$^{-02}$, g = 0.2412), or those who neither exercised nor consumed alcohol (W = 2.0426, (FDR-Corrected) p = 7.19e$^{-02}$, g = 0.2682). Last, we did not find any significant difference between those who exercised but did not consume alcohol and those who did not exercise but consumed alcohol (W = -1.3605, (FDR-Corrected) p = 2.03e$^{-01}$, g = 0.1786) or those who neither exercised nor consumed alcohol (W = 2.214, (FDR-Corrected) p = 6.26e$^{-02}$, g = 0.2907).

#### 13.2 Wake $\boldsymbol{RR \rightarrow HR}$

These differences between the four groups were also present in the case of Wake $RR \rightarrow HR$. Precisely, Kruskal-Wallis showed a significant difference between these four groups (H = 11.6341, p = 8.75e$^{-03}$ (FDR corrected), $\eta^2$ = 0.1489).

Posthoc Wilcoxon rank-sum test further indicated significant differences between two scenarios (1) those who exercised but did not consume alcohol versus those who neither exercised nor consumed alcohol (W = 2.4797, p = 3.07e$^{-02}$ (FDR Corrected), $\eta^2$ = 0.3256) and (2) those who did not exercise but consumed alcohol versus those who neither exercised nor consumed alcohol (W = 3.0111, p = 1.82e$^{-02}$ (FDR corrected), g = 0.3954).

On the other hand, we found no significant differences between those who exercised as well as consumed alcohol versus, those who exercised but did not consume alcohol (W = -0.1441, (FDR-Corrected) p = 8.85e$^{-01}$), g = 0.0189), those who did not exercise but consumed alcohol (W = -1.8011, (FDR-Corrected) p = 1.00e$^{-01}$, g = 0.2365), or those who neither exercised nor consumed alcohol (W = 1.857 (FDR-Corrected), p = 1.00e$^{-01}$, g = 0.2438). Last, we did not find any significant difference between those who exercised but did not consume alcohol and those who did not exercise but consumed alcohol (W = -1.4639, (FDR-Corrected) p = 1.67e$^{-01}$, g = 0.1922).

### 14 Bootstrap Non-Significant Results

#### 14.1 Gender

Female and male participants showed no significant differences with respect to their M(HR) (Figure 18a, test-statistics = -3.142, p = 5.26e$^{-02}$, g = -0.468, Females: M = 71.5107, Mdn = 71.3301, SD = 6.9019, CI$_{95\%}$ = [65.8831, 80.0096], Males: M = 74.8817, Mdn = 74.4721, SD = 7.4279, CI$_{95\%}$ = [68.6607, 84.2455]), M(RR) (Figure 18b, test-statistics = 0.3446, p = 3.40e$^{-01}$, g = 0.0745, Females: M = 17.026, Mdn = 17.111, SD = 1.4123, CI$_{95\%}$ = [15.876, 18.8172], Males: M = 16.9097, Mdn = 16.7664, SD = 1.6693, CI$_{95\%}$ = [15.495, 18.9898]), AR accuracy $R^2$ ((Figure 18c, test-statistics = 0.0028, p = 8.02e$^{-01}$, g = 0.0625, Females: M = 0.8975, Mdn = 0.9028, SD = 0.0372, CI$_{95\%}$ = [0.8661, 0.9428], Males: M = 0.8952, Mdn = 0.9, SD = 0.0387, CI$_{95\%}$ = [0.8624, 0.943]), $H(HR)$ (Figure 18d, test-statistics = -0.1937, p = 4.90e$^{-01}$, g = -0.1136, Females: M = 15.9358, Mdn = 15.8896, SD = 0.8318, CI$_{95\%}$ = [15.2315, 16.974], Males: M = 16.0278, Mdn = 16.0832, SD = 0.7917, CI$_{95\%}$ = [15.3738, 17.067]), $AIS(HR)$ (Figure 18e, test-statistics = 0.0348, p = 8.44e$^{-01}$, g = 0.0283, Females: M = 3.2781, Mdn = 3.304, SD = 0.3807, CI$_{95\%}$ = [2.966, 3.7577], Males: M = 3.2674, Mdn = 3.2692, SD = 0.3783, CI$_{95\%}$ = [2.945, 3.7326]), $HR \rightarrow RR$ (Figure 18f, test-statistics = -0.0087, p = 7.19e$^{-01}$, g = 0.0134, Females: M = 0.0849, Mdn = 0.0759, SD = 0.0543, CI$_{95\%}$ = [0.0389, 0.1508], Males: M=0.0842, Mdn = 0.0847, SD = 0.0448, CI$_{95\%}$ = [0.0457, 0.1376]), and $RR \rightarrow HR$ (Figure 18g, test-statistics = -0.0045, p = 1.48e$^{-01}$, g = -0.1754, Females: M = 0.0201, Mdn = 0.0167, SD = 0.0131, CI$_{95\%}$ = [0.0097, 0.0364], Males: M = 0.0225, Mdn = 0.0212, SD = 0.014, CI$_{95\%}$ = [0.0106, 0.0396]).

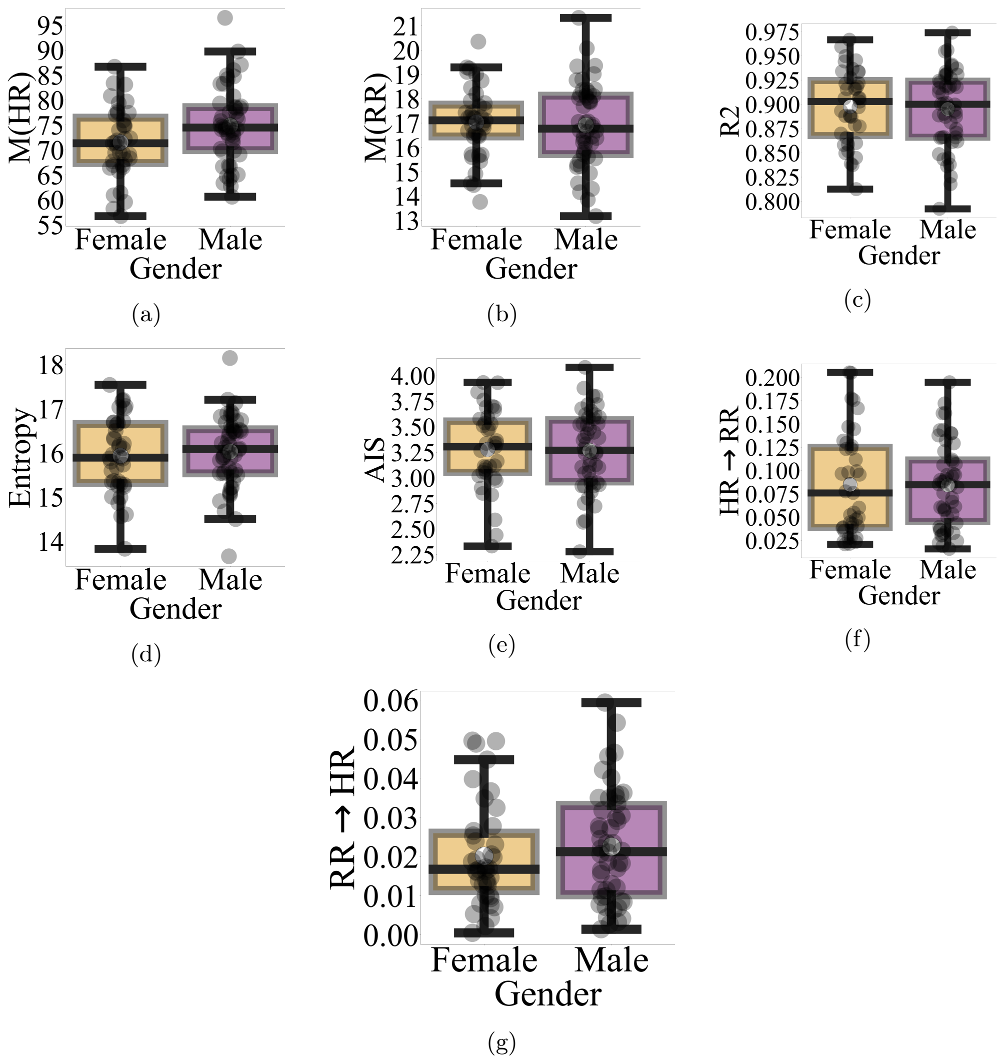

**Fig. 18**: Gender did not show any significant differences with respect to (a) M(HR) (b) M(RR) (c) AR accuracy $R^2$ (d) $H(HR)$ (e) $AIS(HR)$ (f) $HR \rightarrow RR$ (g) $RR \rightarrow HR$.

#### 14.2 Age

Considering the age as a factor, we observed no significant differences between participants with respect to their M(HR) (Figure 19a, test-statistics = 1.2369, p = 5.49e$^{-02}$, g = 0.0267, Females: M = 73.4867, Mdn = 74.1154, SD = 6.7563, CI$_{95\%}$ = [67.8969, 81.8503], Males: M = 73.289 Mdn = 72.8785 SD = 8.205 CI$_{95\%}$ = [66.5056, 84.05]),M(RR) (Figure 19b, test-statistics = -0.3667, p = 2.97e$^{-01}$, g = -0.2951, Females: M = 16.7709, Mdn = 16.8228, SD = 1.4863, CI$_{95\%}$ = [15.4798, 18.5199], Males: M = 17.2273, Mdn = 17.1894, SD = 1.6277, CI$_{95\%}$ = [15.9431, 19.3679]), AR accuracy $R^2$ ((Figure 19c, test-statistics = -0.0138, p = 2.86e$^{-01}$, g = -0.1485, Females: M = 0.8939, Mdn = 0.8924, SD = 0.0361, CI$_{95\%}$ = [0.8624, 0.9363], Males: M = 0.8995, Mdn = 0.9062, SD = 0.0404, CI$_{95\%}$ = [0.867, 0.9512]), $H(HR)$ (Figure 19d, test-statistics = 0.1602, p = 5.88e$^{-01}$, g = -0.0174, Females: M = 15.9816, Mdn = 16.0603, SD = 0.7571, CI$_{95\%}$ = [15.3476, 16.9268, Males: M = 15.9957, Mdn = 15.9001, SD = 0.8806, CI$_{95\%}$ = [15.2667, 17.171]), $I(HR)$ (Figure 19e, test-statistics = 0.0623, p = 1.90e$^{-01}$, g = 0.0119, Females: M = 0.5946, Mdn = 0.5678, SD = 0.2505, CI$_{95\%}$ = [0.3981, 0.9183], Males: M= 0.5913, Mdn = 0.5055, SD = 0.3132, CI$_{95\%}$ = [0.404, 1.1514]), $HR \to RR$ (Figure 19f, test-statistics = 0.0077, p = 7.70e$^{-01}$, g = -0.108, Females: M = 0.0823, Mdn = 0.0836, SD = 0.0473, CI$_{95\%}$ = [0.0417, 0.1379], Males: M = 0.0876, Mdn = 0.0759, SD = 0.0515, CI$_{95\%}$ = [0.0445, 0.1513]), and $RR \to HR$ (Figure 19g, test-statistics = 0.0002, p = 9.26e$^{-01}$, g = 0.1301, Females: M = 0.0222, Mdn = 0.0188, SD = 0.0142, CI$_{95\%}$ = [0.0103, 0.0394], Males: M = 0.0204, Mdn = 0.0185, SD = 0.0127, CI$_{95\%}$ = [0.0097, 0.0358]).

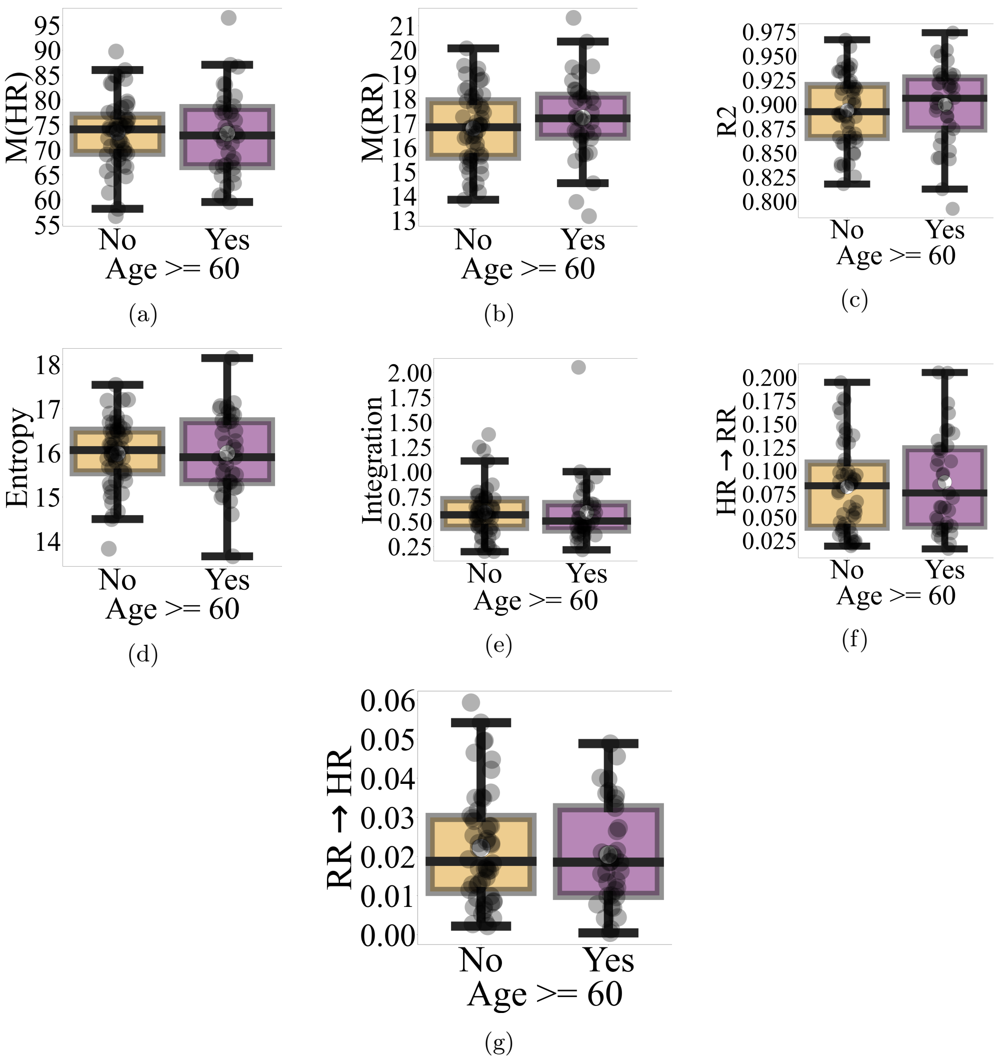

**Fig. 19**: Age did not show any significant differences with respect to (a) M(HR) (b) M(RR) (c) AR accuracy $R^2$ (d) $H(HR)$ (e) $I(HR)$ (f) $HR \rightarrow RR$ (g) $RR \rightarrow HR$.

#### 14.3 Alcohol Consumption

Alcohol consumption had no significant effects on participants' M(HR) (Figure 20a, test-statistics = 1.3524, p = 4.50e$^{-01}$, g = 0.1385, Females: M = 73.9897, Mdn = 74.7716, SD = 7.6199, $CI_{95\%}$ = [67.897, 84.164], Males: M = 72.9684, Mdn = 73.4191, SD = 7.1898, $CI_{95\%}$ = [66.9033, 81.696]), M(RR) (Figure 20b, test-statistics = -0.1526, p = 6.58e$^{-01}$, g = -0.0445, Females: M = 16.9209, Mdn = 16.88, SD = 1.8748, $CI_{95\%}$ = [15.3264, 19.2225], Males: M = 16.9903, Mdn = 17.0326, SD = 1.2814, $CI_{95\%}$ = [15.9229, 18.6037]), AR accuracy $R^2$ ((Figure 20c, test-statistics = 0.0151, p = 2.52e$^{-01}$, g = 0.2557, Females: M = 0.9017, Mdn = 0.9083, SD = 0.0414, $CI_{95\%}$ = [0.8678, 0.9545], Males: M = 0.8921, Mdn = 0.8932, SD = 0.0348, $CI_{95\%}$ = [0.8617, 0.9332]), $AIS(HR)$ (Figure 20d, test-statistics = 0.1185, p = 2.65e$^{-01}$, g = 0.1372, Females: M = 3.3018, Mdn = 3.3661, SD = 0.3896, $CI_{95\%}$ = [2.9871, 3.8077], Males: M = 3.2499, Mdn = 3.2475, SD = 0.37, $CI_{95\%}$ = [2.9341, 3.6945]), $I(HR)$ (Figure 20e, test-statistics = 0.034, p = 5.53e$^{-01}$, g = 0.201, Females: M = 0.6251, Mdn = 0.5756, SD = 0.322, $CI_{95\%}$ = [0.4194, 1.1827], Males: M = 0.5695, Mdn = 0.5416, SD = 0.2378, $CI_{95\%}$ = [0.3829, 0.8892]), and $HR \rightarrow RR$ (Figure 20f, test-statistics = -0.025, p = 2.32e$^{-01}$, g = -0.0801, Females: M = 0.0823, Mdn = 0.0607, SD = 0.051, $CI_{95\%}$ = [0.039, 0.1441], Males: M = 0.0862, Mdn = 0.0857, SD = 0.0476, $CI_{95\%}$ = [0.0458, 0.1439]).

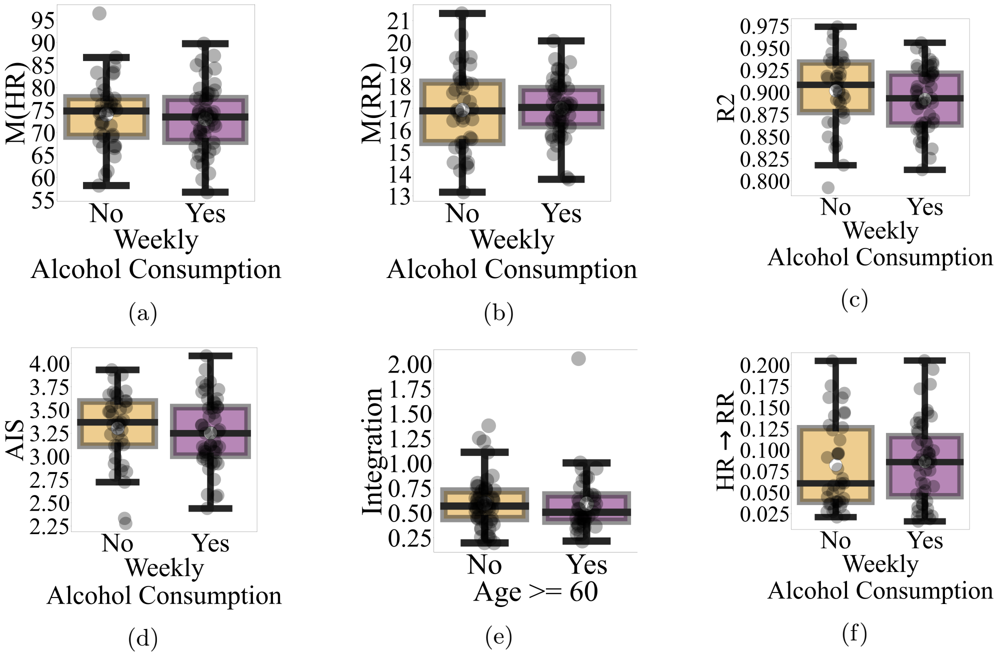

**Fig. 20**: Age did not show any significant differences with respect to (a) M(HR) (b) M(RR) (c) AR accuracy $R^2$ (d) $AIS(HR)$ (e) $I(HR)$ (f) $HR \rightarrow RR$.

#### 14.4 Exercise

Exercise had no significant effects on participants' M(HR) (Figure 21a, test-statistics = -0.527, p = 8.82e$^{-01}$, g = -0.212, Females: M = 72.5288, Mdn = 73.4191, SD = 7.3167, CI$_{95\%}$ = [66.4981, 81.4799], Males: M = 74.0876, Mdn = 73.9461, SD = 7.3815, CI$_{95\%}$ = [67.9286, 83.5571]), M(RR) (Figure 21b, test-statistics = -0.1073, p = 7.12e$^{-01}$, g = -0.1555, Females: M = 16.8245, Mdn = 16.8902, SD = 1.524, CI$_{95\%}$ = [15.5752, 18.733], Males: M = 17.0669, Mdn = 16.9976, SD = 1.5846, CI$_{95\%}$ = [15.7264, 19.0305]), AR accuracy $R^2$ ((Figure 21c, test-statistics = 0.0063, p = 6.02e$^{-01}$, g = 0.0496, Females: M = 0.8973, Mdn = 0.9028, SD = 0.0396, CI$_{95\%}$ = [0.8635, 0.9443], Males: M = 0.8954, Mdn = 0.8965, SD = 0.0368, CI$_{95\%}$ = [0.8646, 0.9416]), $H(HR)$ ((Figure 21d, test-statistics = -0.0821, p = 9.34e$^{-01}$, g = 0.1148, Females: M = 16.0396, Mdn = 15.9632, SD = 0.7239, CI$_{95\%}$ = [15.4261, 16.9163], Males: M = 15.9467, Mdn = 16.0453, SD = 0.8704, CI$_{95\%}$ = [15.2251, 17.0546]), $AIS(HR)$ (Figure 21e, test-statistics = 0.0539, p = 7.53e$^{-01}$, g = 0.0388, Females: M = 3.2803, Mdn = 3.3361, SD = 0.4446, CI$_{95\%}$ = [2.9047, 3.813], Males: M = 3.2656, Mdn = 3.2822, SD = 0.3192, CI$_{95\%}$ = [2.9956, 3.6819]), $I(HR)$ (Figure 21f, test-statistics = 0.0671, p = 2.40e$^{-01}$, g = 0.1103, Females: M = 0.6105, Mdn = 0.588, SD = 0.2133, CI$_{95\%}$ = [0.44, 0.8892], Males: M= 0.5798, Mdn = 0.5209, SD = 0.3193, CI$_{95\%}$ = [0.3619, 1.0968]), $HR \rightarrow RR$ (Figure 21g, test-statistics = -0.0248, p = 2.69e$^{-01}$, g = -0.0539, Females: M = 0.083, Mdn = 0.0606, SD = 0.0545, CI$_{95\%}$ = [0.0376, 0.1479], Males: M = 0.0857, Mdn = 0.0853, SD = 0.0445, CI$_{95\%}$ = [0.0469, 0.1399]), and $RR \rightarrow HR$ (Figure 21h, test-statistics = -0.0038, p = 2.66e$^{-01}$, g = -0.1188, Females: M = 0.0206, Mdn = 0.0167, SD = 0.016, CI$_{95\%}$ = [0.0076, 0.04], Males: M = 0.0222, Mdn = 0.0206, SD = 0.0115, CI$_{95\%}$ = [0.0123, 0.0355]).

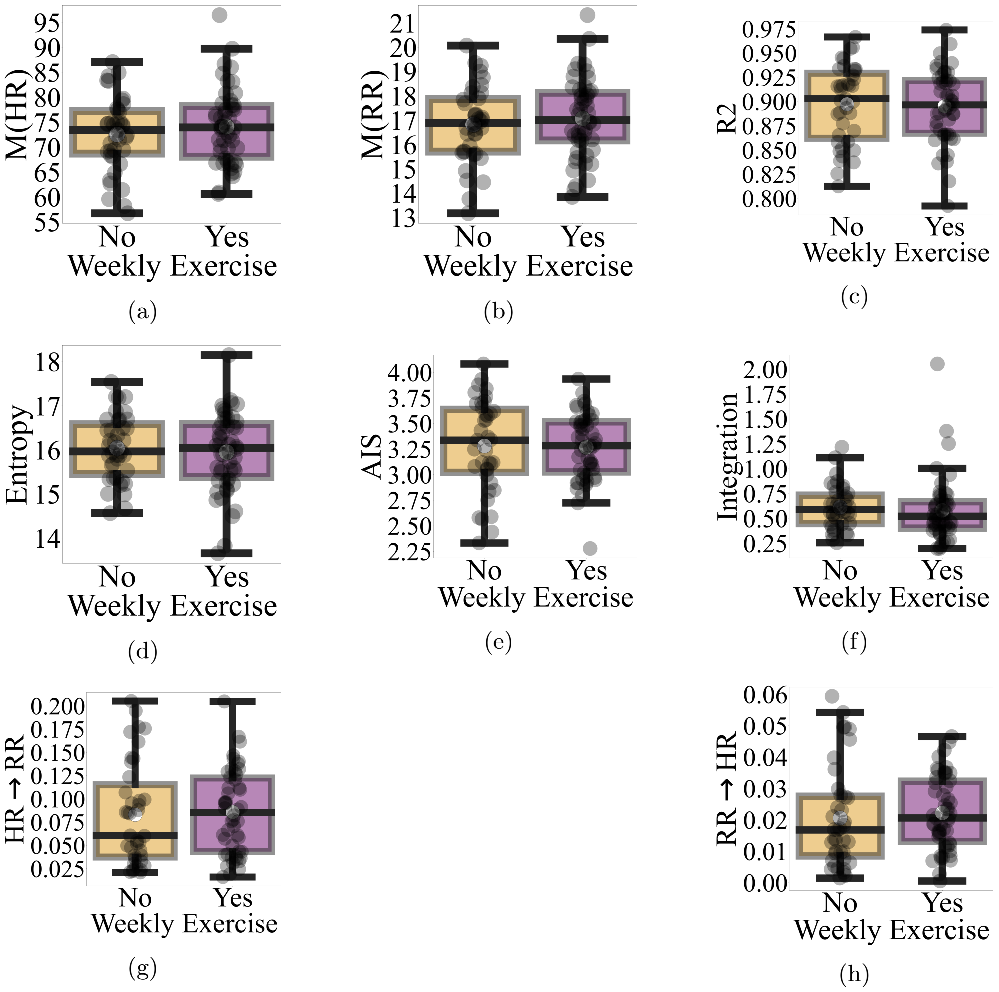

**Fig. 21**: Age did not show any significant differences with respect to (a) M(HR) (b) M(RR) (c) AR accuracy $R^2$ (d) $H(HR)$ (e) $AIS(HR)$ (f) $I(HR)$ (g) $HR \rightarrow RR$ (h) $RR \rightarrow HR$.

### 15 HR $\boldsymbol{AIS - RR \rightarrow HR}$ Plane, Average Daily Sleep, Walking/Standing, and Sitting/Leaning

Within HR $AIS - RR \rightarrow HR$ plane, there was no discernible distribution associated with the participants' average daily sleeping, movement, or resting hours.

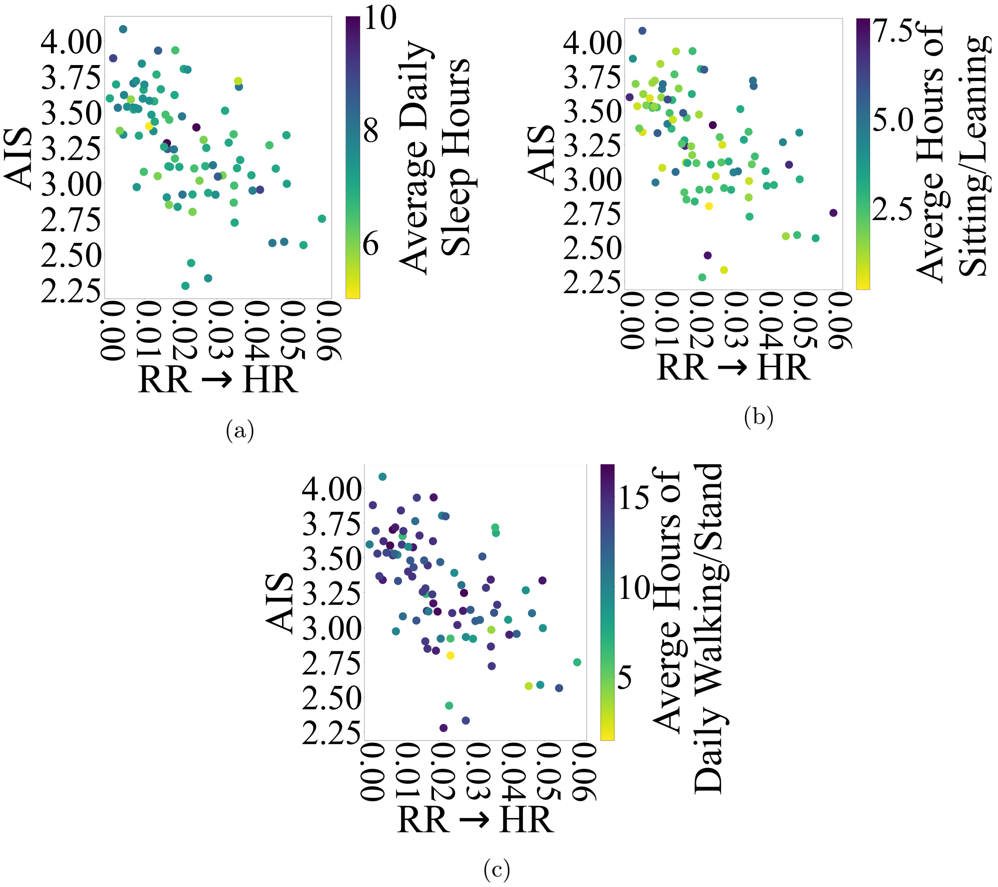

**Fig. 22**: $AIS - RR \rightarrow HR$ plane with respect to participants' average daily hours spent on (a) sleeping (b) sitting/leaning (c) standing/walking.
