## Supplementary Materials 2 for "Information Dynamics of the Heart and Respiration Rates: a Novel Venue for Digital Phenotyping in Humans"

### Supplementary Materials 2 (SM2):
Respiratory Modulation of the Heart Rate: a Potential Marker of the Effect of Alcohol on Cardiorespiratory Function in Human

Soheil Keshmiri[1*], Sutashu Tomonaga[2], Haruo Mizutani[3], Kenji Doya[2]

[1]Optical Neuroimaging Unit, Okinawa Institute of Science and Technology, Okinawa, Japan.
[2]Neural Computation Unit (NCU), Okinawa Institute of Science and Technology, Okinawa, Japan.
[3]Suntory Global Innovation Center Limited (SGIC), Suntory, Kyoto, Japan.

Contributing authors:;
;

**E Exercise** **19**

**F BMI- and HR-Related Gender Differences** **19**

**G Effect of Weekly Alcohol Consumption on BMI and HR (12:00 – 18:00 PM)** **19**

**H Relation Between Smoking And Participants' Age, BMI, Early Morning (6:00 – 7:00 AM) HR and RR, Overall and Wake RR** **20**

### Appendix A HR and RR Correlation

We observed that overall, the participants' HR and RR were not correlated (Figure A1a, r = 0.1999, p = $6.04e^{-02}$). Whereas this observation was also true for their HR and RR during sleep (Figure A1b, r = 0.1682, p = $1.15e^{-01}$) their wake HR and RR showed a correlation that did not pass the Bonferroni threshold (i.e., p = $6.0e^{-04}$)(Figure A1c, r = 0.3254, p = $1.86e^{-03}$).

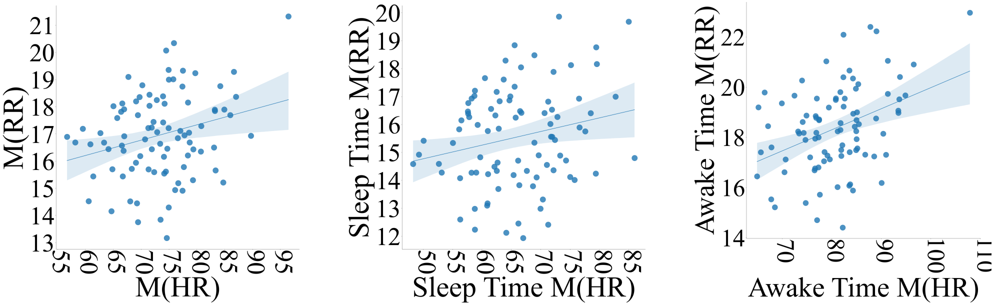

(a) (b) (c)

**Fig. A1**: Correlation between HR and RR (a) Overall (b) Sleep (c) Wake.

Figures A2a and A2b show the HR — RR scatter plots for females (Spearman's rank correlation coefficient M = 0.5713, SD = 0.1755, Mdn = 0.6052, $CI_{95\%}$ = [0.2228, 0.8078], Min = 0.0927, Max = 0.8281) and males (Spearman's rank correlation coefficient M = 0.5573, SD = 0.1312, Mdn = 0.5522, $CI_{95\%}$ = [0.2676, 0.7503], Min = 0.2344, Max = 0.7694) participants. In these subplots, data points are color-coded as per their respective time-of-day (in an hourly basis). Comparing the females' and males' correlations, their descriptive statistics did not show any tangible difference.

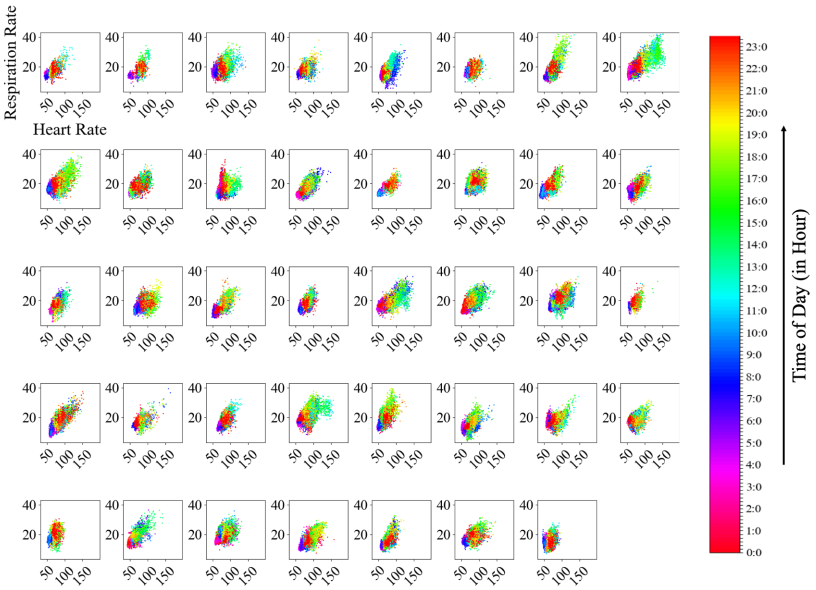

(a)

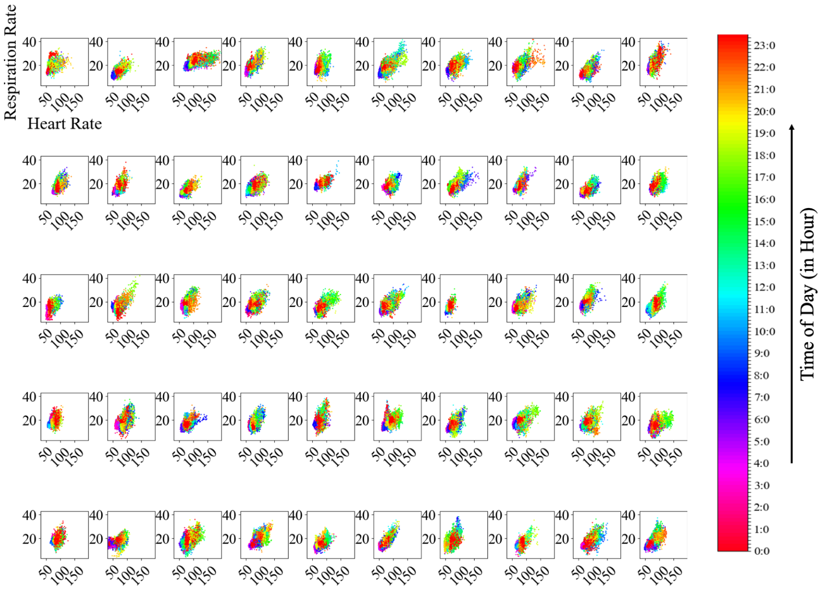

(b)

**Fig. A2**: HR — RR correlation of (a) females (b) males participants. Data points correspond to 1-minute non-overlapping moving average on HR and RR time series. They are color-coded as per time-of-day (in an hourly basis).

### Appendix B HR and RR Time Series

#### B.1 1 Day of Experiment

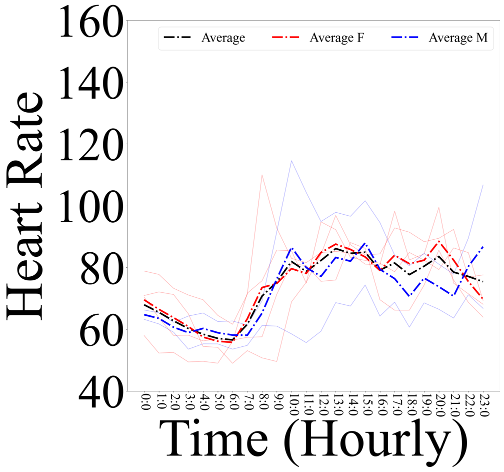

(a) HR

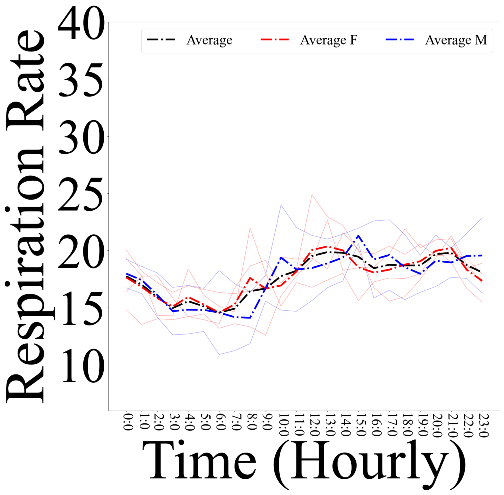

(b) RR

**Fig. B3**: (a) HR (b) RR time series of subset of participants who had only 1 day of experiment available (total of 6 participants). In these subplots, HR and RR time series, per participant, are averaged in a 1-hour non-overlapping interval. Female and male participants are presented in solid red and blue lines. Dashed red and blue lines are grand average HR and RR for female and male participants. Dashed black lines represent grand average of both female and male participants

#### B.2 2 Days of Experiment

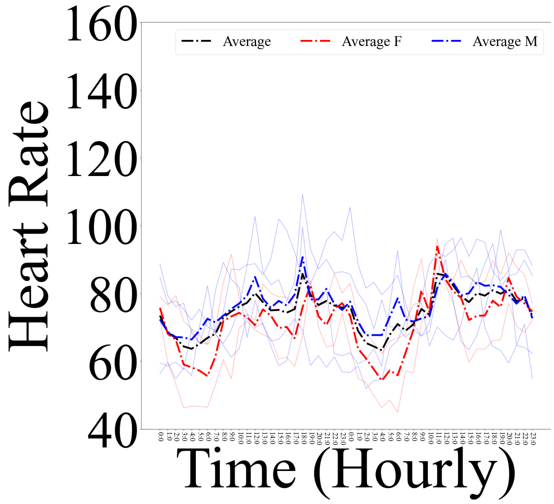

(a) HR

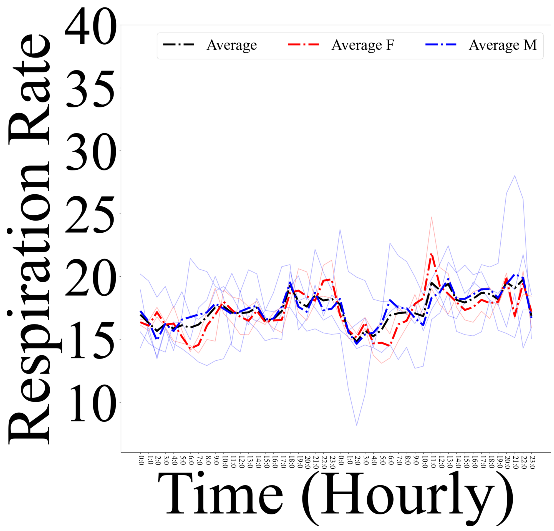

(b) RR

**Fig. B4**: (a) HR (b) RR time series of subset of participants who had only 2 days of experiment available (total of 6 participants). In these subplots, HR and RR time series, per participant, are averaged in a 1-hour non-overlapping interval. Female and male participants are presented in solid red and blue lines. Dashed red and blue lines are grand average HR and RR for female and male participants. Dashed black lines represent grand average of both female and male participants

#### B.3 3 Days of Experiment

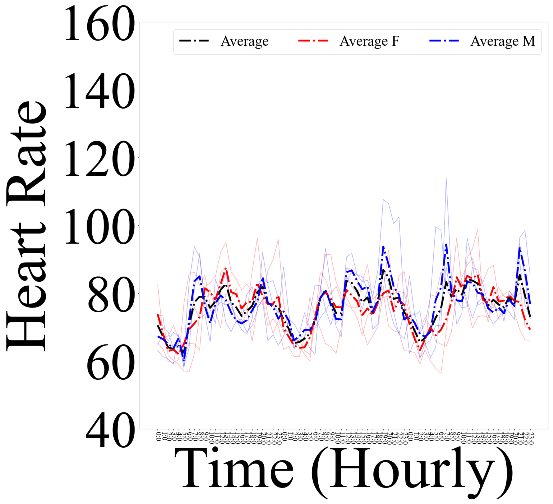

(a) HR

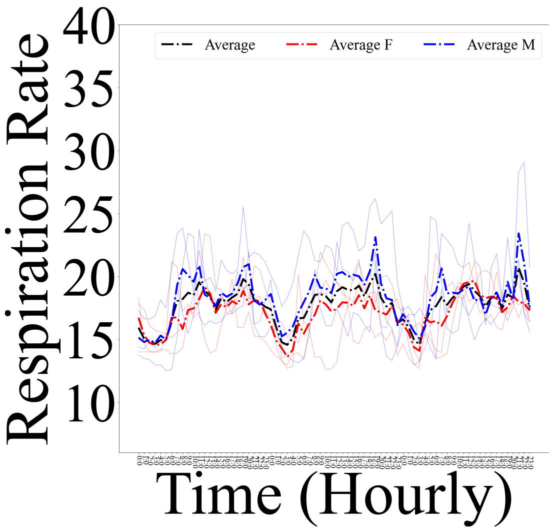

(b) RR

**Fig. B5**: (a) HR (b) RR time series of subset of participants who had only 3 days of experiment available (total of 6 participants). In these subplots, HR and RR time series, per participant, are averaged in a 1-hour non-overlapping interval. Female and male participants are presented in solid red and blue lines. Dashed red and blue lines are grand average HR and RR for female and male participants. Dashed black lines represent grand average of both female and male participants

##### B.4 4 Days of Experiment

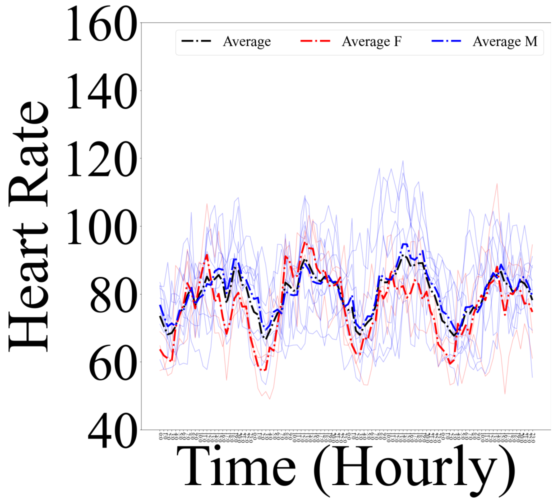

(a) HR

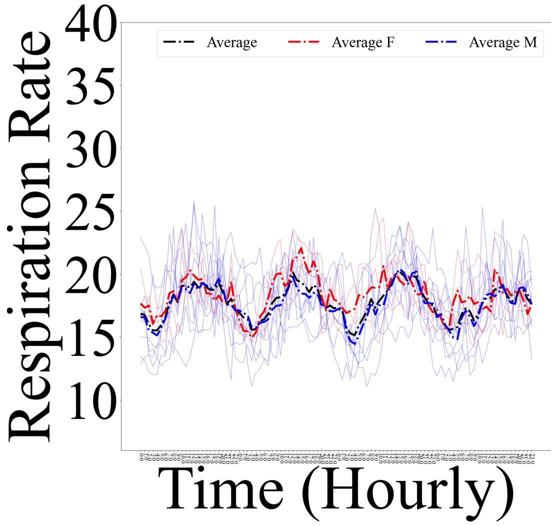

(b) RR

**Fig. B6**: (a) HR (b) RR time series of subset of participants who had only 4 days of their experiment available (total of 12 participants). In these subplots, HR and RR time series, per participant, are averaged in a 1-hour non-overlapping interval. Female and male participants are presented in solid red and blue lines. Dashed red and blue lines are grand average HR and RR for female and male participants. Dashed black lines represent grand average of both female and male participants

#### B.5 5 Days of Experiment

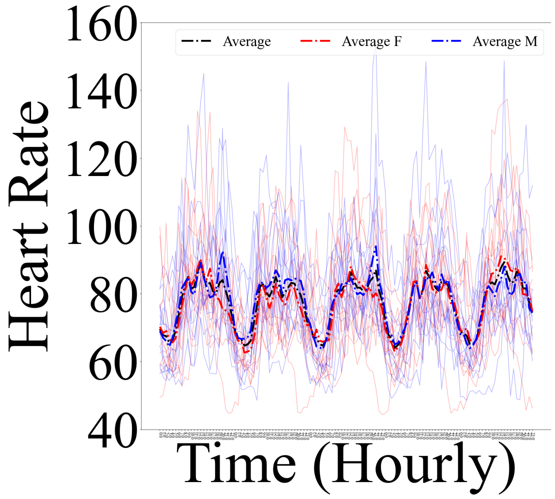

(a) HR

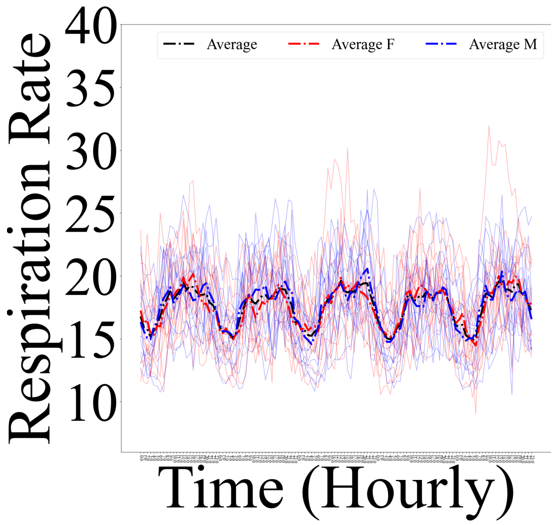

(b) RR

**Fig. B7**: (a) HR (b) RR time series of subset of participants who had only 5 days of their experiment available (total of 26 participants). In these subplots, HR and RR time series, per participant, are averaged in a 1-hour non-overlapping interval. Female and male participants are presented in solid red and blue lines. Dashed red and blue lines are grand average HR and RR for female and male participants. Dashed black line represent grand average of both female and male participants

##### B.6 6 Days of Experiment

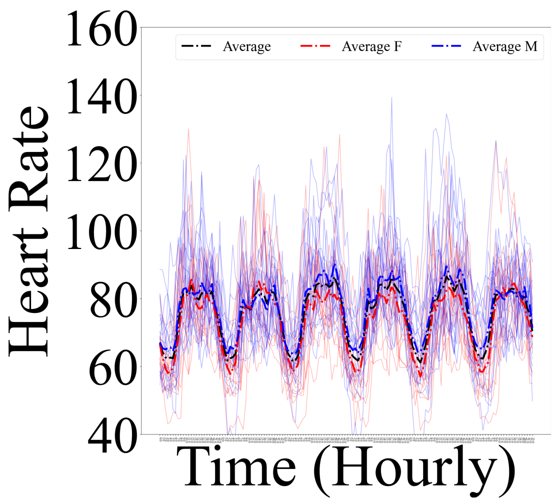

(a)

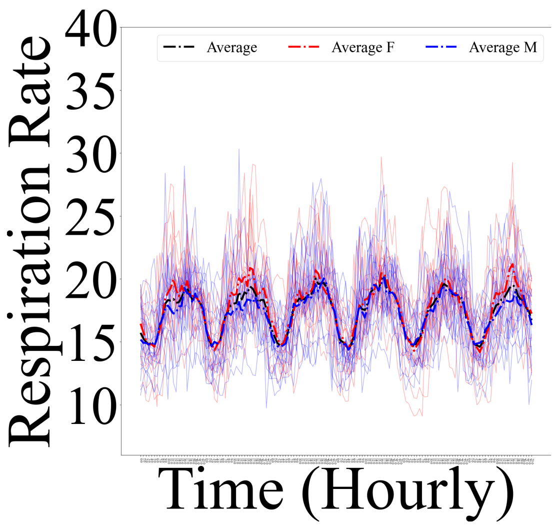

(b)

**Fig. B8**: (a) HR (b) RR time series for subset of participants who had all 6 days of their HR and RR recordings available (total of 33 participants). In these subplots, HR and RR time series, per participant, are averaged in a 1-hour non-overlapping interval. Female and male participants are presented in solid red and blue lines. Dashed red and blue lines are females' and males' grand averages. Dashed black line represents grand average of both female and male participants together.

### Appendix C HR — RR Trajectories

#### C.1 1 Day of Experiment

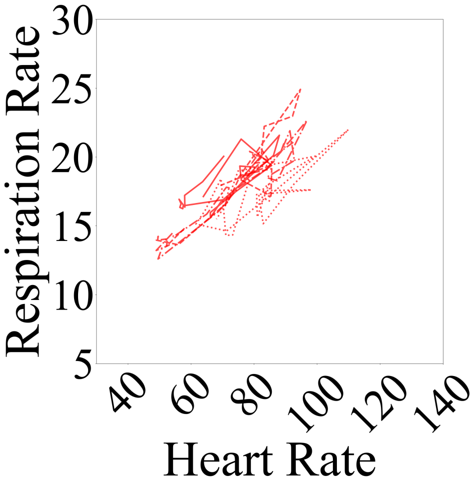

(a) Females

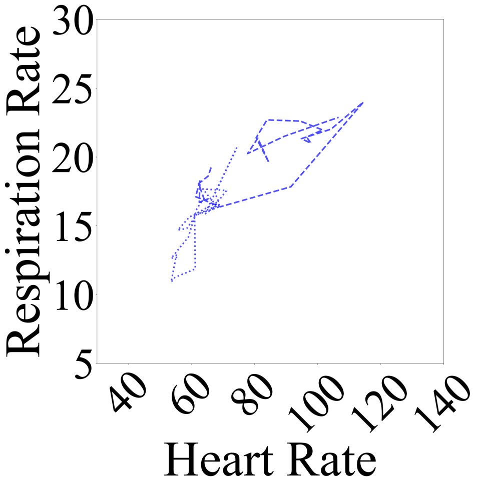

(b) Males

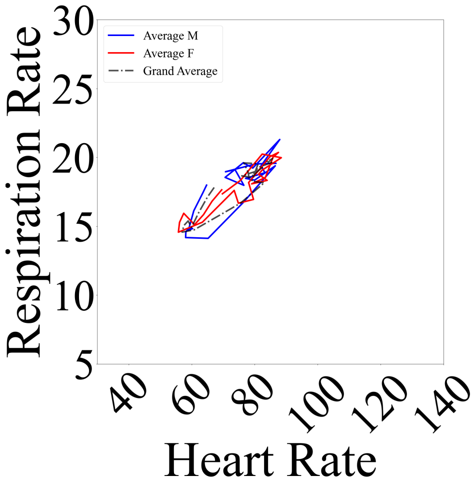

(c) Grand Averages

**Fig. C9**: HR — RR trajectories for 4 randomly selected (a) female (b) male participants with 1 day of their HR and RR time series available. (c) female (in red) and male (in blue) groups' averaged trajectories along with their grand average (in black).

##### C.2 2 Days of Experiment

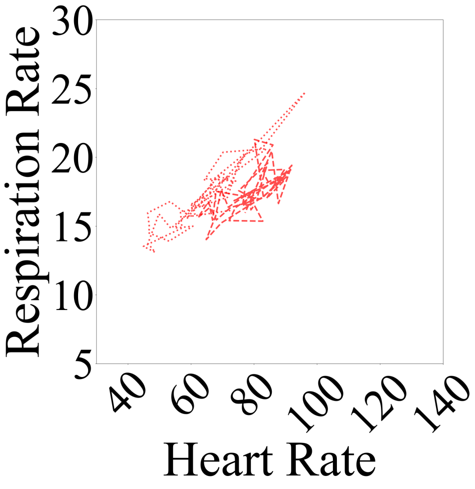

(a) Females

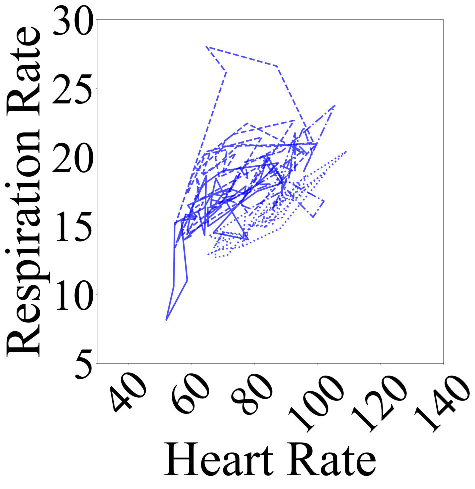

(b) Males

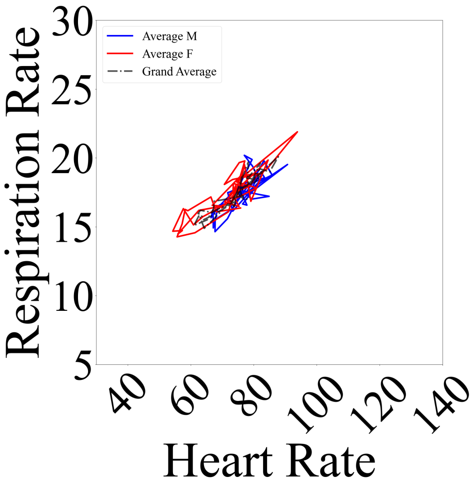

(c) Grand Averages

**Fig. C10**: HR — RR trajectories for 4 randomly selected (a) female (b) male participants with 2 days of their HR and RR time series available. (c) female (in red) and male (in blue) groups' averaged trajectories along with their grand average (in black).

#### C.3 3 Days of Experiment

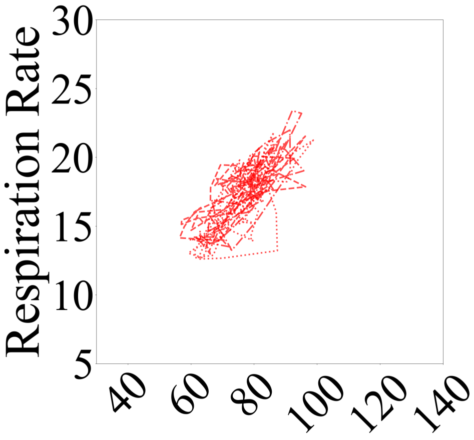

(a) Females

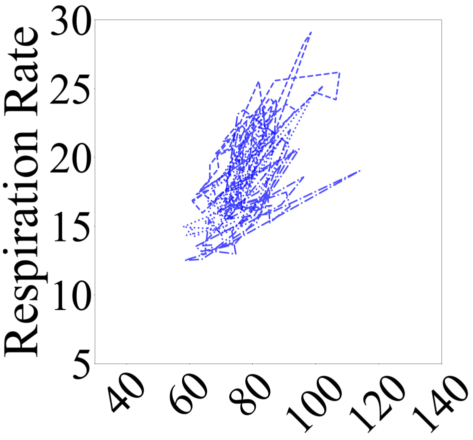

(b) Males

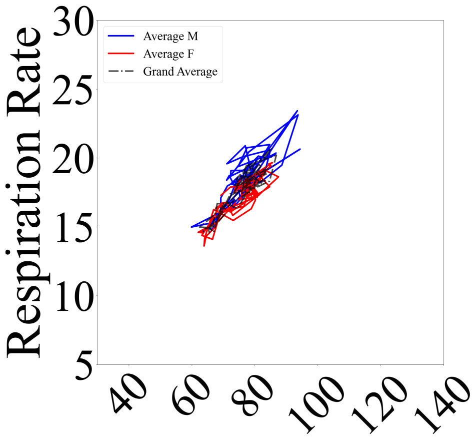

(c) Grand Averages

**Fig. C11**: HR — RR trajectories for 4 randomly selected (a) female (b) male participants with 3 days of their HR and RR time series available. (c) female (in red) and male (in blue) groups' averaged trajectories along with their grand average (in black).

#### C.4 4 Days of Experiment

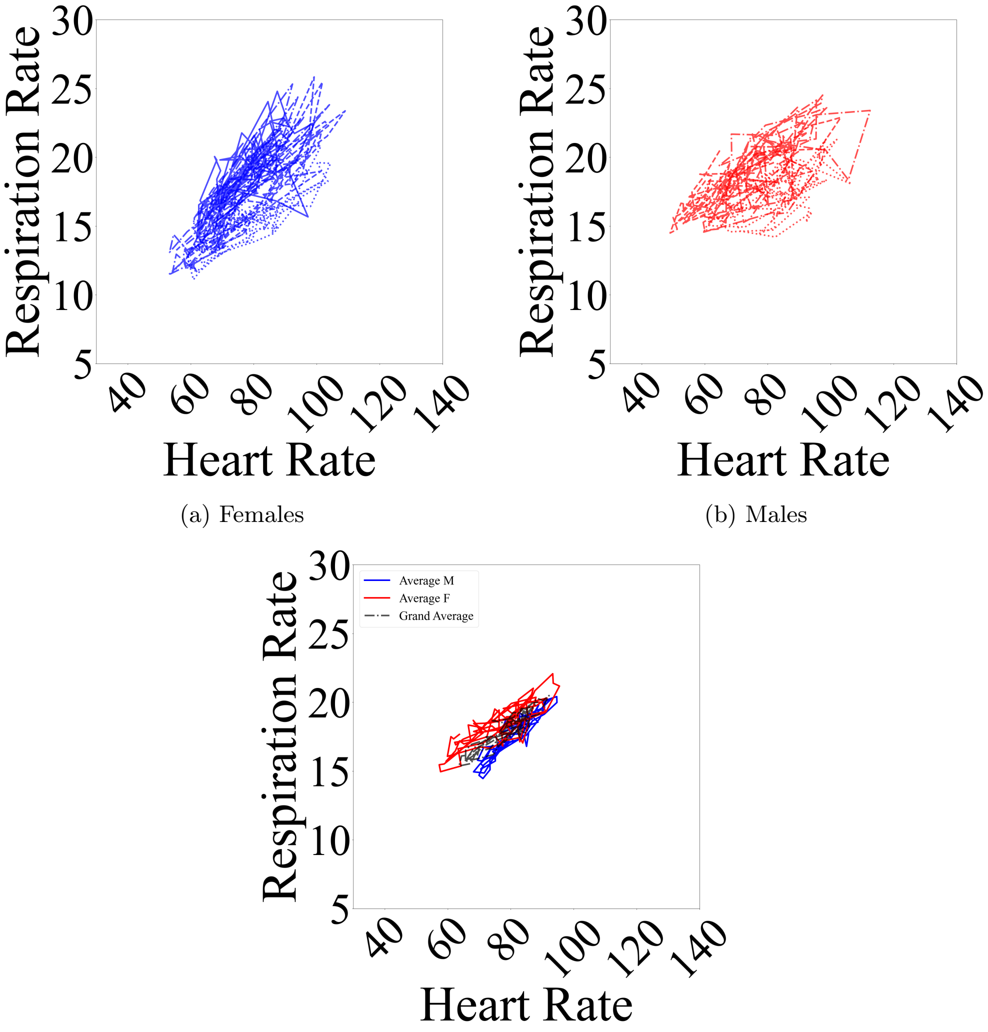

(a) Females

(b) Males

(c) Grand Averages

**Fig. C12**: HR — RR trajectories for 4 randomly selected (a) female (b) male participants with 4 days of their HR and RR time series available. (c) female (in red) and male (in blue) groups' averaged trajectories along with their grand average (in black).

#### C.5 5 Days of Experiment

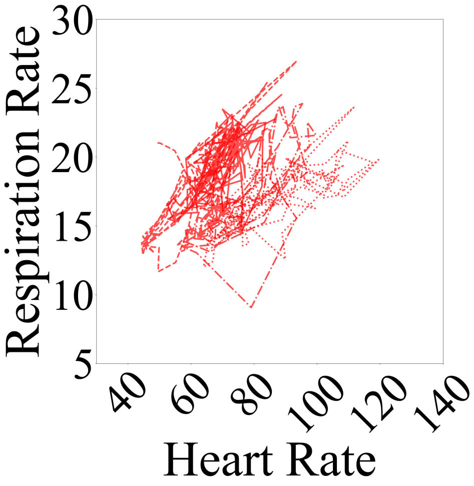

(a) Females

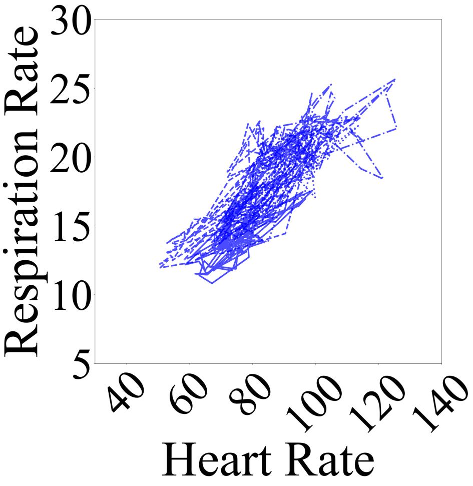

(b) Males

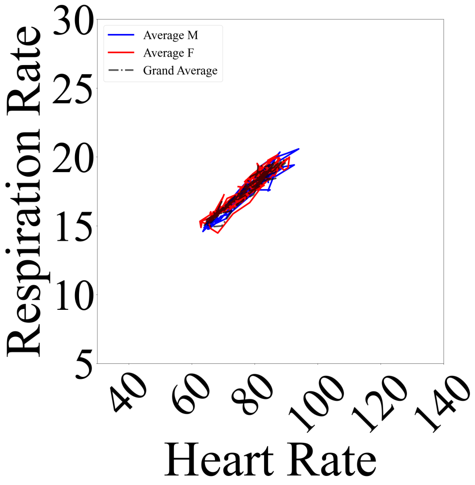

(c) Grand Averages

**Fig. C13**: HR — RR trajectories for 4 randomly selected (a) female (b) male participants with 5 days of their HR and RR time series available. (c) female (in red) and male (in blue) groups' averaged trajectories along with their grand average (in black).

#### C.6 6 Days of Experiment

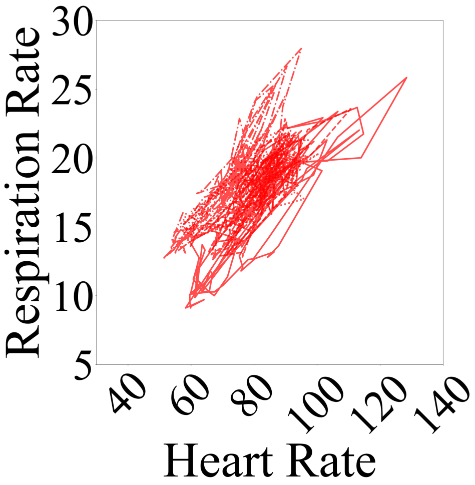

(a) Females

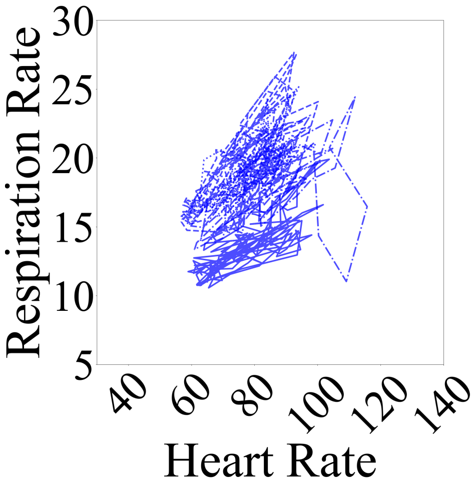

(b) Males

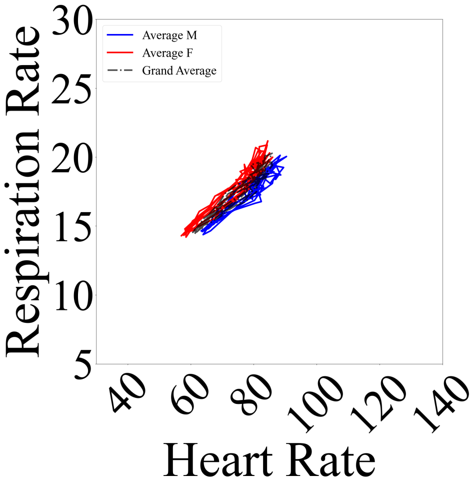

(c) Grand Averages

**Fig. C14**: HR — RR trajectories for 4 randomly selected (a) female (b) male participants with all 6 days of their HR and RR recordings available. (c) female (in red) and male (in blue) averaged trajectories along with their grand average (in black). See SM1 for sample HR — RR trajectories of participants who had 1 through 5 days of their data available.

### Appendix D HR — RR Correlation

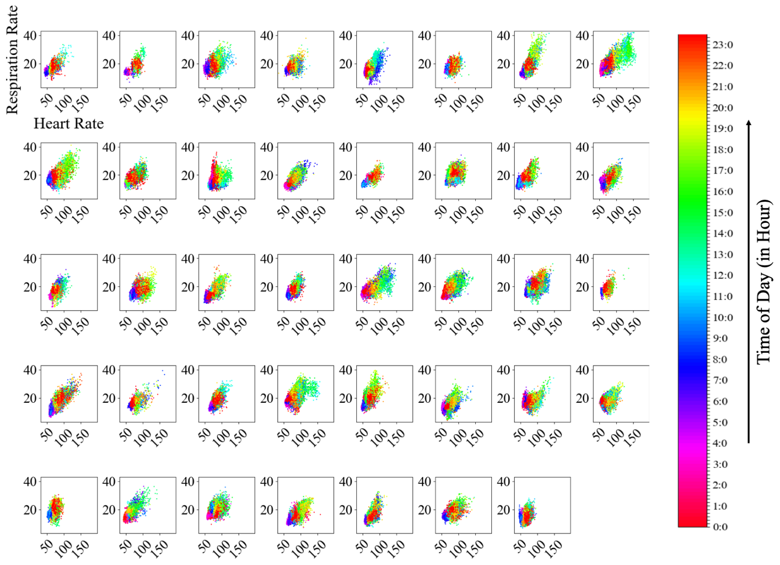

(a)

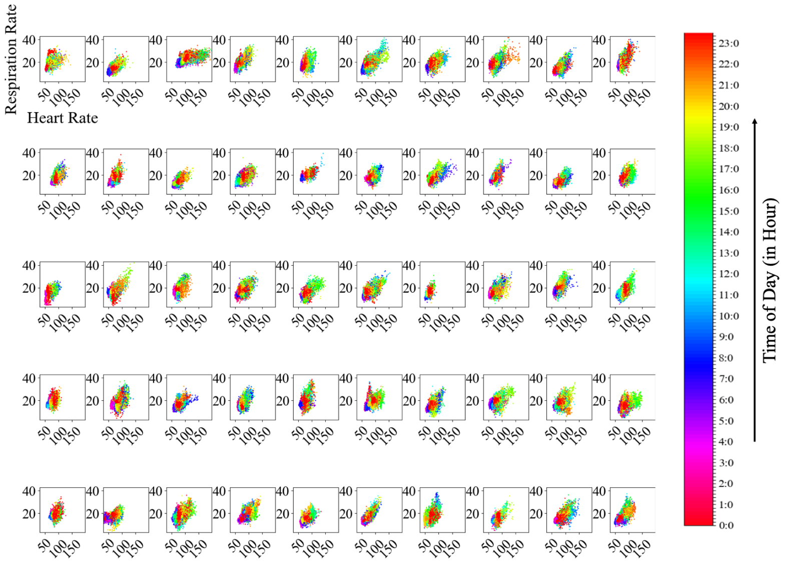

(b)

**Fig. D15**: HR — RR correlation of (a) females (b) males participants. Data points correspond to 1-minute non-overlapping moving average on HR and RR time series. They are color-coded as per time-of-day (in an hourly basis).

##### D.0.1 Participants' HR State-Space

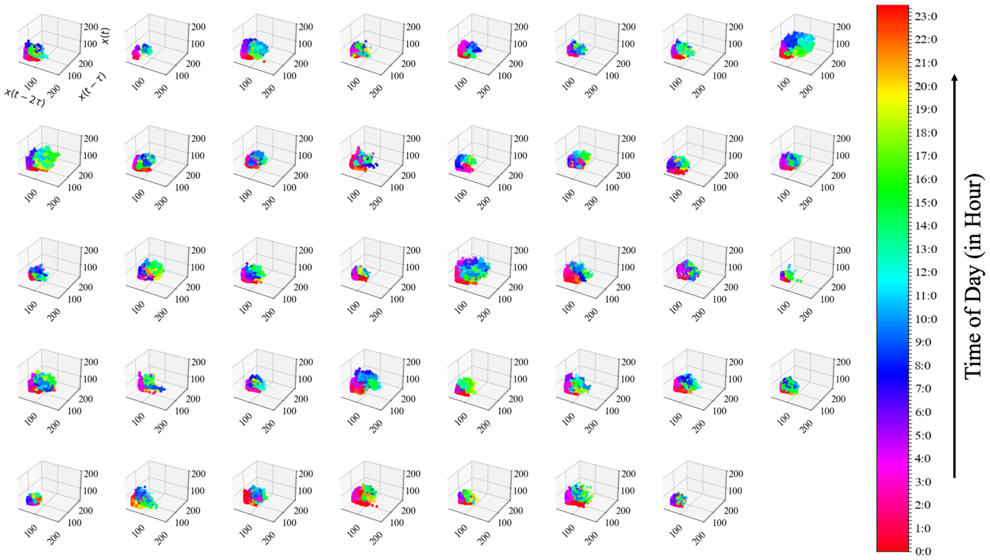

(a)

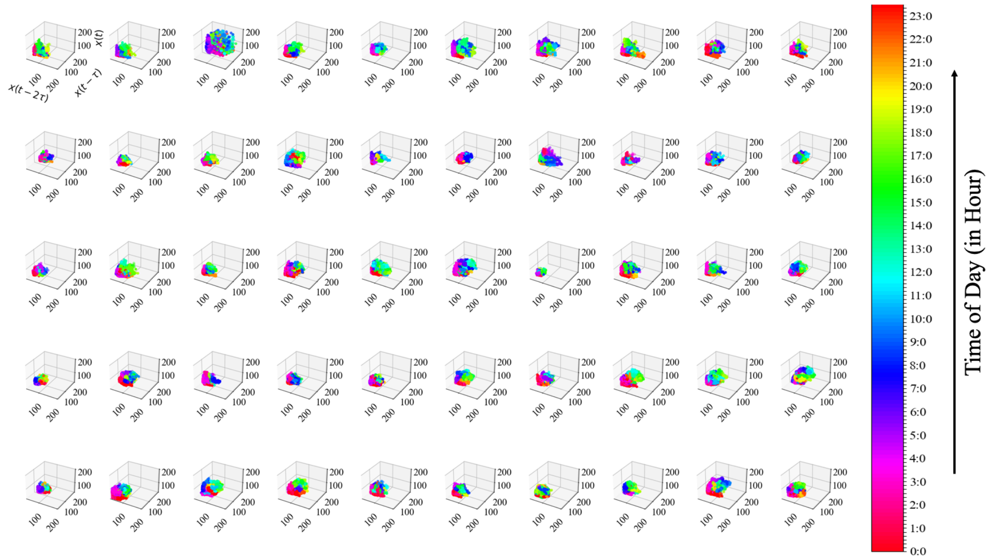

(b)

**Fig. D16**: HR state-space of (a) females (b) males participants, using delay embedding $\mu_\tau = 128$ i.e., 2 hours 8 minutes.

### Appendix E Exercise

Overall, older participants exercised significantly more than younger individuals in our sample (Figure E17, test-statistics = -5.00, p = 4.50$e^{-02}$, g = -0.4082, No Exercise: M = 56.5897, Mdn = 54.0, SD = 5.6601, $CI_{95\%}$ = [51.7532, 63.0993], Exercise: M = 59.0200, Mdn = 59.00, SD = 6.1725, $CI_{95\%}$ = [53.5215, 65.9967]).

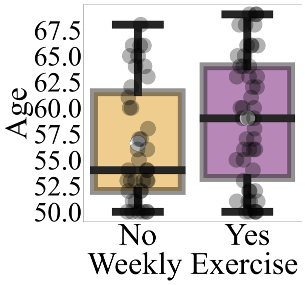

**Fig. E17**: Older participants exercised significantly more than younger individuals in our sample.

### Appendix F BMI- and HR-Related Gender Differences

Females and males differed significantly with respect to their Sleep HR (Figure F18a, test-statistics = -5.0733, p = 1.96$e^{-02}$, g = -0.5440, Females: M = 63.6169, Mdn = 62.7786, SD = 7.6828, $CI_{95\%}$ = [57.2283, 72.9174], Males: M = 67.8306, Mdn = 67.8519, SD = 7.7956, $CI_{95\%}$ = [61.2273, 77.3428]) and their BMI (Figure F18b, test-statistics = -2.4450, p = 3.20$e^{-03}$, g = -0.7579, Females: M = 21.4903, Mdn = 20.9572, SD = 3.0319, $CI_{95\%}$ = [19.1466, 25.658], Males: M = M = 24.0239, Mdn = 23.4022, SD = 3.5654, $CI_{95\%}$ = [21.0301, 28.4912]).

### Appendix G Effect of Weekly Alcohol Consumption on BMI and HR (12:00 – 18:00 PM)

Weekly alcohol consumption had significant effects on "Non-Drinkers"' vs. "Drinkers"' BMI and afternoon HR in that the latter had significantly higher BMI(Figure G19a, test-statistics = -2.0724, p = 1.42$e^{-02}$, g = -0.3676, Non-Drinkers: M = 22.1739, Mdn = 21.2510, SD = 3.7793, $CI_{95\%}$ = [19.4346, 27.4019], Drinkers: M = 23.4649, Mdn = 23.3234, SD = 3.3009, $CI_{95\%}$ = [20.6544, 27.3921]) and lower HR in the after noon (Figure G19b, test-statistics = 4.0527, p = 1.58$e^{-02}$, g = 0.5095, Non-Drinkers: M=85.3705, Mdn = 84.1349, SD = 9.1376, $CI_{95\%}$ = [77.9797, 97.1628], Drinkers: M = 81.0995, Mdn = 80.0822, SD = 7.7780, $CI_{95\%}$ = [74.4859, 90.3296]).

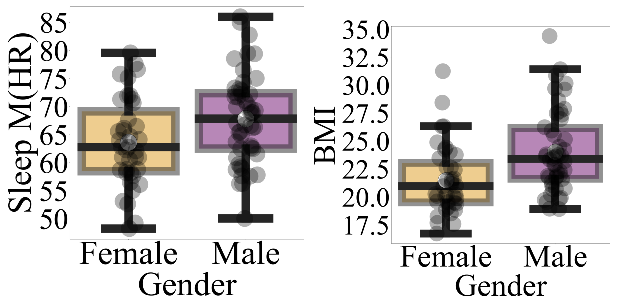

(a) (b)

**Fig. F18**: Gender differences in (a) Sleep M(HR) (b) BMI based on non-parametric bootstrap (10,000 rounds) permutation test of difference in two groups' Mdn.

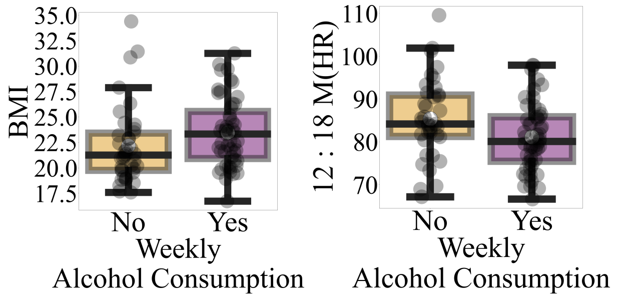

(a) (b)

**Fig. G19**: Effect of weekly alcohol consumption on individuals' (a) BMI (b) M(HR) (12:00 – 18:00 PM) based on non-parametric bootstrap (10,000 rounds) permutation test of difference in two groups' Mdn.

### Appendix H Relation Between Smoking And Participants' Age, BMI, Early Morning (6:00 – 7:00 AM) HR and RR, Overall and Wake RR

There were significantly more older than younger individuals with smoking habit [1] (Figure H20a, test-statistics = -7.00, p = 9.00e$^{-03}$, g = -0.6912, Non-Smokers: M = 56.7419, Mdn = 56.00, SD = 5.5152, $CI_{95\%}$ = [51.8753, 63.0173], Smokers: M = 60.7407, Mdn = 63.00, SD = 6.3747, $CI_{95\%}$ = [55.3382, 68.1498]).

"Smokers" had significantly higher BMI (Figure H20b, test-statistics = -1.9897, p = 3.64e$^{-02}$, g = -0.6330, Non-Smokers: M = 22.2558, Mdn = 21.6431, SD = 3.2253,

[1] "Smokers" included those who quit smoking. This choice was due to the limited number of smokers in our sample (6 smokers in total, See main manuscript, Table 2.

$CI_{95\%}$ = [19.5396, 26.1829], Smokers: M = 24.4244, Mdn = 23.6328, SD = 3.8554, $CI_{95\%}$ = [21.4239, 29.4868]), early morning (6:00 – 7: 00 AM) HR (Figure H20c, test-statistics = -5.6840, p = 3.44e$^{-02}$, g = -0.5504, Non-Smokers: M = 67.0160, Mdn = 66.2926, SD = 10.5300, $CI_{95\%}$ = [57.9503, 79.5833], Smokers: M = 72.8926, Mdn = 71.9767, SD = 11.0155, $CI_{95\%}$ = [64.1545, 87.4602]), early morning (6:00 – 7: 00 AM) RR (Figure H20d, test-statistics = -1.5181, p = 7.20e$^{-03}$, g = -0.6292, Non-Smokers: M = 15.4820, Mdn = 15.3252, SD = 2.0744, $CI_{95\%}$ = [13.7169, 17.9443], Smokers: M = 16.8918, Mdn = 16.8433, SD = 2.5891, $CI_{95\%}$ = [14.8374, 20.2956]), Overall RR (Figure H20e, test-statistics = -0.8672, p = 4.86e$^{-02}$, g = -0.4020, Non-Smokers: M = 16.7734, Mdn = 16.8228, SD = 1.439, $CI_{95\%}$ = [15.5438, 18.4676, Smokers: M = 17.3906, Mdn = 17.6900, SD = 1.7412, $CI_{95\%}$ = [16.03, 19.7258]), and Wake RR (Figure H20f, test-statistics = -0.7920, p = 4.24e$^{-02}$, g = -0.3917, Non-Smokers: M = 18.1714, Mdn = 18.1510, SD = 1.6502, $CI_{95\%}$ = [16.7713, 20.1408], Smokers: M = 18.8286, Mdn = 18.9430, SD = 1.7400, $CI_{95\%}$ = [17.5398, 21.2323]).

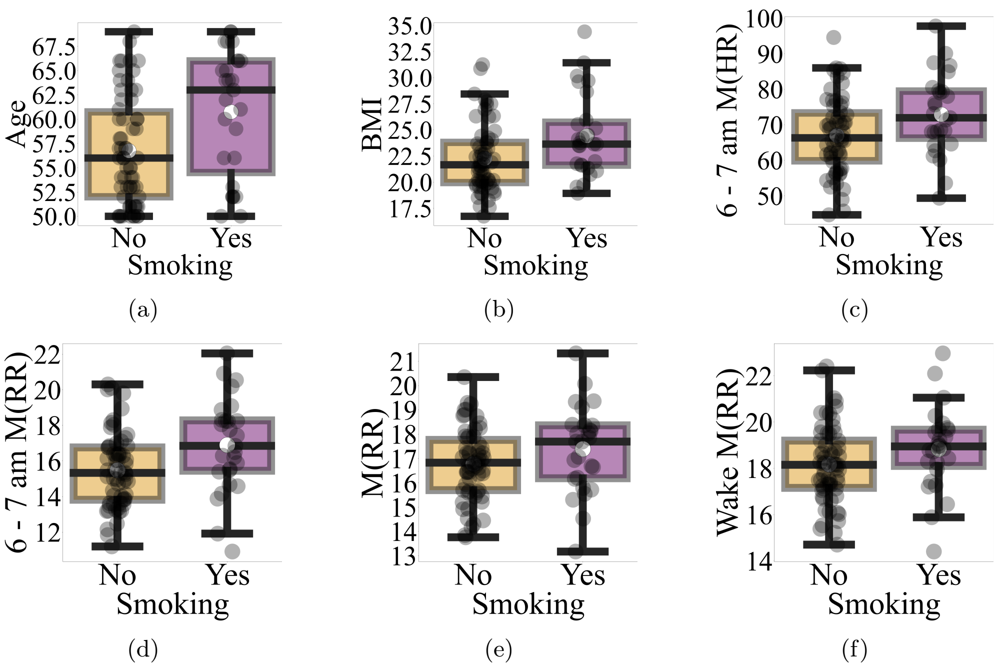

**Fig. H20**: Relation between smoking and participants' (a) Age (b) BMI (c) Early Morning (6:00 – 7:00 AM) HR (d) Early Morning (6:00 – 7:00 AM) RR (e) Overall RR (e) Wake RR based on non-parametric bootstrap (10,000 rounds) permutation test of difference in two groups' Mdn.
