## Supplementary Materials 3 for "Information Dynamics of the Heart and Respiration Rates: a Novel Venue for Digital Phenotyping in Humans"

### Supplementary Materials 3 (SM3):
Respiratory Modulation of the Heart Rate Signifies the Effect of Alcohol Consumption on Cardiorespiratory Function in Human

Soheil Keshmiri[1*], Sutashu Tomonaga[2], Haruo Mizutani[3], Kenji Doya[2]

[1]Optical Neuroimaging Unit, Okinawa Institute of Science and Technology, Okinawa, Japan.
[2]Neural Computation Unit (NCU), Okinawa Institute of Science and Technology, Okinawa, Japan.
[3]Suntory Global Innovation Center Limited (SGIC), Suntory, Kyoto, Japan.

Contributing authors:;
;

### Appendix A HR Moments

#### A.1 AR $w_1$ — $w_2$ Plane

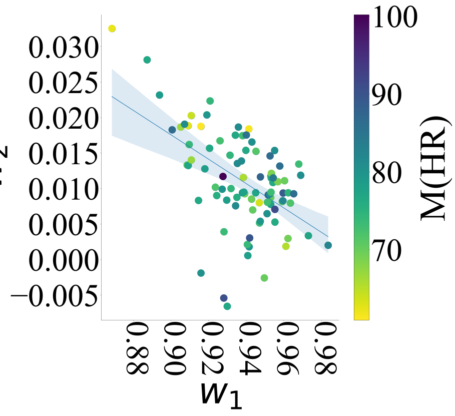

(a)

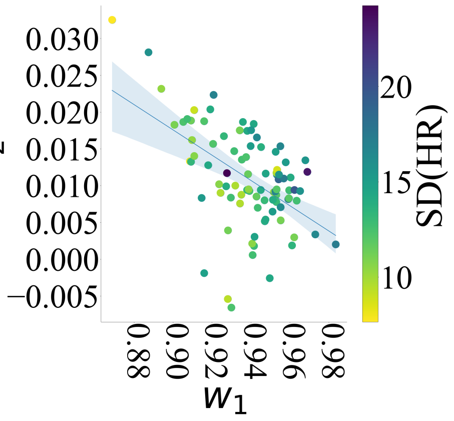

(b)

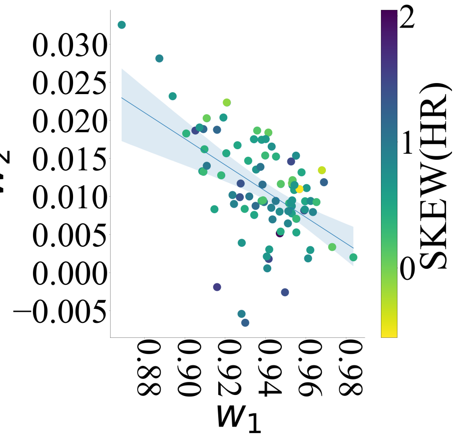

(c) )

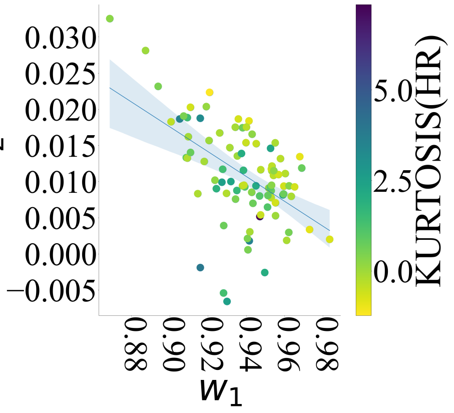

(d)

**Fig. A1**: HR Moments within AR $w_1$ — $w_2$ plane (a) Mean (M) (b) Standard Deviation (SD) (c) Skewness (S) (d) Kurtosis (K). None of HR's moments maintained observed patterns related to AR accuracy $R^2$ and HR state-space information dynamics.

#### A.2 AR $w_1$ — $w_3$ Plane

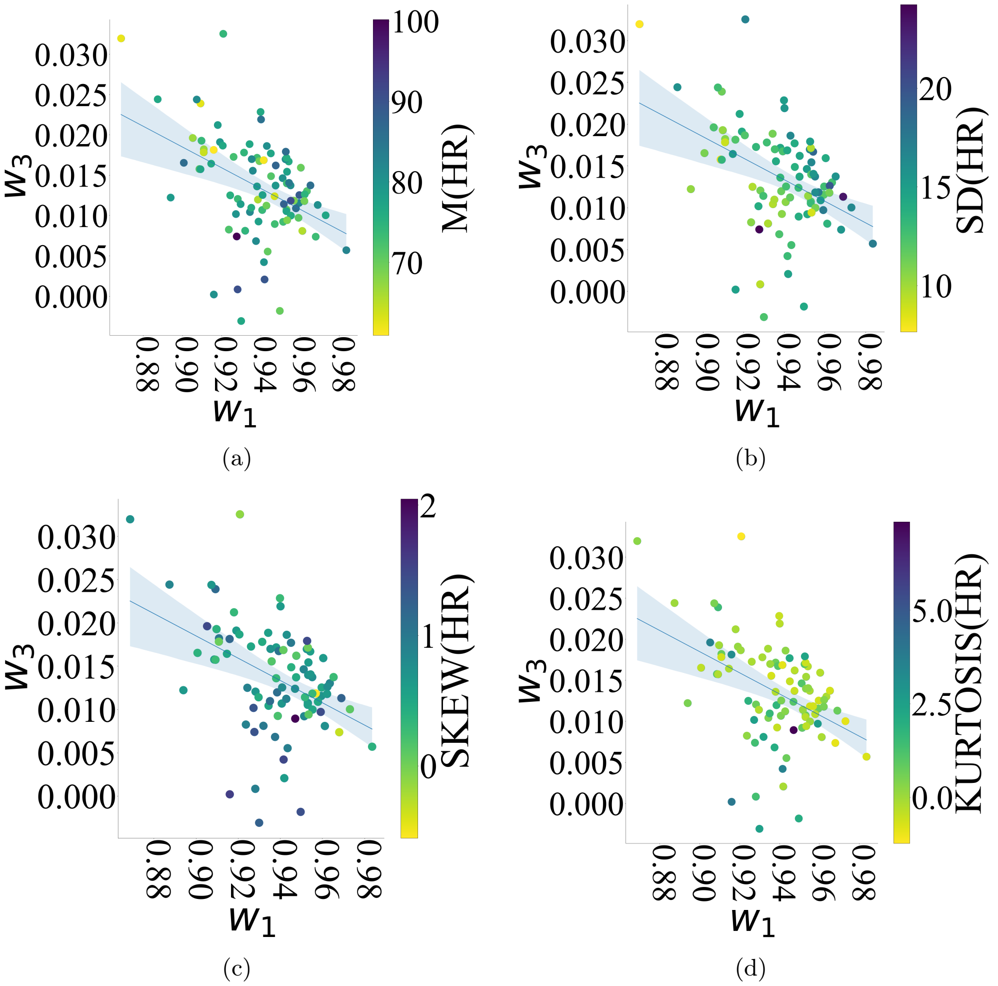

**Fig. A2**: HR Moments within AR $w_1$ — $w_3$ plane (a) Mean (M) (b) Standard Deviation (SD) (c) Skewness (S) (d) Kurtosis (K). None of HR's moments maintained observed patterns related to AR accuracy $R^2$ and HR state-space information dynamics.

### Appendix B HR Powers

#### B.1 AR $w_1$ — $w_2$ Plane

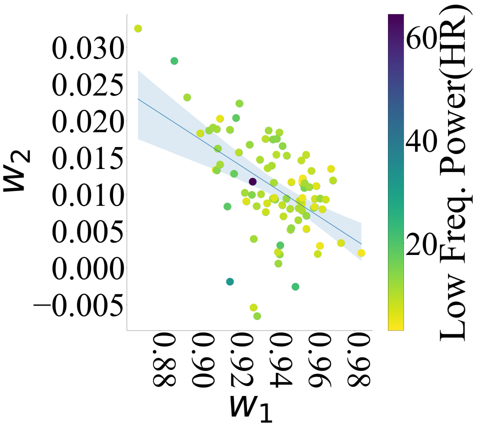

(a)

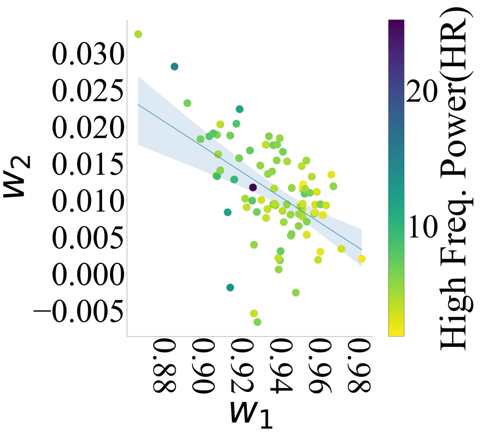

(b)

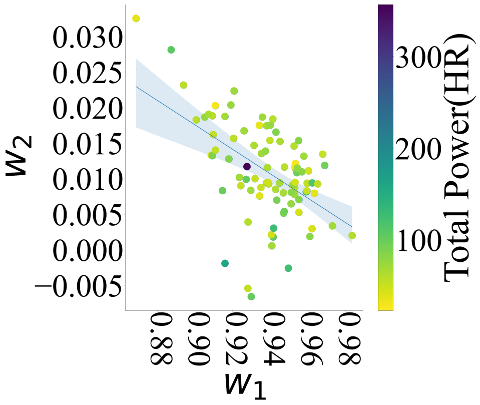

(c)

**Fig. B3**: HR Powers within AR $w_1$ — $w_2$ plane (a) Low Frequency (LF) (M) (b) High Frequency (HF) (c) Total Power (TP). None of HR's (frequency) powers maintained observed patterns related to AR accuracy $R^2$ and HR state-space information dynamics.

##### B.2 AR $w_1$ — $w_3$ Plane

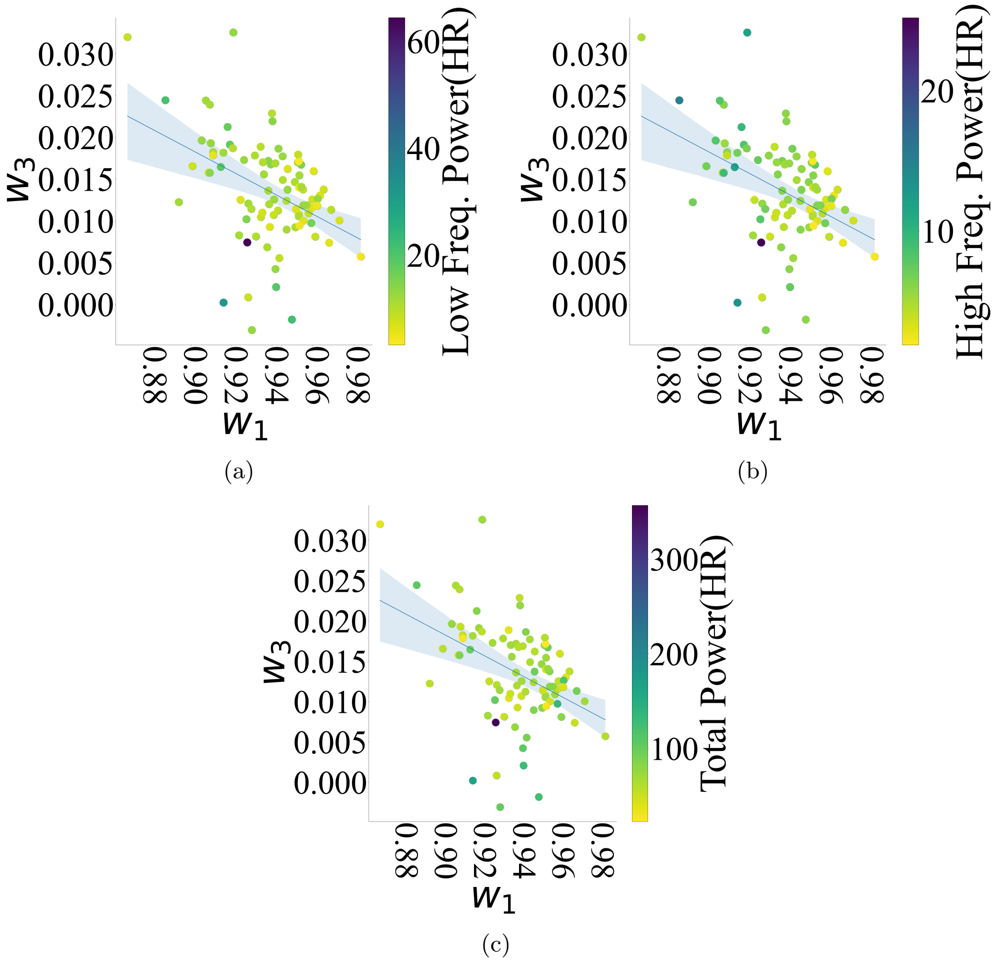

**Fig. B4**: HR Powers within AR $w_1$ — $w_3$ plane (a) Low Frequency (LF) (M) (b) High Frequency (HF) (c) Total Power (TP). None of HR's (frequency) powers maintained observed patterns related to AR accuracy $R^2$ and HR state-space information dynamics.

### Appendix C HR Poincare SD1 and SD2

#### C.1 AR $w_1$ — $w_2$ Plane

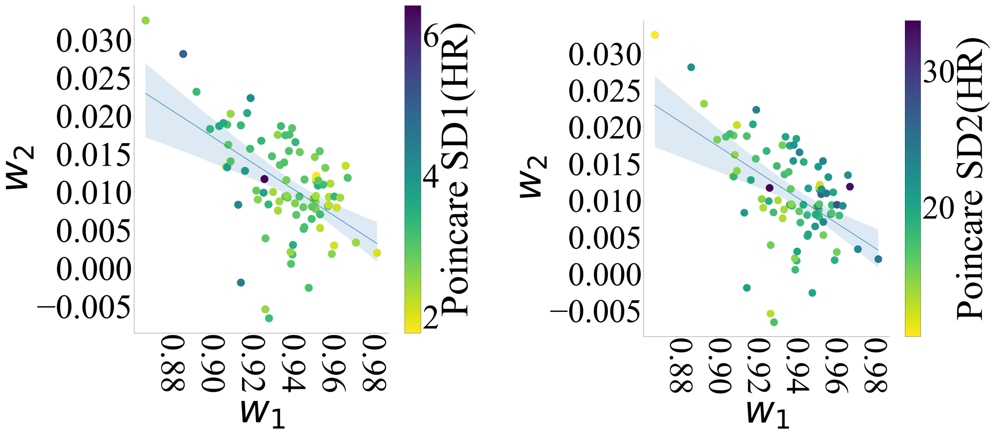

(a) (b)

**Fig. C5**: HR Poincare SD1 and SD2 within AR $w_1$ — $w_2$ plane (a) Poincare SD1 (SD1) (LF) (b) Poincare SD2 (SD2). None of HR's (Poincare) SD1 (minor axis, short-term variability) and SD2 (major axis, long-term variability) maintained observed patterns related to AR accuracy $R^2$ and HR state-space information dynamics.

##### C.2 AR $w_1$ — $w_3$ Plane

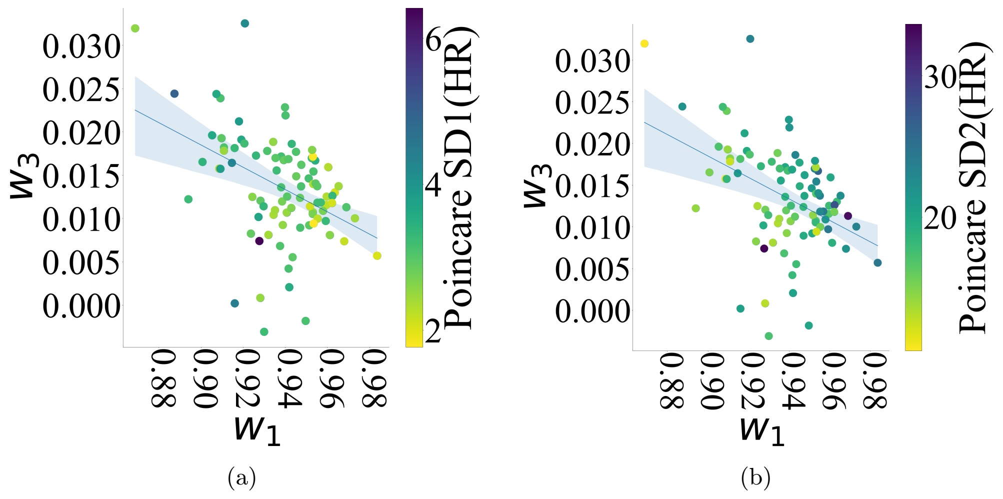

**Fig. C6**: HR Poincare SD1 and SD2 within AR $w_1$ — $w_3$ plane (a) Poincare SD1 (SD1) (LF) (b) Poincare SD2 (SD2). None of HR's (Poincare) SD1 (minor axis, short-term variability) and SD2 (major axis, long-term variability) maintained observed patterns related to AR accuracy $R^2$ and HR state-space information dynamics.

### Appendix D Other Measures Within $HR \rightarrow RR$ — $RR \rightarrow HR$ Plane

#### D.1 Moments

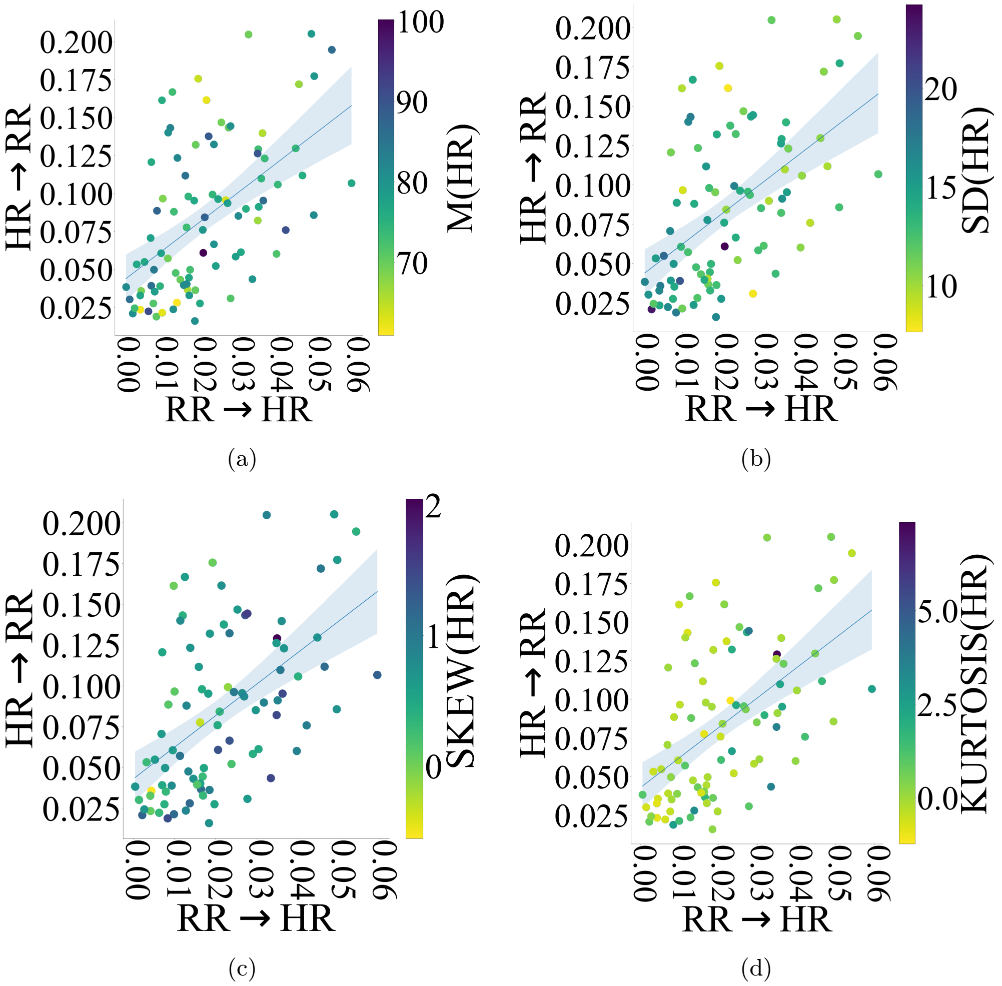

**Fig. D7**: HR Moments within $HR \rightarrow RR$ — $RR \rightarrow HR$ plane (a) Mean (M) (b) Standard Deviation (SD) (c) Skewness (S) (d) Kurtosis (K). None of HR's moments maintained observed patterns related to AR accuracy $R^2$ and HR state-space information dynamics.

#### D.2 HR Powers

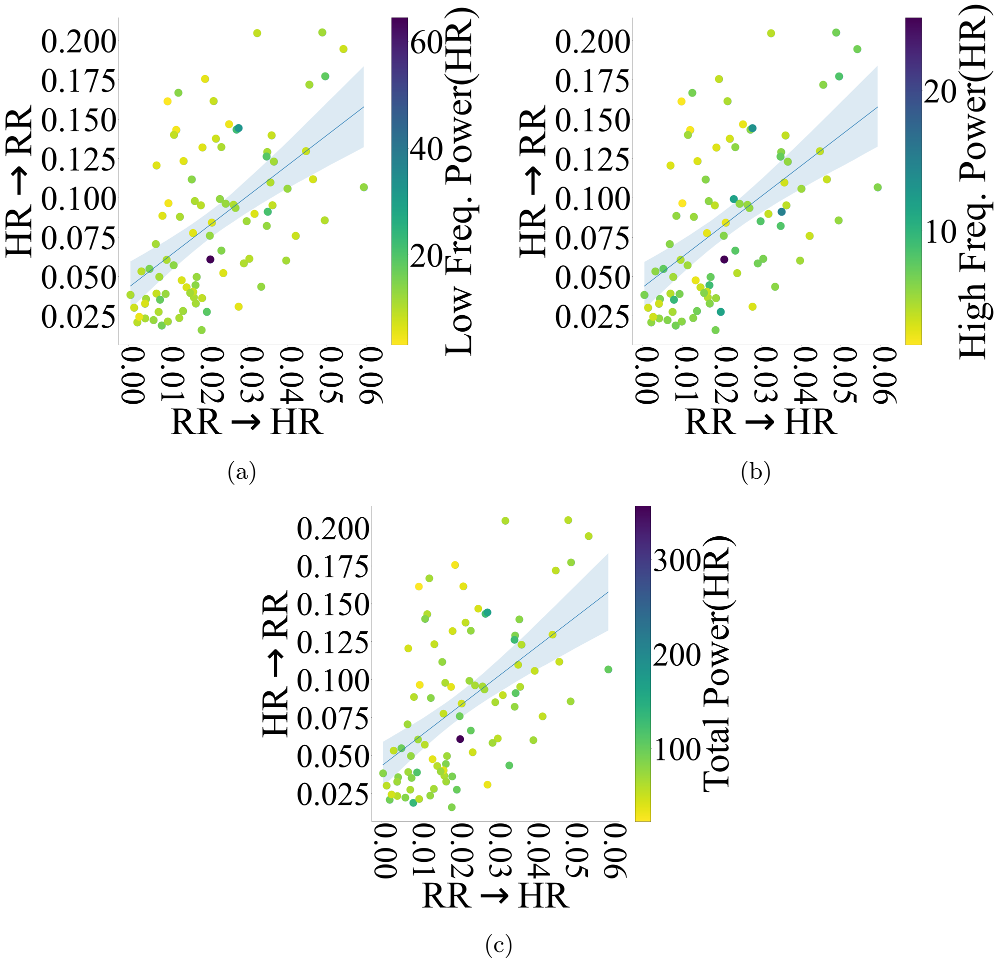

**Fig. D8**: HR Powers within within $HR \rightarrow RR$ — $RR \rightarrow HR$ plane (a) Low Frequency (LF) (M) (b) High Frequency (HF) (c) Total Power (TP). None of HR's (frequency) powers maintained observed patterns related to AR accuracy $R^2$ and HR state-space information dynamics.

#### D.3 Poincare SD1 and SD2

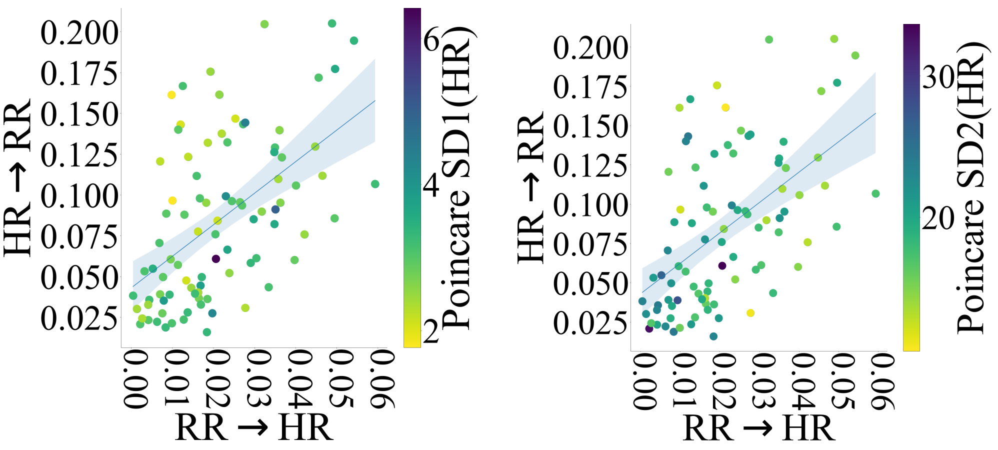

(a) (b)

**Fig. D9**: HR Poincare SD1 and SD2 within $HR \rightarrow RR$ — $RR \rightarrow HR$ plane (a) Poincare SD1 (SD1) (LF) (b) Poincare SD2 (SD2). None of HR's (Poincare) SD1 (minor axis, short-term variability) and SD2 (major axis, long-term variability) maintained observed patterns related to AR accuracy $R^2$ and HR state-space information dynamics.

### Appendix E Other Measures Within $RR \rightarrow HR$ — $R^2$ Plane

#### E.1 Moments

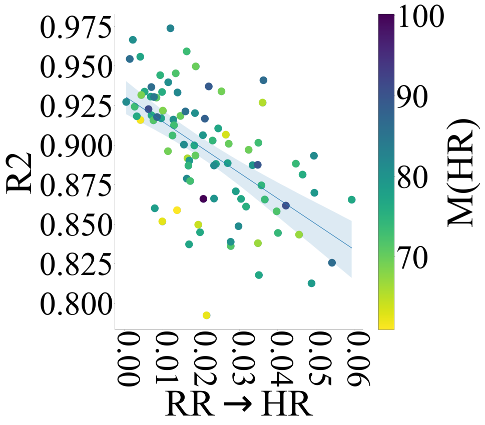

(a)

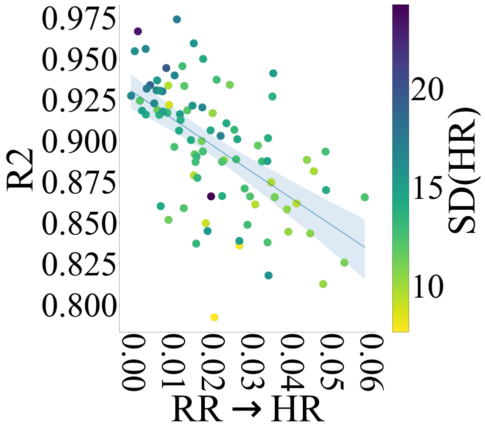

(b)

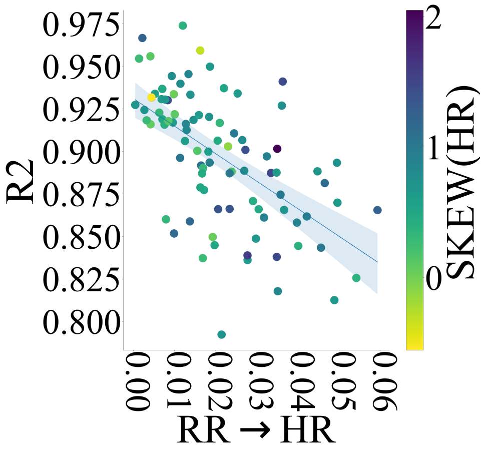

(c)

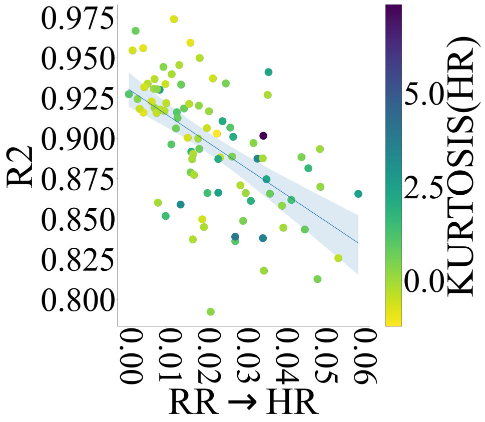

(d)

**Fig. E10**: $RR \rightarrow HR$ and AR accuracy $R^2$ plane with respect to HR's moments (a) Mean (M) (b) Standard Deviation (SD) (c) Skewness (S) (d) Kurtosis (K). None of HR's moments maintained observed patterns related to AR accuracy $R^2$ and HR state-space information dynamics.

#### E.2 HR Powers

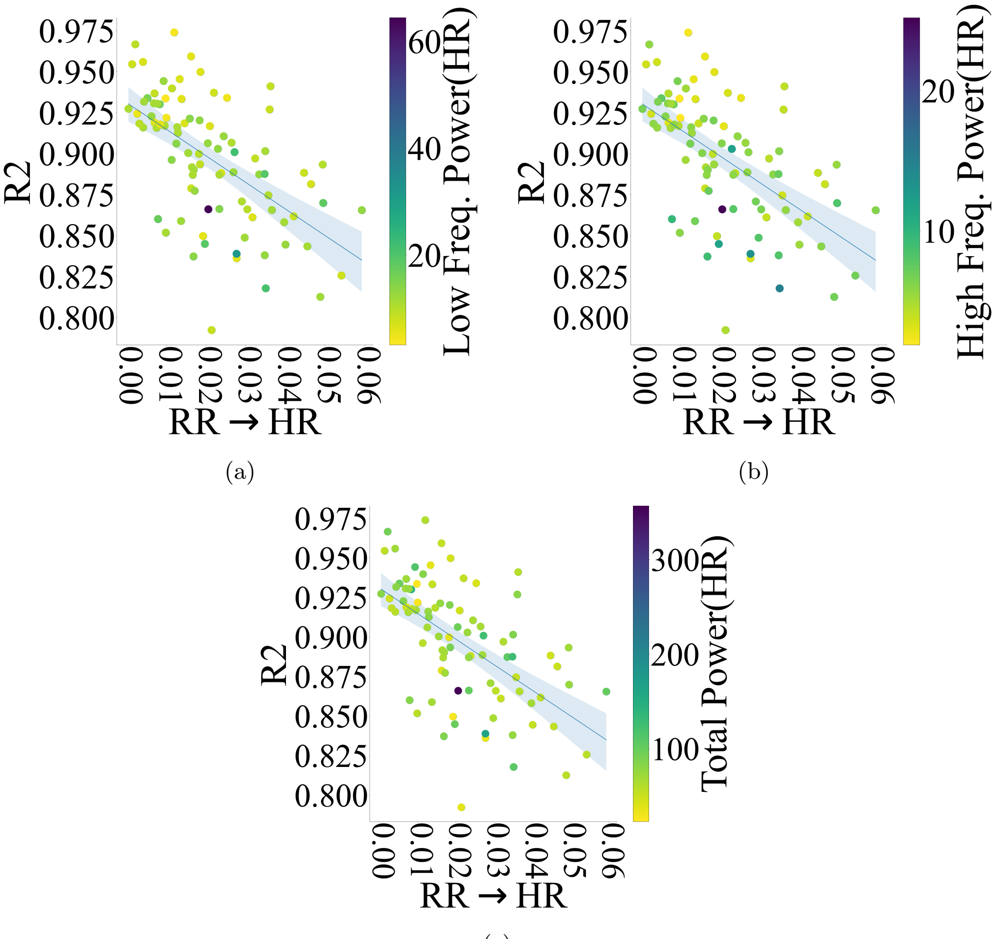

(a) (b) (c)

**Fig. E11**: $RR \rightarrow HR$ and AR accuracy $R^2$ plane with respect to HR Powers (a) Low Frequency (LF) (b) High Frequency (HF) (c) Total Power (TP). None of HR's (frequency) powers maintained observed patterns related to AR accuracy $R^2$ and HR state-space information dynamics.

##### E.3 Poincare SD1 and SD2

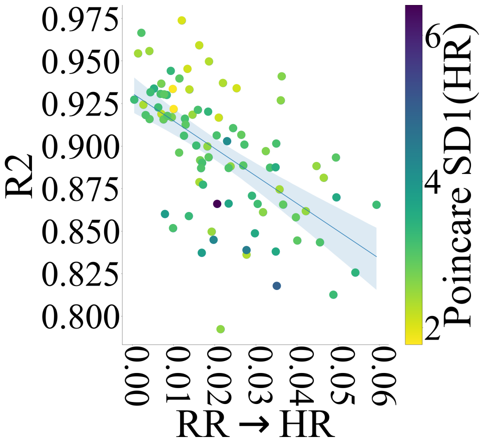

(a)

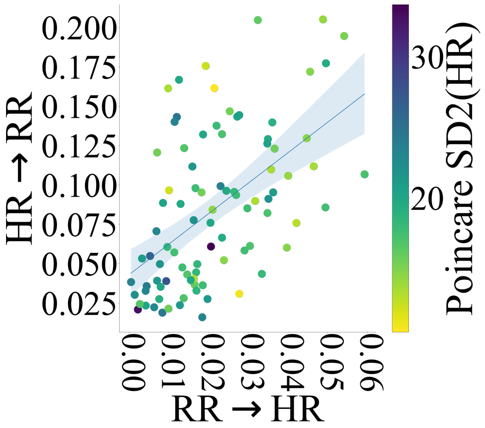

(b)

**Fig. E12**: $RR \to HR$ and AR accuracy $R^2$ plane with respect to Poincare (a) SD1 (SD1) (LF) (b) SD2 (SD2). None of HR's (Poincare) SD1 (minor axis, short-term variability) and SD2 (major axis, long-term variability) maintained observed patterns related to AR accuracy $R^2$ and HR state-space information dynamics.
